# Multiplex profiling of developmental enhancers with quantitative, single-cell expression reporters

**DOI:** 10.1101/2022.12.10.519236

**Authors:** Jean-Benoît Lalanne, Samuel G. Regalado, Silvia Domcke, Diego Calderon, Beth Martin, Tony Li, Chase C. Suiter, Choli Lee, Cole Trapnell, Jay Shendure

**Affiliations:** Department of Genome Sciences, University of Washington, Seattle, WA, USA; Howard Hughes Medical Institute, Seattle, WA, USA; Brotman Baty Institute for Precision Medicine, Seattle, WA, USA; Allen Discovery Center for Cell Lineage Tracing, Seattle, WA, USA

## Abstract

The inability to scalably and precisely measure the activity of developmental enhancers in multicellular systems is a bottleneck in genomics. Here, we develop a dual RNA cassette that decouples the detection and quantification tasks inherent to multiplex single-cell reporter assays, resulting in accurate measurement of reporter expression over a >10,000-fold range of activity with a precision approaching the limit set by Poisson counting noise. Together with RNA barcode circularization, these <u>s</u>ingle-<u>c</u>ell <u>q</u>uantitative <u>e</u>xpression <u>r</u>eporters (scQers) provide high-contrast readouts analogous to classic *in situ* assays, but entirely from sequencing. Screening >200 enhancers in a multicellular *in vitro* model of early mammalian development, we identified numerous autonomous and cell-type-specific elements, including constituents of the *Sox2* control region exclusively active in pluripotent cells, endoderm-specific enhancers, including near *Foxa2* and *Gata4*, and a compact pleiotropic enhancer at the *Lamc1* locus. scQers can be mobilized in developmental systems to quantitatively characterize native, perturbed, and synthetic enhancers at scale, with high sensitivity and at single-cell resolution.

## Main Text

Developmental enhancers direct programs of gene expression that unfold with remarkable cell type and spatiotemporal specificity. This tight control underlies the robust emergence of form and function from a one-cell zygote. Fine regulatory changes of target genes, caused by even single nucleotide changes to individual enhancers, can both give rise to disease (*1*–*3*) as well as drive novelty across evolution (*1, 4*). Genetic methods have identified an extensive list of developmentally important genes in model systems (*5, 6*), yet how the transcription of these genes is regulated by enhancers, and specifically how DNA sequence encodes the requisite functional information, remains incompletely understood even for the best-studied examples (*7*–*10*). More broadly, biochemical marks correlated with enhancer status have now nominated over one million putative cis-regulatory elements (CREs) in the mouse and human genomes (*11*). However, functional profiling of these elements (and variants thereof) across diverse cellular states, particularly in developmental contexts, is lagging due to the lack of scalable approaches.

In mammalian systems, most high-throughput functional studies of CREs have been performed in static contexts, typically cancer cell lines (*12*–*15*). The scalability of these biotypes, in conjunction with massively parallel reporter assays (MPRAs) (*16*–*18*) and related techniques (*19*), have enabled the functional characterization of complex CRE libraries leading to accurate sequence-to-function models (*13, 20*). Extending beyond the unidimensional activity in cell lines, the function of CREs throughout development is inherently about specificity, namely the multidimensional cell-type to cell-type differences in function across *trans*-environments, the study of which requires new experimental and modeling approaches. Reporter assays have been applied to mammalian differentiation models (e.g., neuronal (*21*), naive to epiblast (*22*)), but these remain essentially simple trajectories. Single-cell chromatin accessibility data from systems containing extensive cell type heterogeneity can be used to train models predicting differential accessibility from DNA sequence (*23*–*25*), with promise to also correlatively predict cell-type-specific expression (*26*). However, these models remain one step removed from the functional outcome and are inherently limited given that differentially accessible genomic regions commonly lack autonomous expression-enhancing activity (*27*).

Until now, work on enhancers in multicellular systems has predominantly been carried out with transgenic reporters assayed via *in situs (28–30)*, approaches which remain semi-quantitative and of limited throughput even with automation (*31*). Nonetheless, even at limited scales, these studies reveal the rich phenomenology of metazoan developmental enhancers, namely that kilobase-sized elements can autonomously recapitulate the complex expression patterns of their target genes even when taken out of context. However, particularly as applied in mammalian models such as the mouse, these assays do not afford the scale or turnaround times required for “perturb-test-learn” loops necessary to construct mechanistic sequence-to-function maps. Compendia documenting enhancer activity in development exist (*30, 32*), but moving from catalogs to principles remains a challenge.

Two recent innovations are poised to improve the throughput of mammalian developmental enhancer biology. <u>First</u>, stem-cell-derived models of increasing complexity and fidelity to *in vivo* development, including organoids, gastruloids, and synthetic embryoids (*33*), enable the scalable delivery of genomically integrating reporters (*34*) prior to differentiation. <u>Second</u>, single-cell genomics can finely map cellular states and in principle be combined with multiplex reporter assays to increase the throughput at which enhancers are profiled in multicellular models (**Fig. 1A**). However, in practice, multiplex reporter measurements in a single-cell context pose a fundamentally new challenge compared to bulk modalities: in order to measure the activity of any given candidate CRE, one must first determine <u>which</u> reporters are present in <u>which</u> profiled cells. As such, porting the one-RNA reporter strategy of bulk MPRAs directly to single-cell platforms (**Fig. 1B**), one relies on the barcoded mRNA for both: 1) per-cell reporter detection; and 2) quantification of expression driven by the candidate CRE. The detection task is challenging for lowly expressed reporter transcripts due to chimeric amplicons (i.e., spurious amplification products erroneously swapping barcodes originally from different molecules), which increase the noise floor of sequencing counts in single-cell libraries (*35, 36*). To put it another way, in the simplest adaptation of MPRAs to the single cell context, one cannot distinguish between cells in which a given reporter is <u>not expressed</u> vs. cells in which a given reporter is <u>not present</u> (**Fig. 1B**). This inherently confounds the accurate quantification of enhancer activity.

**Figure 1:**
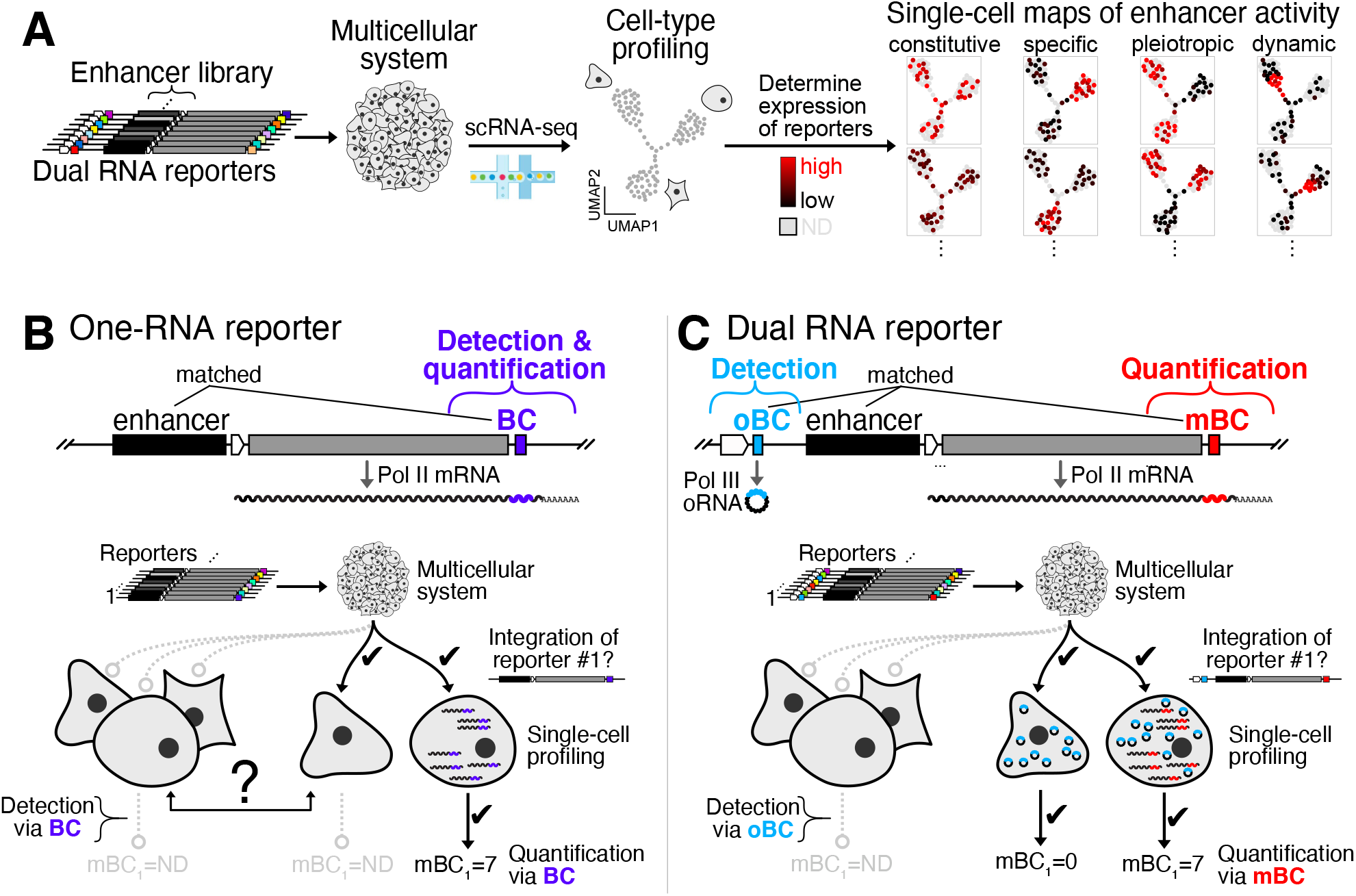
High-contrast single-cell enhancer activity maps with <u>s</u>ingle-<u>c</u>ell <u>q</u>uantitative <u>e</u>xpression <u>r</u>eporters (scQers) (**A**) Multiplex single-cell reporter assays. Introduction of complex libraries of integrating reporters to multicellular systems followed by scRNA-seq and computational deconvolution of reporter expression. (**B**) Traditional multiplex reporters harbor a single barcoded Pol II mRNA (BC, purple) driven by a library of enhancers whose activity is to be profiled. In a multiplex single-cell assay, having a single transcript to both detect presence of any given reporter in a profiled cell and measure expression level is biased. In the limiting case where no mRNA is produced from an enhancer in a given cell type, direct detection of the reporter is not possible (left group vs. middle cell). (**C**) To resolve this dropout problem, a constitutively and highly expressed Pol III-derived circularized barcoded RNA (*37*) (Tornad<u>o b</u>ar<u>c</u>odes, oBC, blue), *a priori* matched with the mBC (red) and enhancer, is appended co-directionally upstream in a dual RNA cassette. The oBC enables robust detection of reporters in single cells, independent of reporter activity, enabling unbiased measurement of mBCs from the enhancer-driven reporter mRNA. See also **Fig. S1-S2**.

To resolve this problem, we developed a dual RNA reporter cassette which separates the detection and quantification tasks (**Fig. 1C**). For reporter detection, we introduce circularized (*37*) Pol III transcribed barcodes which enable near-complete recovery of the identity of the reporter(s) present in any given cell from single-cell RNA-seq data (scRNA-seq). Benchmarking this strategy in cell lines, we demonstrated accurate quantification of reporter mRNA levels over four orders of magnitude with a precision approaching the limit set by Poisson (shot) noise. We then profiled 204 candidate CREs drawn from 23 developmental loci in a stem-cell model of early mammalian development, mouse embryoid bodies. We confirm the specificity of previously characterized canonical elements controlling expression of *Sox2* in pluripotent cells, and discover numerous autonomously active constitutive and lineage-specific regulatory elements. Looking forward, we anticipate that this strategy will enable the scalable, quantitative characterization and dissection of enhancers in multicellular models of development.

## RESULTS

### A dual RNA cassette decouples the detection and quantification tasks in single-cell reporters

We reasoned that detection and quantification can be decoupled via two separate barcoded RNAs linked on individual reporters (**Fig. 1C**). In such a dual RNA cassette, one barcoded RNA, highly and constitutively expressed, serves as the marker for presence/absence of the integrated reporter within any given cell. The second RNA, a Pol II expressed <u>m</u>RNA <u>b</u>ar<u>c</u>oded (hereafter mBC) in its 3’ UTR, serves to quantify CRE activity and is equivalent to the reporter of bulk MPRAs. Provided that the two barcodes are *a priori* matched to one another, as well as to distinct CREs, reporter expression can be deconvoluted in single-cell assays with a dynamic range extending beyond the noise floor inherent to one-RNA approaches.

Dual RNA reporters require the contiguous production of two separate RNAs, which could interfere with CRE function. Given that Pol II promoters can act as enhancers (*38*), we expressed the detection barcode from a Pol III promoter. Interactions are expected to be minimal as a result of the largely orthogonal Pol III and Pol II machineries (the TATA-binding protein being the only shared factor across the two pathways (*39*)) (**Methods**). Our reporter architecture (**Fig. 1C, S1A**) places the hU6-driven detection barcode co-directionally upstream of the quantification cassette to avoid head-on collision (*40, 41*).

To mitigate the instability of short ectopic Pol III RNAs (*42*) and boost capture, we embedded the barcode and single-cell capture sequence within the ‘Tornado’ circularization system (*37*) (**Fig. S2A-B**), which requires no exogenous protein for function. The resulting circular RNA barcodes, referred to as Tornad<u>o b</u>ar<u>c</u>odes (oBC), were expressed at >150-fold higher steady-state levels compared to linear barcodes (**Fig. S2C**) driven by the same Pol III promoter (**Fig. S2D-E**, comparison performed via genome-integrated bulk MPRA, **Methods**), reaching an estimated >75,000 oBC RNA per cell per integrated cassette (**Methods**), in line with previous quantification (*37*). The impact of random barcode sequence on expression was minimal (≤2.6-fold interquartile range, **Fig. S2E**), confirming the robustness of the Tornado system.

The resulting <u>s</u>ingle-<u>c</u>ell <u>q</u>uantitative <u>e</u>xpression <u>r</u>eporters (scQers), each defined by three elements delivered to cells as a single unit – a detection oBC, a CRE, and a quantification mBC – enabled characterization of enhancer activity in multicellular systems.

### Benchmarking with a promoter library in human cell lines

We first established that scQers report transcriptional activity in single-cells with ≈2% dropout, high accuracy over a large dynamic range (<10^−1^ to >10^3^ UMI/cell), and high precision (coefficient of variation <1). To do so, we constructed a minimal library of five Pol II promoters spanning a wide activity range (*45*) (**Fig. 2A, Data S2**), and integrated the payloads by piggyBac (*46*) transposition at high multiplicity of integration (**Methods**) in three human cell lines (HEK293T, HepG2, K562). Cells were bottlenecked to a few hundred clones, expanded, and then both: 1) hand mixed at 1:1:1 ratios and profiled via scRNA-seq (10x Genomics 3’ feature barcoding with custom libraries optimized to increase reporter capture, **Fig. S1B-F, Methods**); and 2) harvested separately for bulk MPRA (**Fig. 2A, Methods**). Thousands of single cells per replicate passed standard quality filters, with cell line identity unambiguously mapped from gene expression (**Fig. 2B, S3A, Methods**).

**Figure 2:**
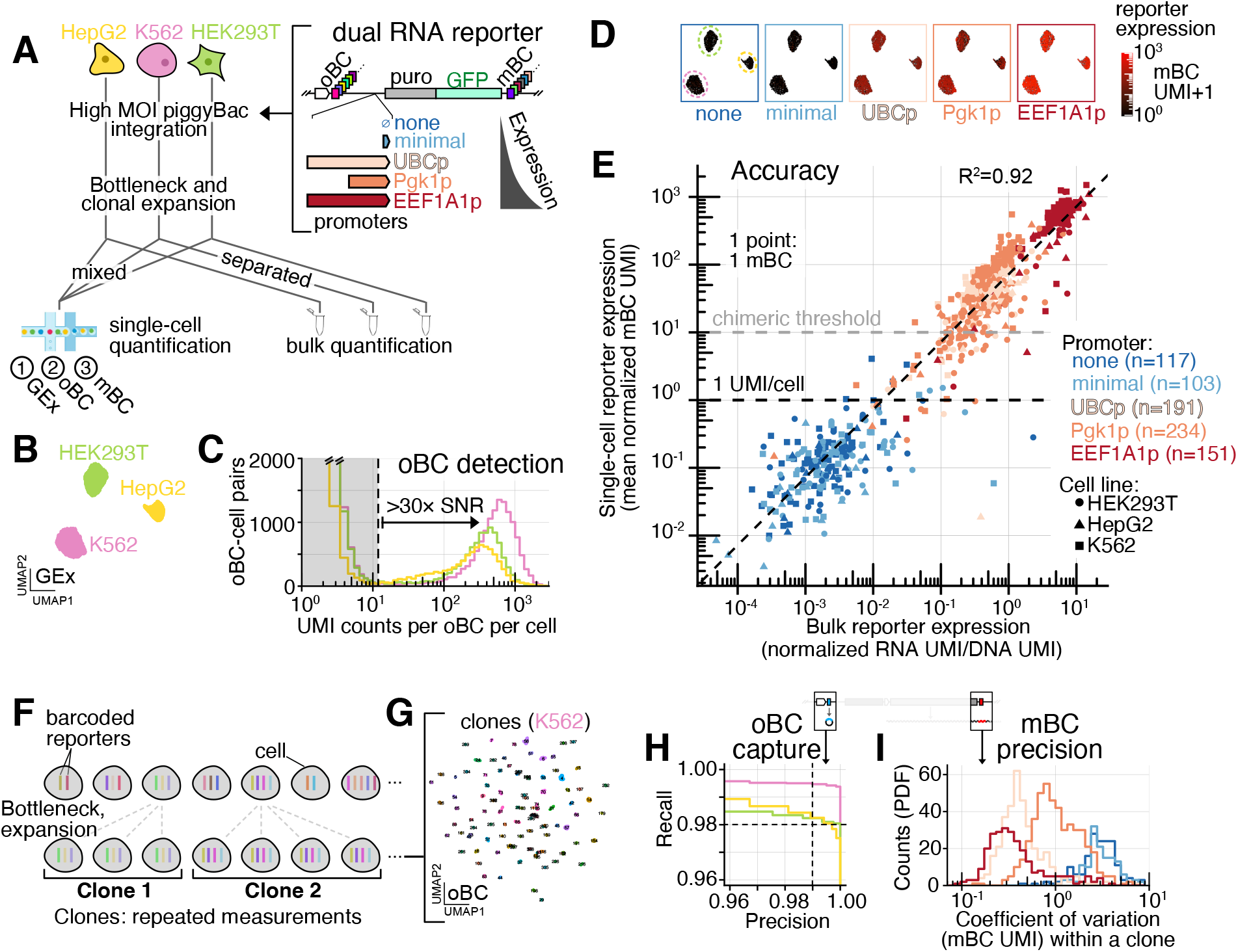
Benchmarking scQers for accuracy, precision, and capture in human cell lines. (**A**) Benchmarking experiment: A library of five promoters (n=1122 uniquely mappable oBC-promoter-mBC triplets, median 205 mBC-oBC pairs per promoter) was integrated in three human cell lines (HepG2, K562, HEK293T) at high multiplicity with the piggyBac transposase. Following integration, bottlenecking and expansion, clonal cells were: 1) separately bulk processed via MPRA; and 2) mixed at 1:1:1 ratio and single-cell profiled (**Fig. S1B**) to generate three libraries: gene expression (GEx), oBC, and mBC, from which the per-cell activity of each promoter can be quantified. (**B**) UMAP projection of quality filtered single-cell transcriptomes from the hand-mixed single-cell assay. The three well-separated clusters correspond to the three cell lines (replicate A; pass-filter cell count: K562 n=2184, HEK293T n=2090, HepG2 n=1231). (**C**) Distribution of the UMI counts per oBC per cell, stratified by cell line. The count distribution is bimodal, with a low-count mode (truncated, gray shading) corresponding to chimeric amplicons, and a high-count mode corresponding to *bona fide* integrations. (**D**) Layering reporter expression on transcriptomic state: UMAP projection cells colored by activity of each promoter (average normalized mBC UMI count for all reporters from the same promoter in each cell). Cell line identity marked in the first panel. Each panel corresponds to a different promoter. A pseudocount of 1 was added to display expression on a logarithmic scale. (**E**) Comparison between the single-cell mBC quantification (y-axis: average normalized mBC UMI over all cells with detected matched oBC, on average n=32 cells/mBC) and bulk MPRA quantification (x-axis, RNA over DNA normalized UMI counts). Each point corresponds to an individual mBC, coloured by its associated promoter. Symbols denote different cell lines. Well-represented mBC are included (>100 bulk DNA UMI, >0 measured mBC UMI in single cells, and ≥5 single-cell integrations detected). The diagonal dashed line follows a 1:1 slope. The chimeric (10 UMI/cell, **Fig. S3E**) and 1 UMI/cell thresholds are highlighted. R^2^ is computed from the log-transformed values. (**F**) Clonally derived cells with a high multiplicity of reporter integrations provide internally controlled replicates of the same measurement for assessing capture of oBC and precision of mBC quantification. (**G**) UMAP projection (oBC expression space) for high-confidence-assignment cells to clonotypes (**Methods**) for K562 (replicate A; n=1430 cells, n=105 clones). (**H**) Precision-recall curves for retrieval of oBC from cells assigned to clones across the cell lines, with consensus clonotypes taken as ground truth (aggregate over all clones with >2 cells assigned across two replicates; K562: 195 clones, 2168 cells; HEK293T: 173 clones, 2019 cells; HepG2: 38 clones, 1453 cells; **Methods**). Cell assignment to clones follows loose cutoffs (allowing for 50% oBC dropout), ensuring an unbiased assessment. Dashed lines: 99% precision (1% FDR), and 98% recall (2% false negative rate, or dropout). (**I**) Distribution of the coefficient of variation (CV; mean over standard deviation) for the normalized mBC UMI counts captured, measured across replicate clonal cells profiled, illustrating that reporter mRNAs driven by active promoters can be captured with low variability (CV<1). Each count corresponds to a reporter-clone pair (n=946 reporters from n=290 clones, across two biological replicates). See also **Fig. S3-S4**.

### oBCs are near-deterministically retrievable in scRNA-seq

oBCs were robustly captured on a per-cell basis. In particular, the distribution of oBC unique molecular identifier (UMI) counts displayed bimodality (**Fig. 2C, S3B**) with a large signal to noise ratio (>30× between minimum and high-count mode). The near-exponential low count mode corresponds to chimeric amplicons, and the approximately log-normal high-count mode to expression from *bona fide* integration events (≈2500 UMI/cell per barcode, zero-truncated Poisson estimator, **Methods**). To assess oBC dropout, we leveraged redundant measurements across clones (**Fig. 2F**). Consensus integration clonotypes were identified in the bottlenecked population by relying on oBC co-detections (*47, 48*) (**Methods, Fig. 2G, S4C-E, Data S7**). Clonotypes served as ground-truth for precision-recall analysis of detected oBCs in clone-assigned cells, revealing a false negative rate (dropout) of <2% at a false discovery rate of 1% (**Fig. 2H, S4A-B, S4D, S4F, Methods**). This represents a >10-fold improvement vis-a-vis capture of sgRNAs in single-cell CRISPR screens (*48*). In sum, oBCs are transcribed tags which effectively eliminate dropout in single-cell assays.

### Accurate reporter mRNA quantification over four orders of magnitude

Comparing reporter expression from single-cell and bulk quantification confirmed the accuracy of scQers. Following detection of reporter integration using oBCs (probability of multiple integrations per cell from the same oBC-promoter-mBC triplet <5%, **Methods**), activity of the associated promoters can be quantified in each cell as the transcriptome-normalized average UMI counts from the matched mBC (**Fig. 2D, S3C, Methods**). Single-cell averaged UMI counts across the different mBCs associated with a given promoter constituted independent measures of activity and spanned over four orders in magnitude for the five promoters (**Fig. 2E, S3D**). Bulk MPRA measurements performed on the same cell populations were quantitatively concordant across the full range of expression (R^2^ of log-transformed expression ≥0.87, **Fig. 2E, S3D**). Single-cell measurements of each mBC from as few as 5-10 cells sufficed for accurate quantification (**Fig. S3F**).

Without filtering, spurious read counts can alter reporter quantification. Indeed, library preparation requires a number of amplification steps that can generate ‘chimeric’ amplicons and lead to spurious cell-to-barcode connections. In saturated libraries, the signature for these molecular products is a rising frequency of counts below ≈10 UMI/cell (e.g., oBC:**Fig. 2C**, mBC: **Fig. S3E**) which can result in a limit of detection substantially higher than 1 UMI/cell. A dual RNA approach does not abrogate chimeras, but it filters mBC reads based on detection of a matched oBC in the same cell, leading to an average decrease in the tallying of chimeric counts by the proportion of cells harboring any given oBC-mBC combination. Consequently, lowly expressed mRNAs driven by the minimal and no promoter basal controls (median expression of ≈0.2 UMI/cell below the 1 UMI/cell regime inaccessible from pooled one-RNA reporters, **Fig. 2E**) remained accurately quantified by scQers, suggesting limited zero-inflation (*49*) in our system. Leveraging our *a priori* matched oBC-mBC pairs, we found that chimeric counts (mBC UMI found in cells without matched oBC detected) constituted a substantial proportion of the signal (chimeric fraction: 90% EEF1A1p, 60% Pgk1p, 51% UBCp, 36% no promoter, 52% minimal promoter). As a result, quantifying activity based on Pol II mBC alone (no conditioning on oBC detection) led to biases and increased variability (**Fig. S3G**, R^2^=0.39 for log-transformed single-cell vs. bulk; 1.5 to 25-fold increased variability, **Fig. S3H**), highlighting the quantitative advantage of dual RNA reporters.

### Measurement precision approaching Poisson counting noise

In addition to assessing oBC dropout, our clonal pool of cells allowed us to quantify variability in mBC capture. Multiply represented clones provide internal replicate measurements of the same set of reporters integrated at fixed genomic locations, controlling for an important source of variation for randomly integrated cassettes (*50*–*52*) (**Fig. 2F**). To assess precision of mBC quantification, we determined the variance in normalized UMI counts for each mBC across all cells assigned to a given clonotype (bottom rows of **Fig. S4G-H** for examples of the mBC UMI clonal distributions). Across all reporters and clones, we find variability consistent with Poisson counting noise at low expression, and a coefficient of variation (standard deviation/mean) substantially below one, at least for two of the three expressed promoters (UBCp and EEF1A1p, **Fig. 2I, S4I**). Variability was not strictly correlated with average expression. Promoter Pgk1p in particular, while expressed more highly than UBCp, exhibited substantially higher cell-to-cell variability (**Fig. S4I**). scQers thus precisely measure reporter mRNA levels.

### Systematic assessment of integration positional effects

Clonal analysis also informed on reporter expression variation driven by positional effects (assuming distinct clones harbor reporters integrated at different genomic locations). We observed promoter and cell line-specific effects, with EEF1A1p and UBCp showing remarkably little clone-to-clone variation (interquartile range across clones, UBCp: <2.4 for all cell lines; EEF1A1p: <1.5 in K562 and HEK293T, 4.1 in HepG2). In contrast, promoter Pgk1p showed both cell line differences in expression (e.g., median 12 normalized mBC UMI in HEK293T vs. 76 in K562) and higher variability across clones (IQR 4.8 in HEK293T, 5.9 in K562, 7.2 in HepG2). Decomposing the mBC UMI variability into positional effects (via clone assignment) vs. the sum of remaining biological and technical noise showed that precision was limited by genomic context, underscoring the low variability of our capture and the importance to average over multiple independent integration positions (fraction mBC UMI variance attributable to clone identity: EEF1Ap=0.60, Pgk1p=0.41, and UBCp=0.57, **Methods**). Still, for the three active promoters considered here, clone-to-clone variability was substantially lower than that of uninsulated reporters (*50*), suggesting that insulators included in our design (**Fig. S1A**) partially mitigated positional variegation.

### Locus-level screen of putative developmental enhancers

Following extensive optimization in cell lines, we sought to apply scQers to discover cell-type-specific enhancers in an *in vitro* model of early mammalian development, mouse embryoid bodies (*53, 54*) (mEBs). We drew putative CREs for testing from the neighborhood of empirically prioritized developmental loci (**Fig. 3A-B**). First, by profiling 21-day differentiated mEBs with scRNA-seq and single-cell ATAC-seq (*55, 56*) (scATAC-seq), we established the transcriptional and chromatin accessibility states of various cell types (**Fig. S5, Methods**). Of note, scATAC-seq data from mEB was highly correlated to *in vivo* data from matched cell types in E7.5-E8.5 embryos (*57*) (R^2^ of log-transformed accessibility across top 65k mEB peaks: e.g., parietal endoderm=0.77, neuroectoderm=0.78, mesoderm=0.76), supporting mEBs as a model of gene regulation in early development. Leveraging these data, we nominated 22 developmental genes with germ-layer specific expression and cell-type-specific chromatin accessibility landscapes (**Data S3, Methods**) such as endoderm regulator *Gata4 (58)*, other lineage-defining transcription factors (*Klf4, Foxa2, Sox17*), and structural genes (laminins, collagens, tubulin). As a comprehensive set (*59*) of CREs to profile from these genes, we selected all regions within ±100 kb of their TSS that were reproducibly and highly accessible in the expression-cognate cell type (e.g., 13 putative CREs near *Gata4* in **Fig. 3A**, other examples: **Fig. S6, 4A**). As controls, we additionally included the four constituents of the core *Sox2* control region (*60, 61*), accessible exclusively in pluripotent cells (**Fig. 3E**). In total, 209 elements were included for profiling (145/209 promoter-distal, i.e., >1 kb from promoters (*62*), median element size: 937 bp, **Data S3**). The five exogenous promoters (same as **Fig. 2A**) were also spiked-in as standards (10% of the transfection). Following library construction and sequential subassemblies (**Fig. S7**, 204/209 CREs represented with >20 oBC-mBC pairs, 88/145/242 10^th^/50^th^/90^th^ percentile number of valid oBC-mBC pairs per CRE), scQers were integrated in mESCs at high MOI using piggyBac (*63, 64*). Reporter-integrated cells were induced to form mEBs, sampled every 2 days for bulk MPRA quantification across differentiation, and scQer-ed at three weeks end-point (**Fig. 3B, Methods**).

**Figure 3.**
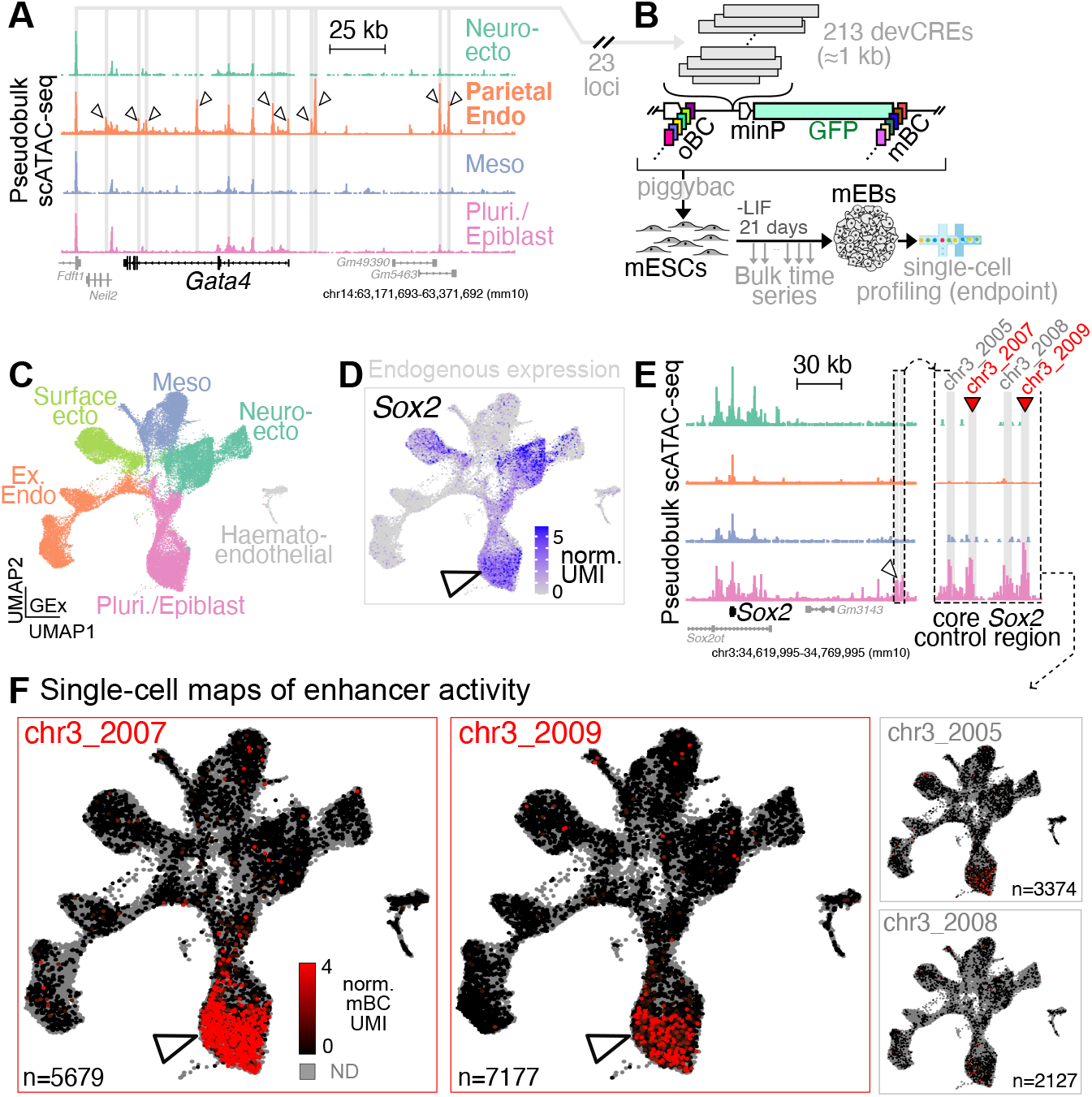
Locus-level screen of developmental enhancers in mouse embryoid bodies. **(A)** Pseudo-bulk pileup of scATAC-seq data at *Gata4* (±100 kb from TSS) as a representative selected developmental locus (carets: differentially accessible peaks). *Gata4* is expressed predominantly in parietal endoderm cells (expression **Fig. 4D**, top row). All reproducibly and highly accessible ATAC peaks (in expression-cognate cell-type) within the 200 kb window were included (n=13 for *Gata4*, gray shading). (**B**) scQers containing 204 putative developmental CREs taken from 23 developmental loci (22 plus Sox2 control region) were integrated at high MOI in mESC using piggyBac. Transfected libraries included 89% CRE series, 10% exogenous promoters (same as in **Fig. 2A**), and 1% constitutive EEF1A1p-mCherry (co-transfected to increase MOI (*63, 64*), **Methods**). Reporter-integrated cells were differentiated to embryoid bodies for 21-days, with bulk sampling every 2 days, and single-cell profiling at three weeks. (**C**) UMAP projection of scRNA-seq (n=43799 quality-filtered cells) from three biological replicates of scQer-integrated 21-day mEB cells, with annotation from integration with *in vivo* data (*65*) (finer cluster resolution **Fig. S5A**), confirming diversity of cell types. (**D**) Endogenous expression (normalized UMI counts) for *Sox2* displayed on UMAP projection, highlighting pleiotropic expression in pluripotent (caret) and ectodermal lineages. (**E**) scATAC pseudobulk pileup for *Sox2* locus. Caret points to the *Sox2* control region (*60, 61*), inset zooms in the core. Regions profiled and differentially accessible in the pluripotent population are shaded in gray. Red carets mark the two cell-type-specific enhancers. (**F**) Single-cell maps of enhancer activity derived from scQers for four CREs (separate panels). Each point represents a single cell. Gray indicates cells with no reporter detected for the specified CRE. Color marks reporter expression (average normalized mBC UMI per cell) from none (black) to high (red) for cells with detected reporters (oBC UMI>10). Color axis truncated to 4 UMI to highlight low mBC UMI counts. Elements chr3_2007 and chr3_2009 have significant expression specific to pluripotent cells (caret) (**Methods, Fig. 4A**, marginal activity from chr3_2005 significant in only 1 of 3 biological replicates), mirroring *Sox2* expression in that cell type (c.f., panel D). Number of cells with detected reporter integrations indicated on each panel. See also **Fig. S5**-**S9**.

### High performance of scQers in a stem-cell derived developmental system

mEBs reproducibly comprised diverse cell-types unambiguously mappable to *in vivo* germ-layers (*65*) (**Methods, Fig. 3C-S5A, S5C**, n=43799 pass-filter cells across three biological replicates [replicates 1 and 2: separate transfections; replicate 2B: ∼500-clone bottleneck of replicate 2 with 12% identified clonotypes overlap to replicate 2, and thus largely orthogonal; all replicates separate mEB inductions], **Methods, Fig. S8C**), confirming successful differentiation despite the presence of reporters at high MOI (**Fig. S8D-E**, median MOI = 23, probability of any oBC-CRE-mBC triplet integrated more than once per cell=1%).

scQers displayed high performance in mEBs. First, oBC were robustly captured (median library complexity=836 UMI/oBC/cell), with a bimodal distribution of oBC UMI/cell (**Fig. S8F**). oBC expression was cell-type independent (**Fig. S8G**), enabling uniformly high recovery (<4% oBC dropout at FDR=1% from precision-recall analysis of clonal cells, **Fig. S8I-K, Methods**). Second, comparison of end-point bulk and single-cell quantification across profiled CREs confirmed accuracy of reporter expression measurement over the full dynamic range (R^2^ log-transformed activity=0.81, **Fig. S8A**) and per-cell-type quantification was reproducible (R^2^ log-transformed across replicates=0.72, **Fig. S8B**). Representation was reasonably uniform across tested CREs (**Fig. S8H**, captured integration events per element 1597/3153/6197 10^th^/50^th^/90^th^ percentiles, and n=17971 to 34745 for exogenous promoters).

### Single-cell maps of activity for the core Sox2 control region enhancers

scQer generated high-contrast single-cell maps of CRE activity (**Fig. S9A**). As a case study, we considered gene expression control of the pleiotropic regulator *Sox2* (**Fig. 3D**). *Sox2* is a key factor in pluripotency maintenance whose dysregulation leads to aberrant differentiation (*61*). Central to *Sox2* control is a distal (≈135 kb from TSS) cluster of CREs necessary for driving high expression in pluripotent cells (*60, 61*), and previously shown to function autonomously (*61, 67*). Of the four differentially accessible elements in pluripotent cells from this core *Sox2* control region (**Fig. 3E** inset), two displayed robust activity (red **Fig. 3F**, 10 to 30-fold higher expression vs. minimal and no promoter basal controls), in agreement with previous characterizations (*8, 61*) (**Fig. S9D, Data S4**). Activity was circumscribed to the pluripotent population (>50-fold higher expression vs. other cell types, e.g., **Fig. S9B** for *Sox2*:chr3_2007). While *Sox2* was expressed in the pluripotent and ectoderm lineages in mEBs (**Fig. 3D**), CREs from the *Sox2* control regions were exclusively active in the pluripotent population (*Essrb*/*Dppa3*-positive cells (*66*), **Fig. S5B**). *Sox2* expression in other cell types thus likely arises from other regulatory elements, such as promoter proximal neural-specific enhancers (*68*). Our results on a previously characterized enhancer cluster confirm that scQers can report cell-type specific expression in a multicellular system with high sensitivity and contrast.

### Systematic identification of active CREs

Beyond the *Sox2* control region, we quantified both activity and cell-type specificity of other tested CREs (n=200), identifying multiple active elements (**Fig. 4A, S10**). For each CRE, average reporter expression was determined across cells with detections, stratified by cell-types. CRE <u>activity</u> was defined as the maximum per-cell-type reporter expression, while CRE <u>specificity</u> was taken as the maximum per-cell-type mBC expression divided by the mean expression in all other cells (**Fig. 4A**). We identified 58/204 endogenous CREs with activity in significant excess of the basal controls in all three replicates (**Methods, Data S5**). The elements with the highest expression were the active exogenous promoters (UBCp, Pgk1p, EEF1A1p) with levels ≈300× to ≈2500× above the basal controls (≈30 to 250 mBC UMI/cell, **Fig. 4A, S10A**). Active endogenous CREs spanned a wide range at lower expressions (maximum per-cell-type expression: ≈0.3 to 20 mBC UMI/cell, **Fig. 4A**). Notably, a sizable fraction (19/58) of the active CREs had expression under 1 mBC UMI/cell, and almost all were below the chimeric read threshold of 10 UMI/mBC/cell.

**Figure 4.**
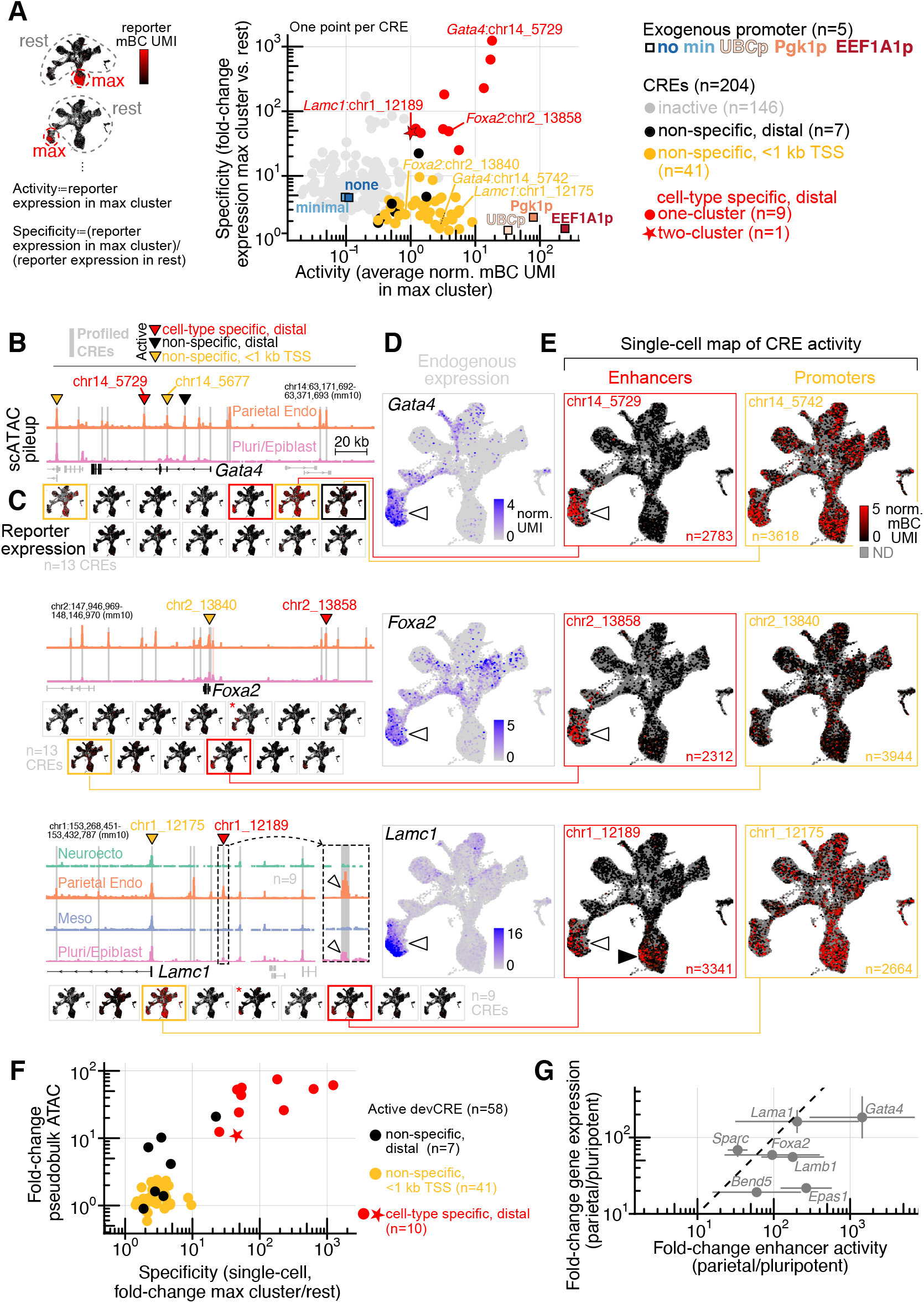
Multiplexed identification of constitutive and autonomous lineage-specific CREs. **(A)** Two-dimensional coarse-grained space of CRE function. <u>Activity</u>: reporter expression (average normalized mBC UMI count) in the maximum-expression cell-type (defined from fine clusters of **Fig. S5A**, illustrated left). <u>Specificity</u>: maximum-expression cell-type reporter level over expression in all other cells (fold-change). Graph shows median quantification across three replicates. Active elements (black: non-specific, distal; orange: non-specific, <1 kb TSS; red: cell-type specific) are identified as having excess expression (bootstrap resampling, **Methods**) in all replicates compared to basal controls (no and minimal promoter, blue) in contrast to inactive elements (gray). Active elements found to have cell-type-specific expression (specificity >5 and significantly higher than cell-type permuted sets, **Methods**) are highlighted (red). CRE *Lamc1*:chr1_1218, found to be active in two cell types, is marked with a star. Exogenous promoters (same as **Fig. 2A**) serving as internal standards are shown as colored squares. Elements shown in panels B and E are indicated. Panels (B-E) are reproduced across rows for the different loci (top to bottom:*Gata4, Foxa2, Lamc1*). (**B**) Pseudobulk pileup of scATAC (pluripotent and parietal endoderm: *Gata4, Foxa2*, also neuroectoderm and mesoderm for *Lamc1*) for 200 kb region centered on gene transcription start site. Gray shading of peaks indicate regions tested in the multiplexed assay (red shaded peak near *Foxa2* TSS: peak not present in the library due to inability to identify specific cloning primers). Carets point to elements identified as active with scQers (same coloring as panel A). Inset for *Lamc1* locus highlights differential accessibility in both pluripotent/epiblast and parietal endoderm cells (white carets), matching the activity profile of the element. (**C**) Single-cell reporter activity maps for all tested elements in the locus. Outline indicates activity of element in assay (same coloring as panel A). Red asterisk mark elements with activity identified in less than 3/3 replicates. (**D**) Endogenous expression (scRNA-seq, normalized UMI counts projected on UMAP) for genes corresponding to loci shown. Caret points to the parietal endoderm cells, displaying differential expression. (**E**) Single-cell reporter expression (normalized mBC UMI, projected on UMAP) for putative promoter (orange) and distal enhancer (red) associated with the gene in the locus. Panels have the same color scale (truncated at 5 mBC UMI to highlight contrast). Shown elements correspond to those labeled in panels A-B. Number of cells with detected reporters per element is indicated. White carets point to parietal endoderm. Black caret (*Lamc1*:chr1_12189 element) marks reporter expression in pluripotent cells, which does not match endogenous expression of the putatively associated gene *Lamc1*. (**F**) Fold-change in ATAC (cognate cluster vs. rest of cells) vs. single-cell reporter expression specificity (definition and color scheme, panel A) for all active elements identified. (**G**) Fold-change in gene expression (y-axis, ratio normalized UMI in parietal endoderm to pluripotent) vs. enhancer induction (x-axis, fold-change reporter levels, average normalized mBC UMI in parietal endoderm over pluripotent) for parietal-endoderm-specific distal enhancers. Dashed line is 1:1. Geometric mean over biological replicates is shown (errorbar: standard deviation of geometric mean). See also **Fig. S10**-**S15**.

Active CREs displayed distinct expression patterns across mEB cell types. Categorizing active CREs as cell-type-specific vs. non-specific via a permutation test (**Methods**), we found 10/58 developmental CREs with reproducible cell-type-specific activity (single-cell activity maps, red in **Fig. 4A-C, S11A-B**). Of the remaining 48 non-specific active elements, 41 (85%) were promoter-proximal (e.g., orange **Fig. 4E, S11D**) compared to 0/10 of cell-type-specific CREs. Conversely, 41/62 tested promoter-proximal elements were found to be active and non-specific (while 0/62 were cell-type-specific). Consistent with their function and distance from TSS, all cell-type-specific enhancers showed >10 fold-change in chromatin accessibility in their cognate cell types, whereas promoters were constitutively open (<3 fold-change, **Fig. 4F**) Single-cell activity maps thus delineated two broad patterns of autonomous function exhibited by accessible regions at developmental loci (**Fig. 4E, S11D**): constitutively active elements (overwhelmingly TSS-proximal) and cell-type-specific elements (overwhelmingly TSS-distal).

Our assay relies on high MOI random integration of reporters for scalable multiplexing, raising concerns that genomic positional effects might dominate the signal (*50, 51*). To assess positional effects, we bottlenecked reporter-integrated mESCs to a few hundred clones in one of the replicate (replicate 2B, **Methods**) prior to mEB induction. Quantifying activity of the 10 cell-type specific enhancers across clones (assuming different integration positions), we found that for most CREs (9/10) retained specificity (>5-fold) across the super-majority (over two-thirds) of well-represented clones (**Fig. S12, Data S8, Methods**), suggesting that positional effects can be averaged over.

### Characterization of lineage-specific, autonomous enhancers

Of the 10 autonomous cell-type-specific enhancers identified, two belonged to the core *Sox2* control region (**Fig. 3F**), while the remaining 8, all from distinct parietal endoderm-expressed loci (red **Fig. 4E, S11D**), included a *Gata4* intronic enhancer 10 kb downstream of the first exon (chr14_5729, **Fig. 4E** second row) and an enhancer 70 kb upstream of *Foxa2* (chr2_13858, **Fig. 4E** third row). One active element at the *Lamc1* locus (chr1_12189, **Fig. 4E** fourth row) was found to be bi-functionally active in two cell types, with concordant chromatin bi-accessibility (inset **Fig. 4B** fourth row). Of note, that we mostly identified endoderm-specific CREs was not unexpected given the uneven sampling of tested elements, in part a result of the high proportion of endoderm cells in the scATAC data restricting power in other cell-types.

Reporter expression driven by developmental CREs mirrored the predominant pattern of expression of their nearby putatively associated gene (**Fig. 3D** vs. **3F, 4D** vs. **4E, S11C** vs. **S11D**, systematic per cell type quantification: **Fig. S13**), except for the bi-functional putative *Lamc1* enhancer (**Fig. 4D** fourth row, black caret), which drove expression in both parietal endoderm and pluripotent cells, in contrast with endogenous *Lamc1* whose expression was restricted to parietal endoderm. For parietal endoderm-specific enhancers, the magnitude of induction was on par with endogenous gene induction (**Fig. 4G**). What proportion of endogenous regulation do the identified autonomous enhancers recapitulate? This question is difficult to directly address because absolute reporter UMI counts cannot be uniformly compared to gene expression UMI counts (*i*.*e*. due to gene-to-gene differences in conversion between endogenous mRNA levels and captured UMI counts). Taking activity of the active promoter putatively associated with the induced gene (orange in **Fig. 4E, S11D, S13**) as baseline (with the caveat that mRNA levels driven by promoters in our reporter system might not be perfectly reflective of endogenous activity), we found that the activity of the autonomous enhancers captured a substantial proportion of the expression fold-change, but in 6/7 cases less than a half (shaded **Fig. S11E**), as perhaps expected for multi-CRE landscapes.

Leveraging our time-resolved bulk MPRA (**Fig. S14, Data S6**) on the same samples (scQers with bulk readout on mBC), we found a consistent set of active CREs (53/54 bulk active elements identified as active from scQers, 53/58 scQers identified elements found as bulk active, **Methods**). Importantly, elements found to be cell-type-specific with scQers displayed either temporal increase (red **Fig. S14D**), decrease (core *Sox2* control region, **Fig. S9C**), or non-monotonic behavior (bifunctional CRE, *Lamc1*:chr1_12189 **Fig. S14D**), supporting their classification as developmental enhancers. In contrast, active but non-specific elements displayed little temporal variation across differentiation (e.g., exogenous promoters **Fig. S14C**; endogenous elements, orange **Fig. S14D**), as expected for constitutive promoter-like CREs (**Fig. S14B**).

A number of features were enriched in the 8 active cell-type specific enhancers within all 103 tested distal parietal endoderm elements tested. Active CREs displayed higher chromatin accessibility (1.8-fold more accessible, 2.2-fold more differentially accessible, both p<0.03 B-H corrected one-sided t-test), but showed no difference in evolutionary conservation (average phyloP score (*69*)), nor were they significantly closer to the TSS of their putative target gene. Indeed, at all loci, the autonomously active CRE was not the closest element from the TSS (**Fig. 4B, S11A**). Active elements also showed no evidence of opening earlier than other elements in a pseudotime analysis (*70*) (**Methods**), arguing against them being ‘seed enhancers’ (*71, 72*). With regards to finer-level sequence features, active CREs contained a higher density of endodermal regulator Gata4 binding sites, but only if considering binding sites of intermediate-to-high affinities (between 1.3 and 2.2-fold more binding sites for relative affinity lower thresholds between 0.2 and 0.45, p<0.03 B-H corrected one-sided t-test, 8-mer affinities from Uniprobe (*73*–*75*), **Methods**, binding sites also elevated in neighboring 500 bp windows ±100 kb from TSSs, **Fig. S15C**). While additional examples are needed to draw general conclusions, this suggests clusters of intermediate affinity binding sites of key regulators might be important for mammalian developmental enhancer function, in line with the suboptimization hypothesis (*29, 76*). Two other endodermal regulators, Foxa2 and Sox17, did not show a higher number of binding sites in active CREs. In short, active parietal endoderm CREs displayed significantly elevated ATAC accessibility and Gata4 transcription factor binding sites (**Fig. S15A**), with a logistic classifier using these two properties accurately classifying active/inactive elements (auROC=0.94, **Fig. S15B**, precision=0.6 at recall=0.75).

Overall, scQers enabled the scaled high-sensitivity characterization of both constitutive promoter-like and lineage-specific autonomously active regulatory elements across diverse cell types of 21-day mouse EBs, with enhancer activity profiles matching expression of their putatively associated genes.

## DISCUSSION

Enhancers are believed to orchestrate the precise unfolding of development in metazoans, enabling the emergence of a species’ form and function from a genomic blueprint. However, at least to date, our ability to study developmental enhancers at scale has been constrained, particularly in mammalian systems. On one hand, *in situ* transgenics (*28*–*30*) demonstrate that even 1 kb noncoding sequences can encode patterns of remarkable cell-type and spatiotemporal specificity, but these assays are not readily scalable to the vast numbers of enhancers involved in development (*11*). On the other hand, MPRAs, which are readily scalable, are largely limited to static, homogenous cell lines.

We and others (*77*) have recognized that a simple path forward is to intersect MPRAs with an increasingly sophisticated landscape of single-cell resolution technologies, *e*.*g*. scRNA-seq. Here we overcome the technical challenges of combining these two modalities, resulting in scQers, a new class of MPRA that decouples the detection and quantification of reporters via a dual RNA system and circularization-based enhancement of barcode recovery. scQers extend measurements into a regime fundamentally inaccessible with traditional multiplex reporters, yielding an accurate, precise and high-contrast readout of reporter mRNA levels. In mouse embryoid bodies, scQers permitted the pooled profiling of 204 ≈1 kb-long elements taken from 23 developmental loci, identifying 10 cell-type specific enhancers, several of which autonomously drove a >100-fold increase in activity in their cognate cell types, relative to a rigorously measured baseline. While most of the autonomous enhancer elements identified here displayed expression domains mirroring that of their putatively associated gene, in-genome perturbations will be necessary to confirm their role, if any, in endogenous regulation.

The relatively low enhancer hit rate of our screen suggests that genome integration followed by differentiation prior to measurement provides a strong filter for elements autonomously competent to reconfigure chromatinized landscapes. Indeed, episomal assays as applied in other model systems can report a greater proportion of active elements (*78*) (e.g., *Lama1*:chr17_7791 contains a parietal endoderm enhancer as identified by episomal reporters (*79*) that was not reproducibly functional in our genome-integrated assay). Beyond these technical differences, given the complex multi-enhancer landscapes considered here, some tested CREs might contribute to regulation, but only in the presence of (or by directly serving as) cooperating elements, in line with recently described facilitators (*9*) or chromatin-dependent enhancers (*13*) (e.g., tested but inactive *Sox2*:chr3_2005, which overlaps with facilitator DHS23 (*8*)).

What is the advantage of a single-cell assay over multiple bulk assays performed in a variety of cell lines? Developmental systems display a continuum of states, contrasting with discontiguous, terminal states. Constructing maps of enhancer activity along developmental manifolds has the potential to reveal the effects of subtle changes in the milieu of *trans*-acting factors, enabling finer assessment of function and dysregulation. As predictive models of enhancer activity become more refined (*13, 20, 24, 26*), quantitative experimental approaches are needed to efficiently iterate through design-test-learn loops to validate underlying mechanistic hypotheses. Benchmarks in cell lines and a proof-of-principle screen in a multicellular stem-cell model establish scQers as a scalable platform for enhancer biology and should be portable to numerous other developmental systems (e.g., zebrafish (*80*), *C. intestinalis (29)*, the chicken neural crest (*78*), sophisticated stem-cell models like synthetic embryoids (*81, 82*), or *in vivo* neuronal subtypes with AAV derivatives (*83*)).

## Acknowledgments

We thank N. Ahituv, M. Kircher, R. Ziffra, G. Gordon, A. Ellis, J. Tome, and the entire Shendure lab for discussions; participants of the gene regulation subgroup (F. Chardon, W. Chen, X. Li, T. McDiarmid) for criticisms and advice; T. McDiarmid for noting the high instability of non-complexed sgRNAs. Plasmid pAV-U6+27-Tornado-Broccoli was a kind gift from S. Jaffrey (Addgene plasmid # 124360).

## Author contributions

JBL and SGR conceptualized dual reporters. JBL cloned scQer libraries, planned and carried out experiments in human cell lines and Pol III MPRA. SGR and JBL planned and carried out experiments in mEBs. JBL analyzed data, generated figures, and wrote the manuscript with edits from JS and comments from SGR and DC. SGR generated scATAC data in mEBs. SGR and SD generated the mESC line, established mEBs protocols, and performed early profiling of mEBs. BM provided constructs, and protocols for cloning of MPRA cassettes. DC suggested analyses and provided computer scripts for subassembly. TL performed bioinformatic analyses on CREs. CCS provided starting protocols for library subassembly. CL provided assistance with FACS. CT and JS supervised the study.

## Data availability

Raw sequencing data and processed files generated in this study have been deposited to GEO, with accession number GSE217690. Code and scripts used for analyses have been deposited on github (<u>shendurelab/scQers</u>), together with the maps of plasmids and custom sequencing amplicons structures used in this work. Already published data used: transcription factor binding data (Uniprobe (*73*): Gata4 (*74*), Sox17 (*75*), Foxa2 (*75*)), mouse embryo *in vivo* scRNA-seq (*65*) (obtained from R library: “MouseGastrulationData”) and scATAC-seq (*57*) (GEO: GSE205117).

## DATA TABLES

**Data S1**: oligos and plasmids used in this work.

**Data S2**: sequences of exogenous promoters used for benchmarking and as internal standards (**Fig. 2A**).

**Data S3**: details of profiled CREs (positions and sequences tested, **Fig. 3A-B, 4A, S10B**).

**Data S4**: positions of core SCR elements tested in this and previous works (**Fig. S9D**).

**Data S5**: quantification of activity and specificity for all CREs measured with single-cell reporter (**Fig. S4A**).

**Data S6**: quantification of activity for all CREs from bulk MPRA time series (**Fig. S14**).

**Data S7**: high confidence clonotypes and clonotype-assigned cells, human cell line experiments (**Fig. 2G, S4**).

**Data S8**: high confidence clonotypes and clonotype-assigned cells, mEB experiments (**Fig. S8I-K**).

## SUPPLEMENTARY FIGURES

**Figure S1:**
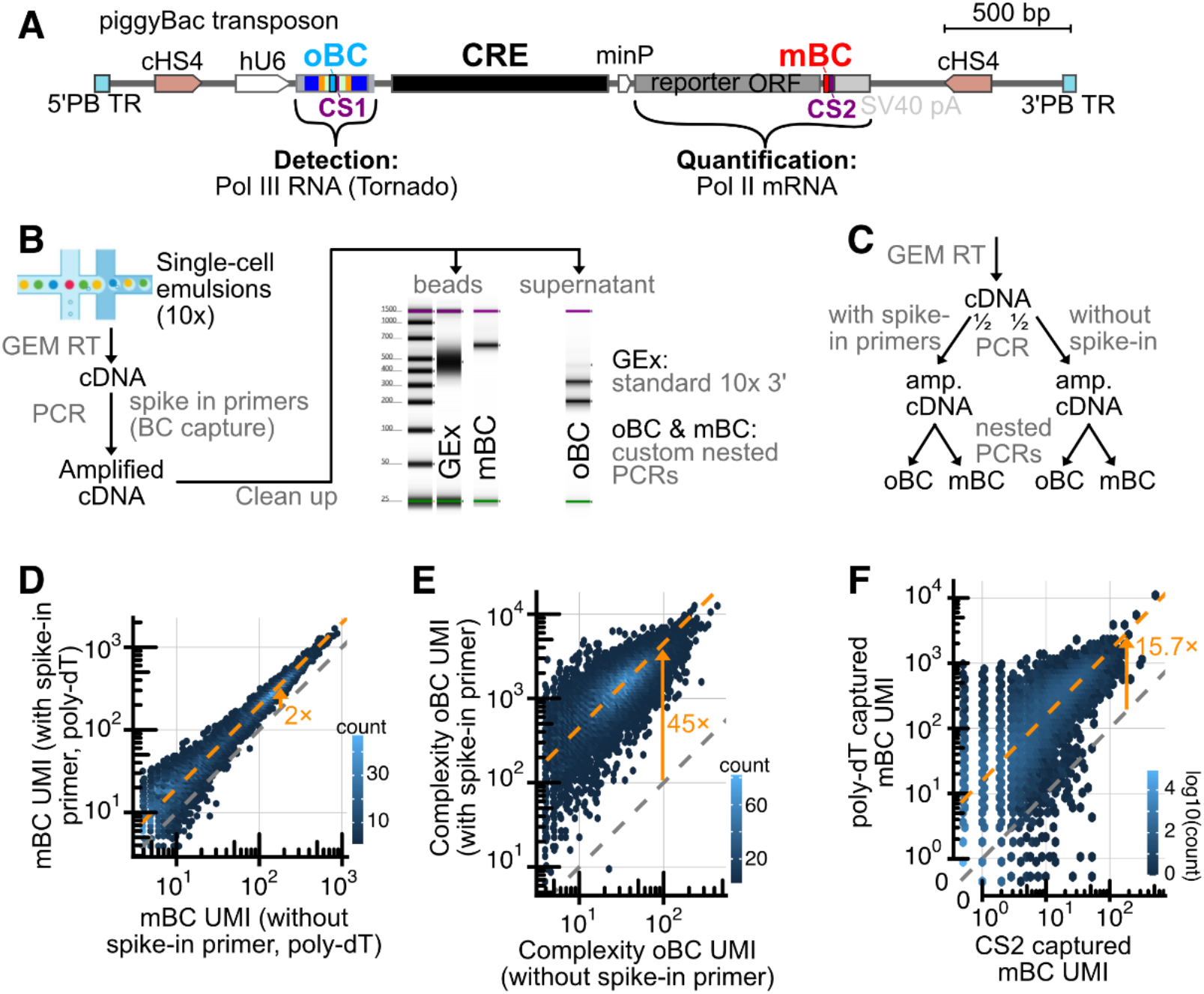
Dual RNA reporter cassette, single-cell assay, and barcode capture optimization. **A** At-scale schematic of the dual RNA reporter cassette in piggyBac transposon (between terminal repeats: PB TR). Flanked by convergent insulators (core chicken hypersensitive site-4 from beta-globin locus, cHS4 (*43*)), the human U6 (hU6) driven Tornado barcode cassette (oBC-CS1, details shown in **Fig. S2A-B**) is co-directionally placed upstream of the CRE library driving an open reading frame-containing reporter transcript (puromycin-P2A-GFP in the case of the promoter series in cell lines, **Fig. 2A**, and GFP alone for mEB experiment, **Fig. 3B**), barcoded in its 3’ untranslated region upstream of an inserted capture sequence 2 (CS2), and of the SV40 polyadenylation sequence (SV40 pA). **B** Schematic of the single-cell reporter assay. After 10x Genomics (V3.1, 3’ gene expression with feature barcode) GEM reverse transcription, primers (specific to oBC and mBC RNAs) are spiked-in the cDNA amplification mix (*44*). Post-cDNA amplification, in addition to standard gene expression (GEx) library generation, nested PCRs from bead fraction (mBC) and supernatant (oBC) are performed to obtain custom single-cell reporter libraries. Amplification of barcodes proceed from different fractions as reporter mRNAs harboring the mBC are long (>800 bp), purifying with the beads, whereas oBC are short (134 bp), remaining in the supernatant. Representative tapestation traces of resulting libraries are shown (showing laddering products from oBC libraries). **C** Experiment to assess improvement in UMI capture by spiking in primers in initial cDNA amplification. For the experiment with promoter series in cell lines (**Fig. 2A**), replicate B’s cDNA was split in two prior to cDNA amplification. One half, replicate B1, received spike-in primers to the oBC and mBC reporters, and the other half, replicate B2, did not. An additional round of PCR downstream of the first cDNA amplification was performed to obtain libraries in replicate B2 (**Methods**). **D** and **E** Comparison of number of UMIs captured for the same cell barcode and reporter barcodes between replicates B1 (with spike-in primers) and B2 (without spike-in primers) for mBC (panel D: 2.0× median increase in UMIs captured, orange arrow. n=8395 mBC-cell barcode pairs with >3 UMI) and oBC (panel E: 45× median increase in UMIs captured, orange arrow. n=19323 oBC-cell barcode pairs with >3 UMI), respectively. The higher boost in capture resulting from spike-in primers for the oBC vs. mBC was likely due to the circular nature of the barcode: given the absence of 5’ end from which template switching can occur from oBC RNAs, the initial cDNA amplification (primed from the template switching oligo) effectively cannot happen except from the linear intermediates towards oBC formation, presumed to be at much lower abundance; in contrast, the spike-in primers directly target sequences flanking the barcode in the circular oBC. **F** Comparison of captured mBC UMI from poly-dT vs. capture sequence 2 (CS2) on-bead reverse transcription primers (for the same mBC-cell barcode pairs). As expected from primer stoichiometry on beads, >15× increase (orange arrow) in captured mBC UMI is seen from the poly-dT vs. CS2 primers (n=21492 mBC-cell barcode pairs with poly-dT and CS2 mBC >0 across both replicate A and B1). CS2 thus adds marginal value for capture of the Pol II-derived polyA-tailed mBC transcripts.

**Figure S2:**
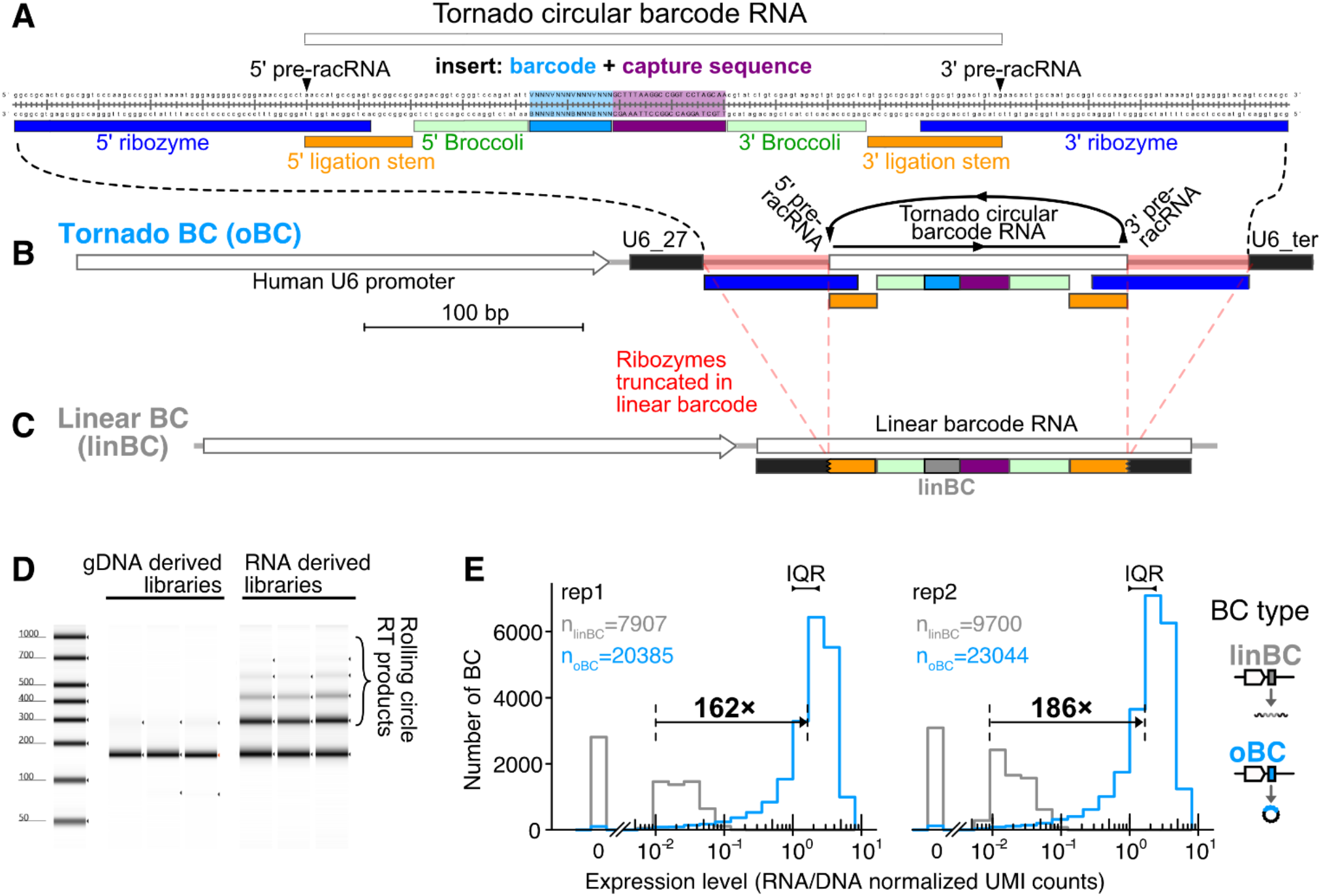
Tornado barcodes are highly expressed Pol III driven RNAs. **A** Sequence of the Tornado system (*37*) with 16 bp barcode (5’ VNNNVNNNVNNNVNNN, light blue) and downstream capture sequence 1 (CS1; burgundy) inserted in the loop of Broccoli. 5’ and 3’ (pre-racRNA) ends cleaved by ribozymes prior to circularization are highlighted (black carets). The circular product is 134 nt long. **B** and **C** Schematic of the human U6 promoter driven cassettes tested in a head-to-head MPRA experiments (integrated via piggyBac; **Methods**) to compare expression of the circular version of the barcode (Tornad<u>o b</u>ar<u>c</u>ode, or oBC, B) to the linear barcode (<u>lin</u>ear <u>b</u>ar<u>c</u>ode, linBC, C), which is the same construct but with ‘Twister’ ribozymes removed (red highlight in B). **D** Representative tapestation traces of genomic DNA-derived vs. RNA-derived amplicon libraries prepared from the oBC vs. linBC MPRA experiment. RNA-derived libraries show clear rolling circle reverse transcription products laddering of the expected periodicity (+134 bp) expected from circular RNAs. **E** Distribution of MPRA-derived activity estimates (RNA/DNA normalized UMI) for the thousands of different, well-represented (>50 DNA UMI) barcodes of both types (hU6-driven oBC [blue] vs. hU6-driven linBC [gray]) as assessed by bulk MPRA, highlighting both the large difference in steady-state expression (>150× difference in median between linBC and oBC), and tight distribution (interquartile range <3×) for the oBC. Sub-panels correspond to two independent biological replicates.

**Figure S3.**
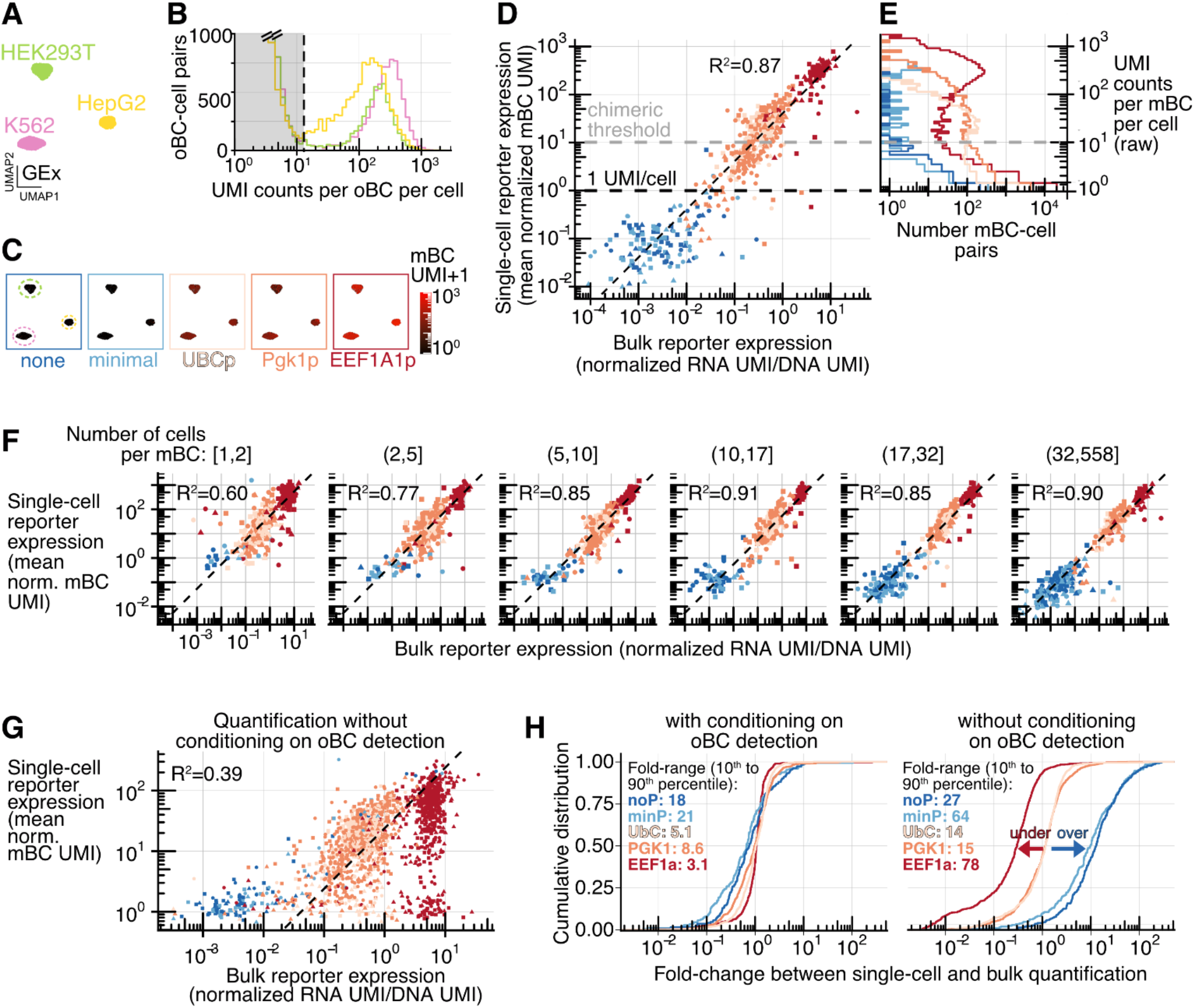
Assessment of accuracy of single-cell dual RNA reporters. **A-D** Same as **Fig. 2A-C**, but with data from replicate B1. A: Gene expression, B: oBC UMI count distribution, C: single-cell measure of reporter expression in single-cells (GEx UMAP projected), D: comparison of bulk vs. single-cell quantification of mBC quantification. **E** Raw distribution of UMI counts per mBC per cell barcode (for valid mBC and cell barcodes pairs, not conditioning on oBC detection) stratified by associated promoter. The 10 mBC UMI/cell threshold (“chimeric threshold”) reflects that even for highly expressed promoters, mBC UMI counts rise below that point, as a result of chimeric amplicons generated during library preparation. Without conditioning on oBC detection, these molecular species limit the dynamic range of reliable measurements with one-RNA reporters (see panel G). **F** Assessment of reporter mRNA measurement accuracy vs. number of integration events captured (both replicates). Single-cell vs. bulk quantification (same as **Fig. 2E** and **S3D**), but stratified by the number of cells per mBC over which the single-cell measurement is averaged (split in equal number of mBC bins). Even with as few as 5 to 10 cells captured per mBC, the correspondence with bulk measurement is on par with estimates from more highly represented mBCs (R^2^ on log-transformed values ≥ 0.85). **G** Single-cell vs. bulk quantification of mBC expression without conditioning on oBC detection (assuming all mBC capture events are valid, both replicates). In contrast to oBC conditioned measurements, quantification has a hard floor at 1 UMI/cell (slight variation around 1 from gene expression normalization) and a limited dynamic range (y-axis spans ≈200× compared to >10^4^× with oBC conditioning, c.f., **Fig. 2E** and panel D). Only well-represented mBC are included (same criterion as **Fig. 2E**: >100 DNA UMI bulk, ≥5 cells with mBC detected). Dashed line marks the 1:1 slope, highlighting systematic biases. **H** Cumulative distribution of fold-change between single-cell and bulk mBC quantification (median normalized), for both replicates, with (left) and without (right) conditional oBC detection. While the quantification conditioning on oBC is largely unbiased (centered and close to 1), quantification is biased at the high (underestimation for highly expressed EEF1A1 promoter, red arrow) and low (overestimation for low expression minimal/no promoters, blue arrow) ends of the expression spectrum. In addition to removing systematic biases, conditioning on oBC also reduces variability (quantified as the spread in fold-change, with the range spanned from 10^th^ to 90^th^ percentile for each promoter displayed on plot).

**Figure S4.**
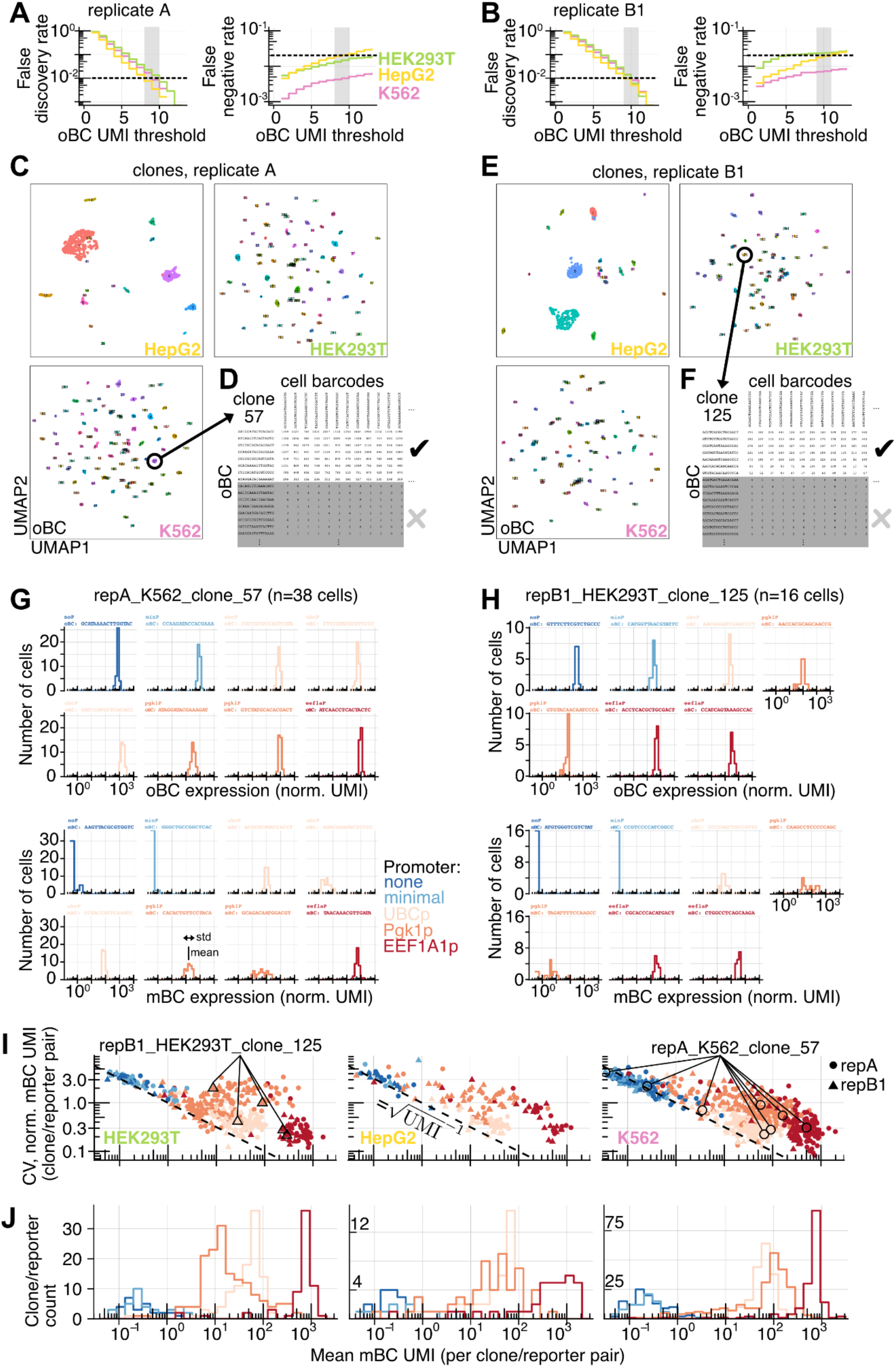
Benchmarking oBC detection and mBC capture precision with clonal analysis. **A-B** Systematic analysis of oBC dropout across all high-confidence clones. False discovery rate (left, false positives/[true positives + false positives]), and false negative rate (right panels, false negatives/[false negatives + true positives]) as function of the oBC UMI threshold used for detection. Analyses are performed on high-confidence clones represented by at least 3 cells. Consensus reconstructed clonotypes (**Methods**) are taken as ground truth and cells are assigned to these clonotypes with stringent threshold to remove doublets, but loose threshold to allow for up to 50% oBC dropouts per clone. At an FDR of 1% (gray shading), there are about 2% dropout (false negative rate) observed (slightly reduced performance from replicate B1 likely from halved complexity, see **Fig. S1C**). Panel A: replicate A, Panel B: replicate B1. **C** and **E** oBC expression space UMAP from cells assigned to high-confidence clones (colored by mapped clone identity) with at least three cells assigned, separated by cell lines. Panel C: replicate A (K562: 105 clones, 1430 cells; HEK293T: 92 clones, 1330 cells; HepG2: 17 clones, 916 cells), Panel E: replicate B1 (K562: 90 clones, 738 cells; HEK293T: 81 clones, 689 cells; HepG2: 21 clones, 537 cells). **D** and **F** Example of raw (error corrected) UMI counts (table truncated) per cell barcode and oBC across assigned cells in clones highlighted respectively in panels C and E (oBC ordered from high to low counts). Panel D: clone repA_K562_clone57 with 38 cells assigned. Panel F: clone repB1_HEK293T_clone_125 with 16 cells assigned. Grey shading delineates oBCs not assigned to the clones, highlighting the sharp distinction in UMI counts. **G** and **H** Example of mBC (top) and oBC (bottom) UMI count distributions across all cells assigned to specific clones (highlighted in panels C and E). Each sub-panel corresponds to a reporter integrated in the clone. Panel G: clone repA_K562_clone57, with 8 integrated reporters. Panel H: clone repB1_HEK293T_clone_125, with 7 integrated reporters. Panels in respective positions within the oBC and mBC set are matched (e.g., in repA_K562_clone57, EEF1A1 promoter with oBC:ATCAACCTCACTACTC and mBC: TAACAAACGTTGATA). **I** Coefficient of variation analysis of mBC UMI count measurements across all reporter-clone pairs stratified by cell line (left: HEK293T, middle: HepG2, right: K562). Mean over standard deviation (see panel G bottom: Pgk1 promoter with mBC:CACACTGTTCCTACA as schematic of both quantities) of normalized mBC UMI counts for reporters in clones as a function of mean normalized mBC UMI (reporters with >0.05 mBC UMI mean expression in clones with >4 cells assigned; replicate A: K562: 392 reporters from 83 clones, HEK293T: 198 reporters from 70 clones, HepG2: 58 reporters from 12 clones; replicate B1: K562: 213 reporters from 58 clones, HEK293T: 123 reporters from 51 clones, HepG2: 95 reporters from 14 clones). Dashed line indicates the Poisson counting scaling CV=√(UMI count)^-1^. Each point represents the quantification for a specific reporter within a clone, with point shape marking replicates and color promoter type. As examples, reporters shown in panels G (clone repA_K562_clone57) and H (clone repB1_HEK293T_clone_125) are highlighted in black (no and minimal promoter reporters from repB1_HEK293T_clone_125 have 0 mBC UMI and therefore do not appear). **J** Assessment of position-dependent variability of integrated reporters. Panels show the distribution in mean normalized mBC UMI (expression) across reporters integrated over different clones, stratified by cell line (left: HEK293T, middle: HepG2, right: K562) and promoter type (color). Same clone/reporter pairs as panel I. To account for halved library complexity in replicate B1 (see **Fig. S1C** description), reporter expression values from those clones were multiplied by two (most of the variability from some promoters otherwise coming from this technical factor).

**Figure S5.**
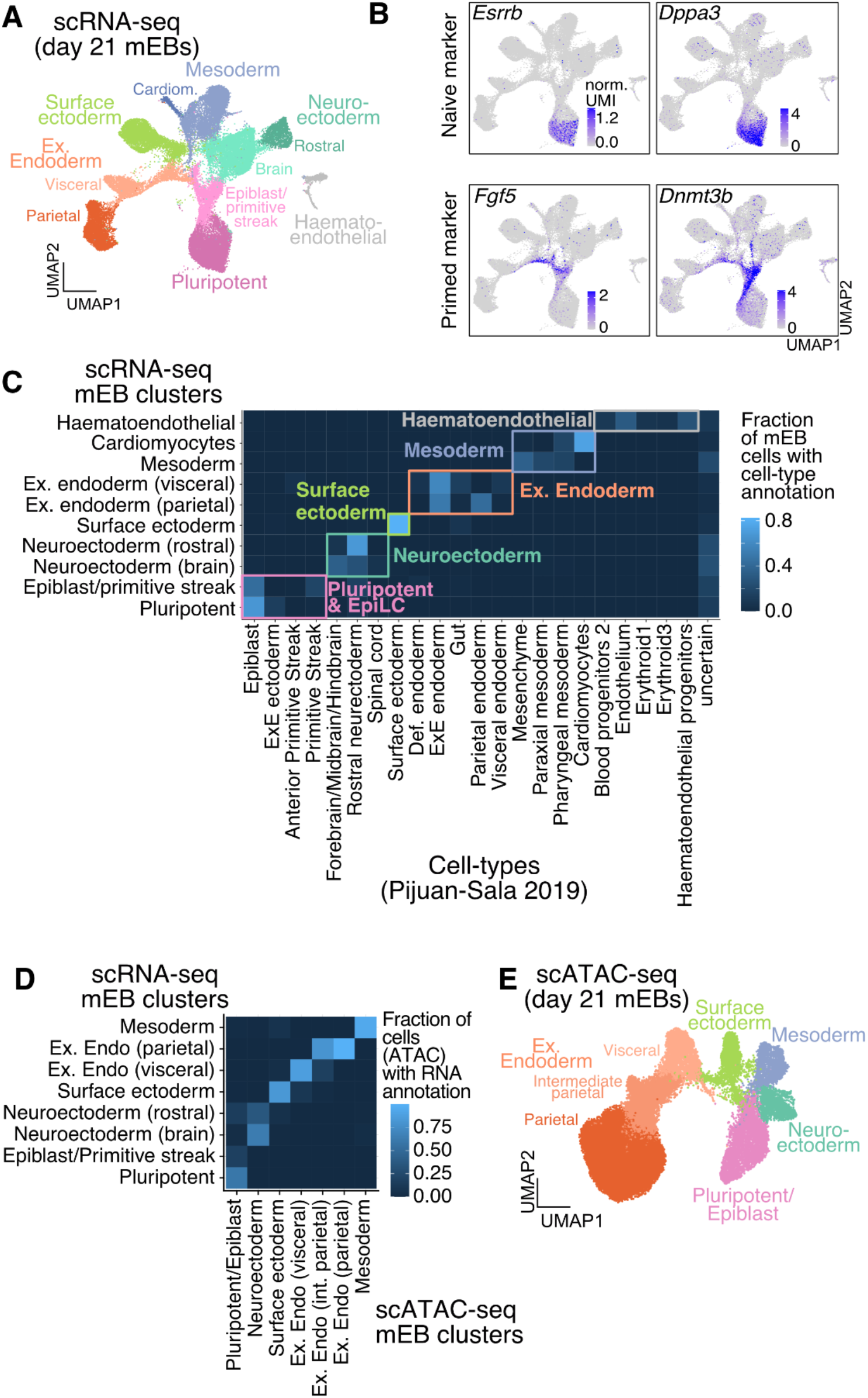
Molecular profiling and integration of single-cell data from 21-day mouse embryoid bodies. **A** UMAP of scRNA-seq data from quality-filtered cells from scQer-integrated, day 21 mEBs (same as **Fig. 3C**) annotated with fine cell types derived from label transfer of *in vivo* dataset (*65*), as shown in panel C. These cluster definitions are used to quantify CRE activity over cell types (e.g., **Fig. 4A, S9B, S13**). **B** Example of naive and primed pluripotent stem cell marker gene expression (normalized UMI counts) displayed on UMAP, used to annotate the respective cells as pluripotent and epiblast/primitive streak. **C** Heatmap displaying fraction of mEB-derived cells (from each cluster in panel A) with label transferred (**Methods**) to cell-types from *in vivo* data from Pijuan-Sala *et al (65)*. Cell types with no associated cells in mEBs (with maximum fraction < 5%) are not shown for brevity. Clusters coarse-grained for representation (**Fig. 3C**) are boxed. Uncertain column corresponds to cells that had ambiguous label transfer. The mEB cluster marked as pluripotent was manually annotated from specific expression of canonical marker genes (*66*) in those cells (panel B) as a result of a lack of naive mESC in the integration dataset. **D** Integration of scATAC-seq and scRNA-seq for cluster annotation. Heatmap showing fraction of nuclei from scATAC-seq-derived clusters predicted to be from cell-type identified in scRNA-seq data (**Methods**), displaying unambiguous matches. Cell types not found to be major clusters in scATAC-seq data are not shown. **E** UMAP of scATAC-seq data from quality filtered cells (n=46408, two biological replicates) from day 21 mEBs. Clusters are labeled based on integration with scRNA-seq data (panel A, panel E).

**Figure S6.**
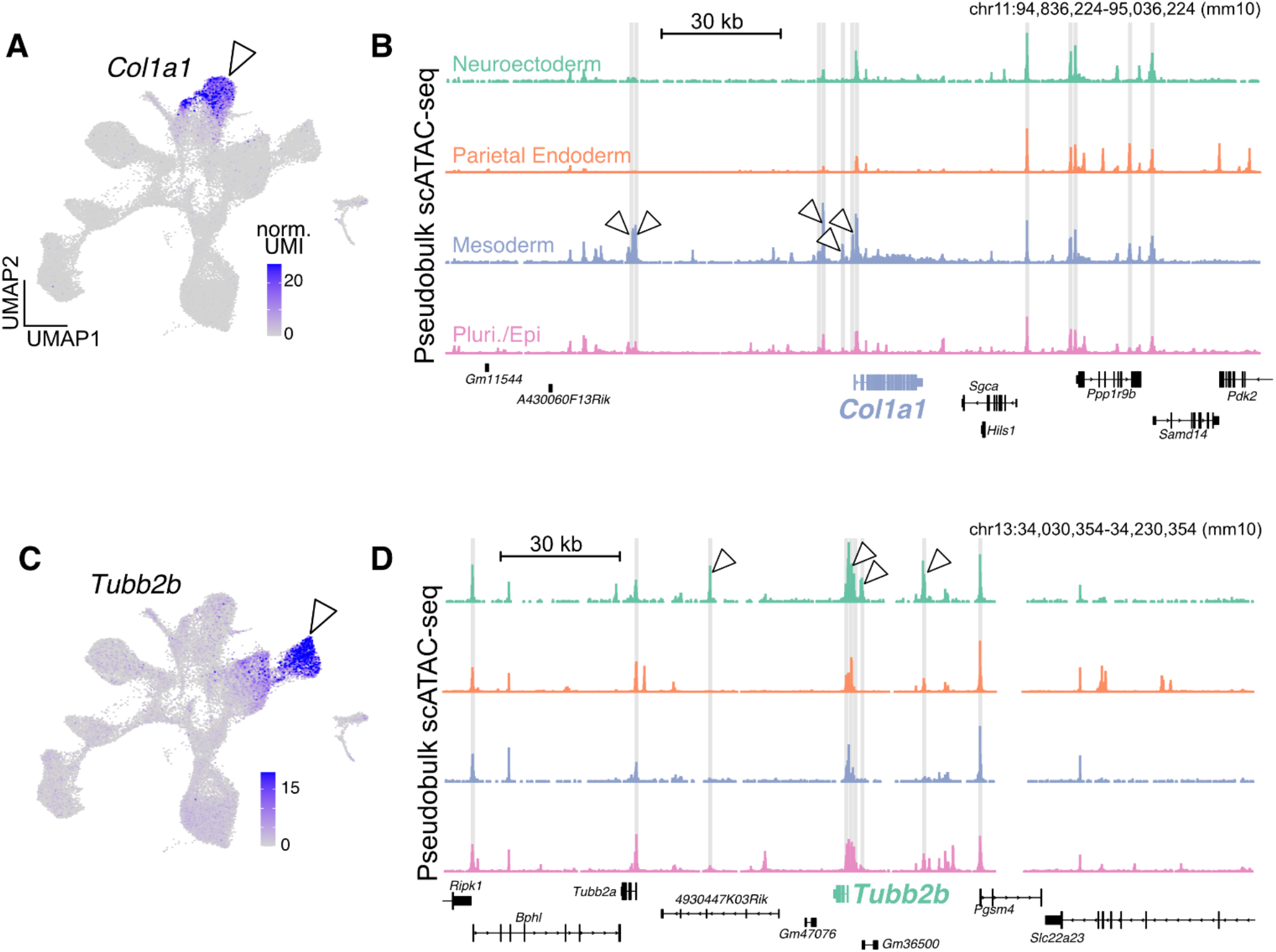
Additional examples of developmental loci with putative CREs selected for profiling. **A** and **C** Example of differentially expressed genes (carets, *Col1a1* expressed in mesoderm, and *Tubb2b* expressed in neuroectoderm respectively) selected as part of the 22 developmental loci for CRE selection. Gene expression normalized UMI counts for respective genes are shown on UMAPs. **B** and **D** scATAC-seq pseudobulk pile up (±100 kb from TSS) for genes shown on the left. Elements selected for screening are shaded in gray (c.f., **Fig. 3A**). Differentially accessible peaks are marked by carets.

**Figure S7.**
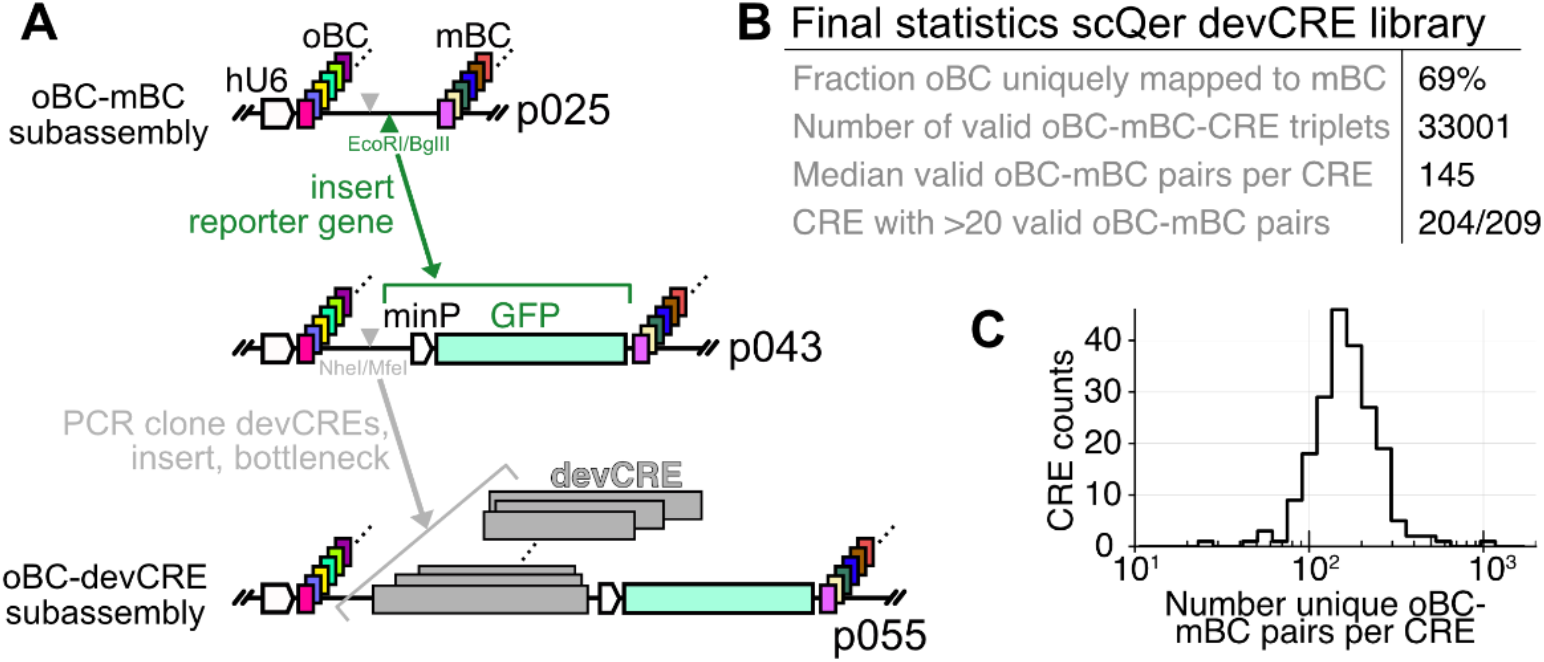
scQer library construction and oBC-CRE-mBC subassemblies. **A** Schematic of procedure to construct doubly barcoded dual RNA reporters. First, a high-complexity (∼1 M) library of doubly barcoded (oBC and mBC, separated by multiple cloning site dock) piggyBac transposons is constructed. At this step, oBC and mBC matches are determined (PCR-based library construction, **Methods**). The minimal promoter with GFP cassette is then inserted, and complexity maintained as much as possible. >200 CREs were PCR-cloned (**Methods**), pooled at 1:1 ratios by mass, and inserted in the doubly barcoded minP-GFP backbone by isothermal assembly. The resulting library was bottlenecked to ∼50k clones. CRE and oBC matches were then determined on the bottlenecked library (tagmentation with semi-specific PCR, **Methods**). In combination with the initial oBC-mBC pairs, this completes the determination of oBC-CRE-mBC triplets needed to deconvolute single-cell data for reporter activity. Plasmid names (p025, p043, p055) are indicated. **B** Compilation of statistics from scQers library used to screen putative CREs in mEBs. **C** Distribution of number of unique oBC-mBC pairs per CRE following the subassembly and quality filters, displaying largely uniform representation of the >200 putative regulatory elements tested (experiment **Fig. 3B**).

**Figure S8.**
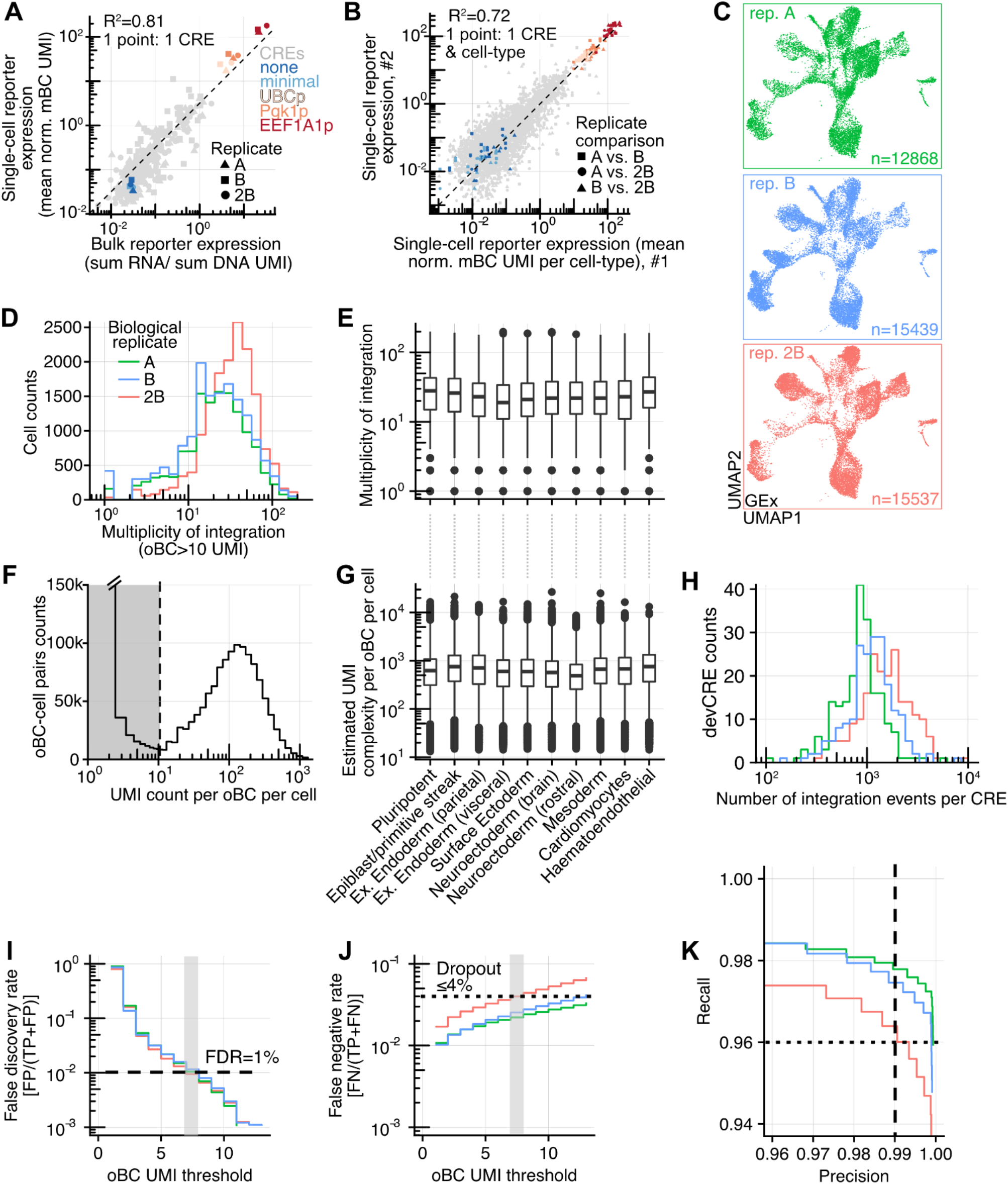
Quality metrics of single-cell reporter assay in mEBs. **A** Comparison between single-cell (average gene expression normalized mBC UMI count across all cells with detected reporter) and bulk quantification (day 21 samples, RNA/DNA ratio of summed, 1% winsorized, UMI counts across all barcodes) for well-represented CREs (>100 integrations and >30 total mBC UMI in single-cell assay and >35 mBC with at least 20 DNA UMI in bulk assay). CREs (gray) and promoters coloured according to **Fig. 2A**, dashed marks a 1:1 slope. R^2^ on log-transformed values across all replicates. **B** Comparison of per-cell type reporter quantification (average normalized mBC UMI over cells in clusters of **Fig. S5A**) across biological replicates for CREs with >0 activity. Each point corresponds to a CRE in a cell-type (10 points per CRE). Symbols mark different replicate pairs compared (e.g., squares for x-axis replicate A vs. y-axis replicate B). **C** scRNA-seq UMAP (same as **Fig. 3C, S5A**) stratified by biological replicate (no batch correction) showing reproducibility of cell-types obtained in embryoid bodies derived from reporter-containing mESC. Number of cells for each replicate indicated in each panel. **D** Distribution of multiplicity of integrations (a number of oBC with >10 UMI per cell) across individual cells and stratified by replicate (median: repA=20, repB=19, rep2B=31). High MOI in rep2B likely results from further selecting mCherry+ cells (1% co-transfection), not performed for replicates A and B. **E** Distribution (box plot) of multiplicity of integration stratified by cell types (see **Fig. S5A**). Cell type annotations same as in panel G. **F** Distribution of oBC UMI counts per cell (similar to **Fig. 2C**) highlighting robust circular barcode RNA capture in differentiated cells. Sharp bimodality and high signal-to-noise enables high-recovery reporter integration detection. **G** Box plot of estimated total UMI complexity (zero-truncated Poisson) for each captured oBC (>10 UMI) in all cells stratified by cell type, displaying similar levels irrespective of cell type. **H** Distribution of number of captured integration events per CRE (not including exogenous promoter series, determined from oBC UMI >10 from oBC-associated CRE) stratified by replicates, showing reasonably uniform coverage across profiled elements. **I-K** Precision-recall analysis of oBC detection (similar to **Fig. 2H, S4A-B**) for mEB-derived cells. Despite only replicate 2B being directly bottlenecked, replicates A and B also displayed (modest) clonal expansion (**Methods**), which enabled analysis of oBC dropout in these samples as well. High-confidence clones with at least two assigned cells are included (repA: 600 clones, 3977 cells; repB: 635 clones, 6465 cells; rep2B: 325 clones, 8518 cells), with results unchanged if restricting to more highly represented clones. Consensus clonotypes served as ground truth for analysis. Panels H and I respectively show the false discovery rate (FP/[FP+TP]) and false negative rate (FN/[FN+TP]) as a function of the UMI threshold used to assign barcodes to cells. At 1% FDR, false negative (dropout) is less than 4%. oBC libraries from replicate 2B were not sequenced as deeply (average saturation 6.0% vs. 18.7%), suggesting that part of the dropout is due to incomplete sequencing coverage and that dropout is below 4%.

**Figure S9.**
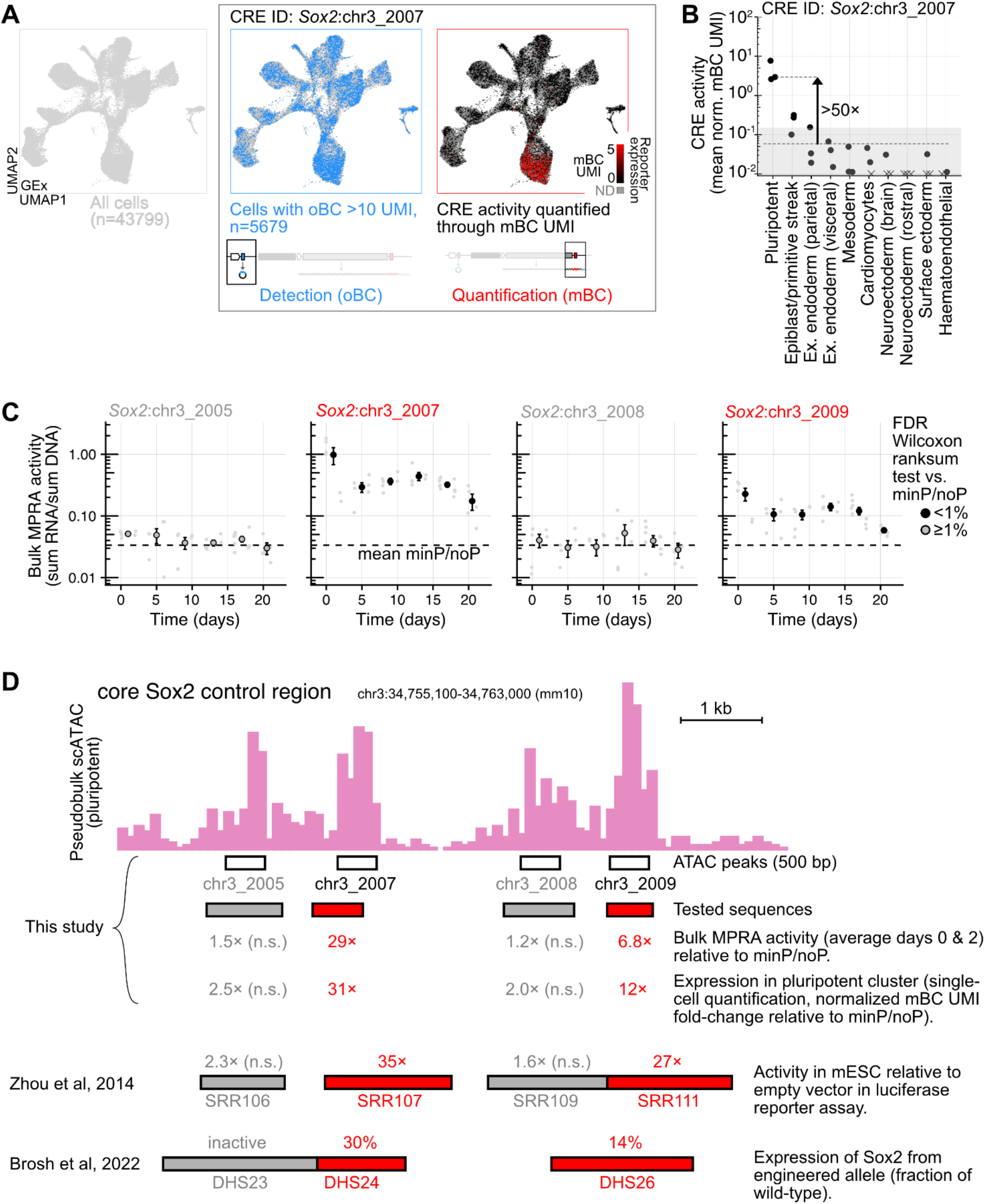
Details on activity of constituent elements of the Sox2 control region. **A** Illustration of the steps to construct a single-cell map of enhancer activity for a given regulatory element. Left: All cells passing quality filters are initially considered. Middle: Reporter detection. The list of oBCs associated with the CRE of interest (here *Sox2*:chr3_2007, see **Fig. 3F**) from the predetermined oBC-CRE-mBC triplets are identified. Cell barcodes with one (or more) CRE-associated oBC with >10 UMI are retained (n=5679), shown in blue on the UMAP (gray corresponding to cells with no detected reporters to the CRE of interest). Right: Expression quantification. From the oBC-CRE-mBC triplet table, the UMI counts to CRE-associated mBC are collected. In cases where multiple reporters to the same CRE (but different oBC-mBC pairs) are detected in the same cell, the average mBC UMI is taken. To correct for the fact that some cell types have more RNA (or other technical factors), we normalize the mBC expression by the total UMI to the transcriptome for each considered cell (**Methods**). The resulting single-cell reporter expression can then be layered on the low dimensional projection (black low to high red), enabling visualization of enhancer activity across the manifold of cell states in the system. **B** Quantification of the average reporter expression (average normalized mBC UMI, see panel A) across cells from different cell types (defined as clusters in **Fig. S5A**). Each dot corresponds to a biological replicate. Crosses correspond to cell types/replicates with average expression below 0.01 mBC UMI/cell. Arrow marks the fold change in expression between the maximum cluster (pluripotent) and the rest of cells (defined as specificity in **Fig. 4A**). Gray shading marks the noise floor determined from variability from the basal expression controls (minimal and no promoter). **C** Bulk MPRA quantification of the four constituents of the core *Sox2* control region (see **Fig. S14** for all CREs), showing consistent results with single-cell quantification (inactive: *Sox2*:chr3_2005, *Sox2*:chr3_2008; active: *Sox2*:chr3_2007, *Sox2*:chr3_2009). Small gray points mark individual replicates and time points. Large points are the average over replicates from consecutive time points, and are filled if significantly above the basal expression controls (ranksum test, B-H corrected, <1% FDR). Error bars show the standard error of the mean. Dashed line indicates the mean of basal expression control (minimal and no promoters). The observed decrease in activity over time for *Sox2*:chr3_2007 and *Sox2*:chr3_2009 is consistent with pluripotent cells being progressively depleted from the population, thereby leading to decreased activity when averaged over all cells in bulk. **D** Sox2 control region scATAC pseudobulk pileup in pluripotent/epiblast cluster (reproducing **Fig. 3E**). Under pileup, elements tested (in the same genomic position reference frame as the pileup, **Data S4** for positions) are indicated both from this study (top: 500 bp regions peak from ArchR pipeline; bottom: PCR-amplified tested sequences, Methods), and two previous studies quantifying reporter activity, Zhou et al (*61*), and Brosh et al (*8*). Gray regions were not found to be significantly active. Red regions were found to have activity in pluripotent cells (measured activity is indicated). *Sox2*:chr3_2007 from this study was not entirely nested in previously tested elements (SRR107 and DHS24), suggesting that even higher activity than measured might be achievable with a more inclusive element. The slight misalignment from the ATAC peak for *Sox2*:chr3_2007 resulted from lack of identifiable specific PCR cloning primers in the immediate 3’ region.

**Figure S10.**
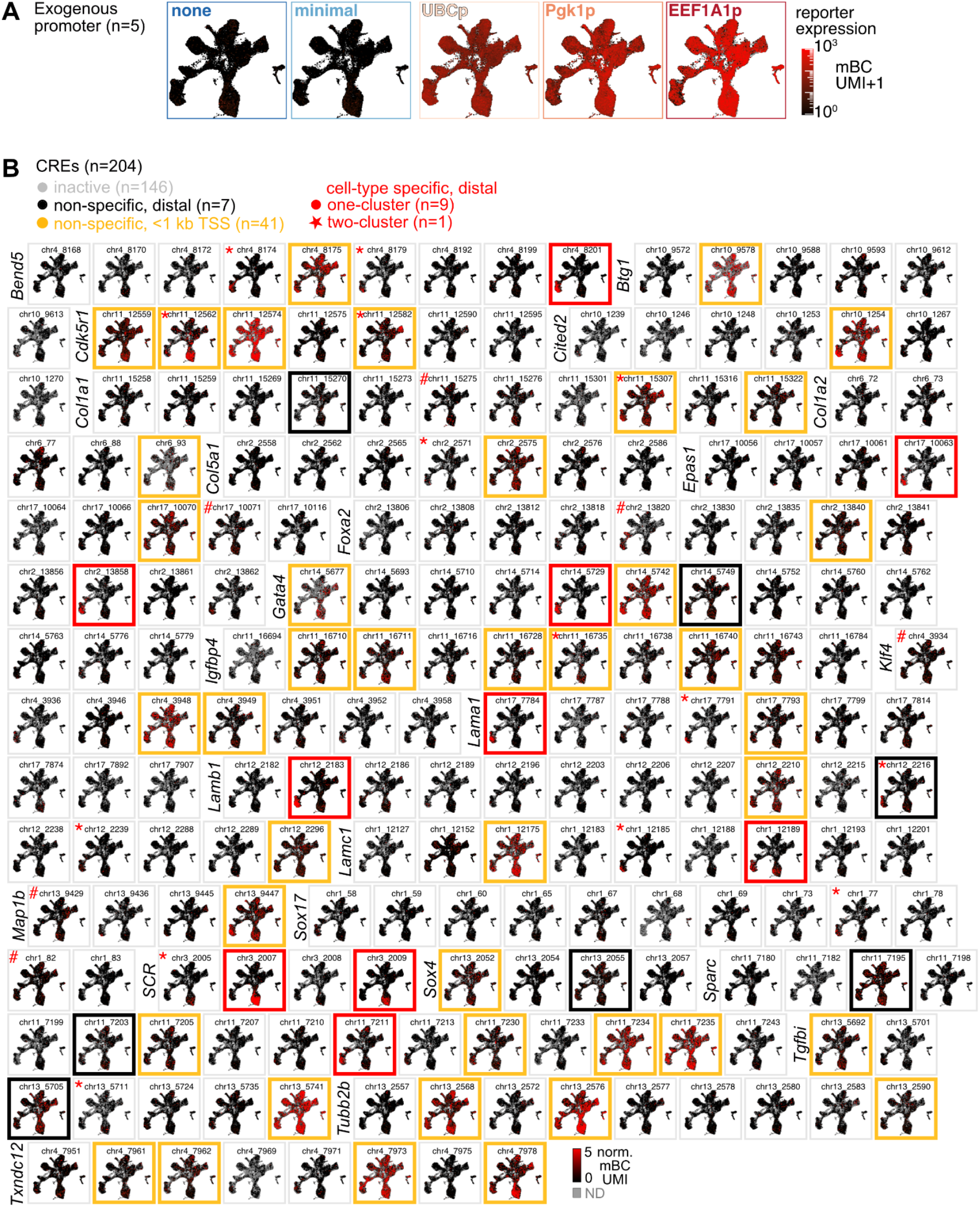
Systematic characterization of 204 putative CREs in mouse embryoid bodies. **A** Single-cell reporter expression (average normalized mBC UMI per cell) for the five exogenous promoters used as internal controls. Color scale is logarithmic (with a pseudocount of 1). **B** Single-cell reporter expression maps for the 204 profiled CREs. Elements are organized by locus (horizontally). Map outlines indicate the element class as classified in the two-dimensional phenotypic space from **Fig. 4A**. Elements marked with # are found to be active (non-specific) in 2/3 replicates. Elements marked with * are found to be active and specific in at least one replicate with our thresholds. Each map is shown to the same color scale (normalized mBC UMI from 0 and truncated to 5).

**Figure S11.**
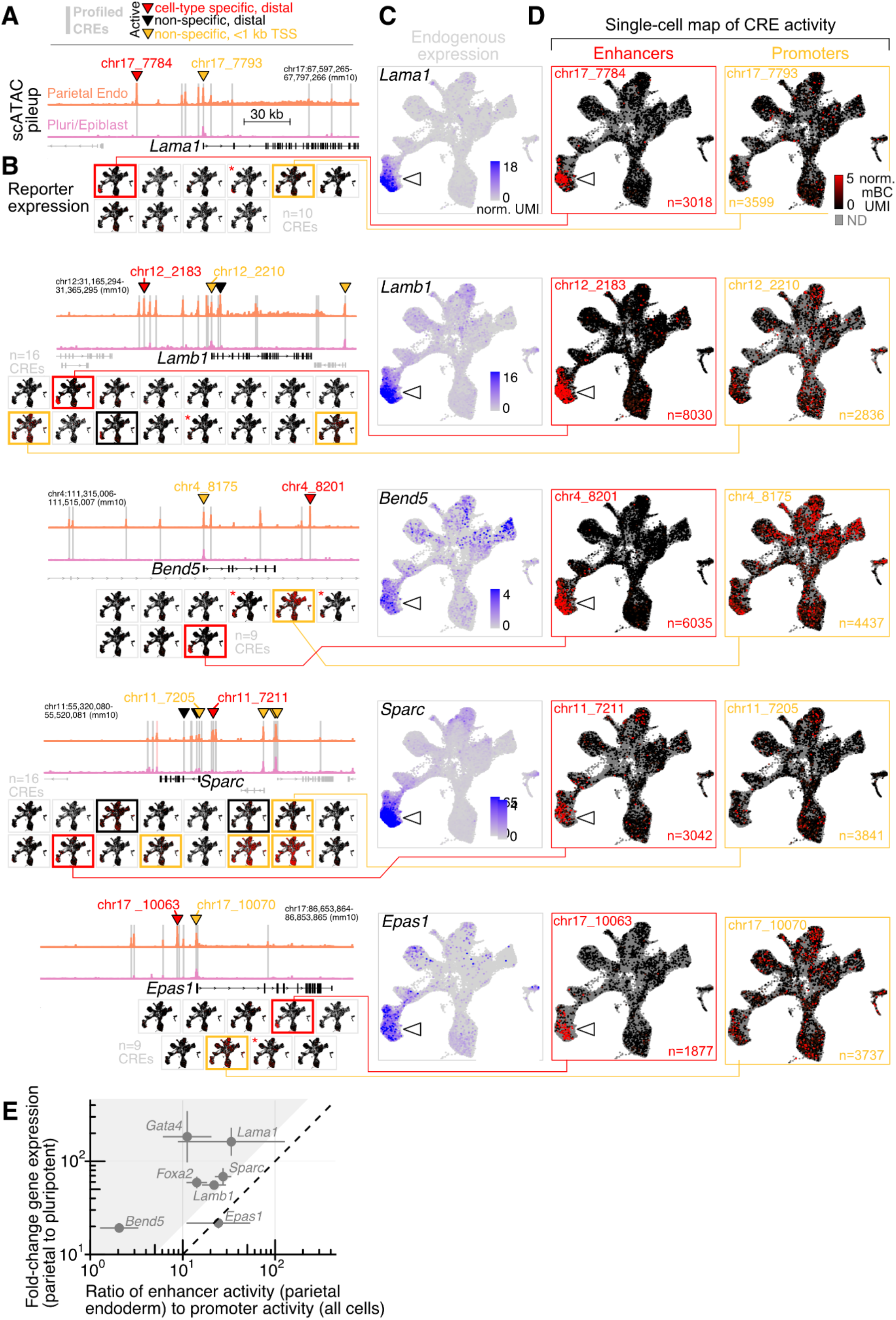
Additional loci with lineage specific distal enhancers. **A-D** Same as **Fig. 4B-E**, but for the additional five loci for which cell-type specific enhancers were identified. Each panel A-D is reproduced across rows for the different loci (top to bottom: *Lama1, Lamb1, Bend5, Sparc, Epas1*). The pink shaded element at the *Sparc* locus (chr11_7186) could not be cloned by PCR due to inability to identify specific primers in the vicinity. **E** Assessing recapitulation of endogenous expression from identified autonomous enhancers. Each point corresponds to one of 7 parietal endoderm genes with putatively associated identified active enhancer and promoter shown in Fig. 4 and panels A-D above (e.g., *Lamb1*: enhancer chr12_2183, promoter chr12_2210; enhancer associations to genes are putative). Endogenous gene induction (y-axis): Fold-change in endogenous gene expression (average in normalized UMI counts) from pluripotent to parietal endoderm. Enhancer induction over promoter baseline (x-axis): enhancer activity in parietal endoderm (reporter level, average normalized mBC UMI parietal endoderm) over mean activity of associated promoter in all cells (reporter level, average normalized mBC UMI). Dashed line is 1:1. Shaded area corresponds to enhancer induction < 0.5×(gene expression). Geometric mean over biological replicates is shown (errorbar: standard deviation of geometric mean).

**Figure S12.**
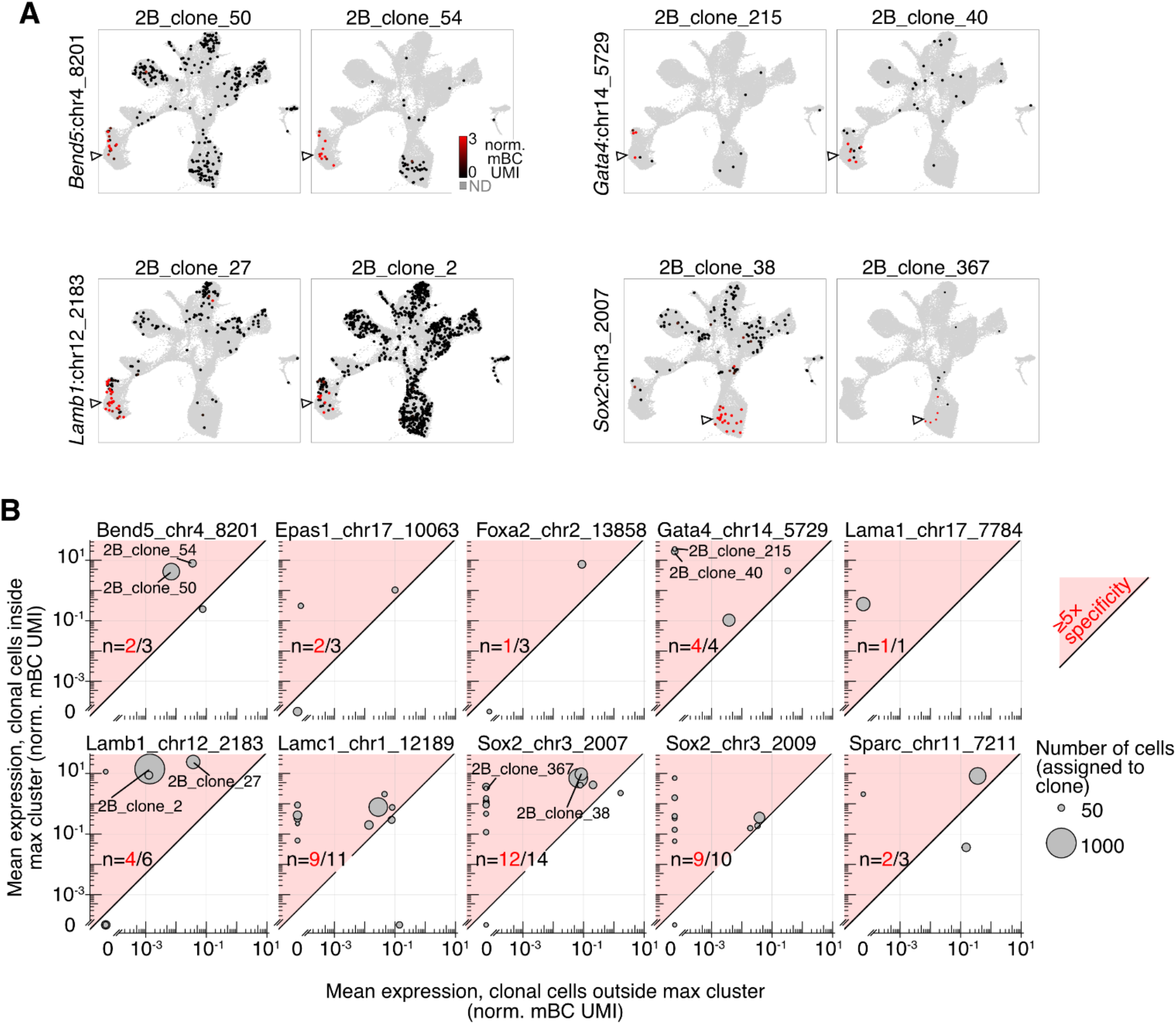
Cell-type specific CRE expression across clones to assess positional integration effects. **A** Example of single-cell map of enhancer activity for cells assigned to high-confidence clones for four CREs (two representative clones per element shown, marked in panel B). Carets indicate the cluster in which expression is expected based on quantification over all cells. Gray points in the background are all other cells not assigned to the clone. **B** Systematic quantification of specificity (activity in expected maximum-expression cluster vs. rest of cells, **Fig. 4A**) across all well-represented clones (5 cells in expected maximum expression cluster(s) and 5 cells in other clusters) for the 10 CREs identified as active and specific. Each clone is represented by a circle, whose area corresponds to the number of cells assigned to it. Clones shown in panel A are indicated. Red shading delineates the region where specificity is in excess of 5-fold. Fractions of clones meeting this criterion for distinct CRE are indicated on each panel. 9/10 CREs have ≥⅔ of their clones with >5-fold specificity.

**Figure S13.**
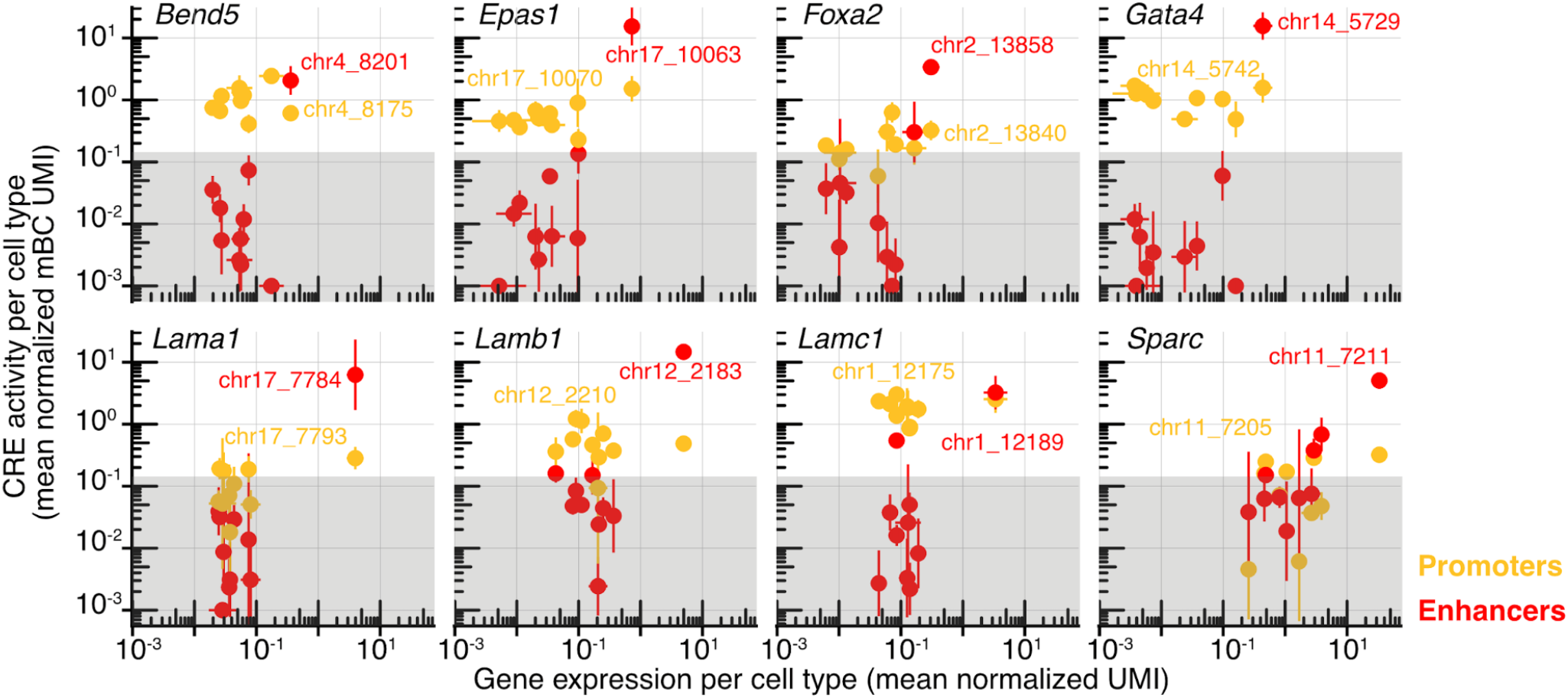
Comparison of per-cell-type gene expression and CRE activity for parietal endoderm loci hits. Quantification of data shown in **Fig. 4D-E** and **S11C-D**. CRE activity (y-axis, mean normalized mBC UMI) compared to putatively associated gene expression (x-axis, mean normalize UMI) stratified per cell type (each point corresponds to average across all cells from fine clusters of **Fig. S5A**, shown is the geometric mean across biological replicates). Each panel corresponds to a locus shown in **Fig. 4, S11** with orange and red points corresponding to activity of the promoter (TSS-proximal) and cell-type specific distal enhancers, respectively. Gray shading marks the limit of detection based on variability of basal controls (no and minimal promoter). Promoters have largely constant expression across cell types, whereas developmental enhancers in some cases have >10^3^ induction in the cognate cell-type (parietal endoderm). Error bars: standard deviation of geometric mean across biological replicates.

**Figure S14.**
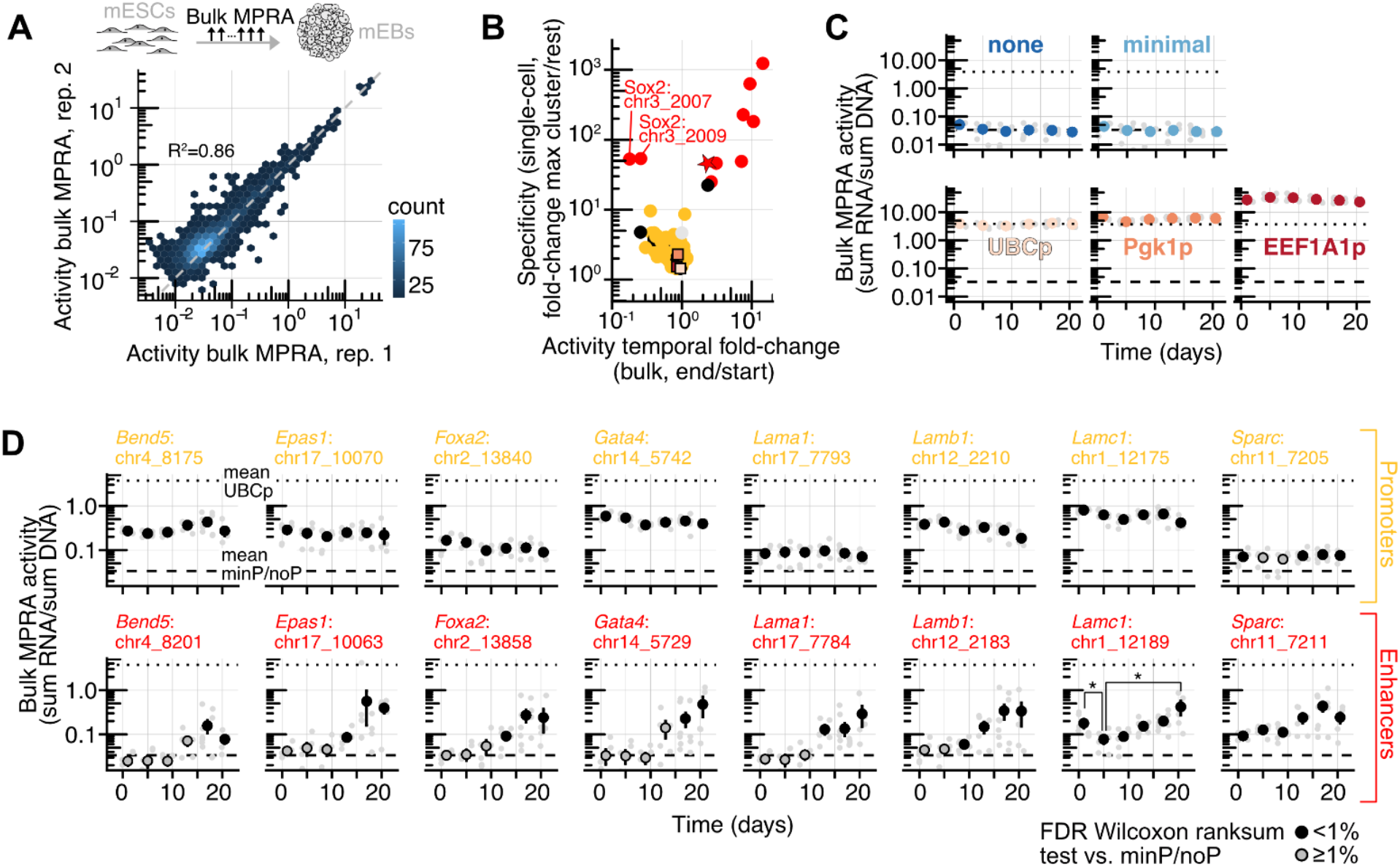
Cell-type-specific CREs are temporally dynamic along mEB differentiation. **A** Reproducibility of bulk MPRA measurement. Comparison of bulk MPRA activity (RNA/DNA ratio of summed 1% winsorized normalized UMI counts) for all CREs in two biological replicates (>10 measured barcodes in both replicates, including exogenous promoters) at all time points (n=2508 comparisons, R^2^ from log-transformed activity). **B** Differentiating EBs were sampled every two days at passage from all replicates, and bulk RNA/DNA MPRA libraries were generated. Fold-change in bulk MPRA activity across time course (mean activity day 20.5 over mean day 1) was compared to the observed specificity of elements as quantified from the scQer end-point quantification (**Fig. 4A**). Elements shown found to be active in either bulk or single-cell assays are shown and coloured according to class (red: cell-type specific, orange: non-specific, <1 kb from TSS, black: non-specific, distal ≥1 kb TSS). The one gray point corresponds to the single element found to be active in bulk but not single-cell assay. Active exogenous promoters (UBCp, Pgk1p, EEF1A1p, panel B) are shown as squares. There is a correspondence between cell-type specificity and temporal change from the bulk assay. Bulk temporal fold-change is 5-10x smaller compared to single cell quantification likely due to bulk assay averaging activity from all cell-types. **C** Activity traces of bulk MPRA time quantification for the exogenous promoters included as internal controls. Small gray points correspond to activity (RNA/DNA ratio of summed 1% winsorized normalized UMI counts) from different replicates/time points. Large black points are the average of two replicates from adjacent time points. Error bars correspond to standard deviation of the mean. Average of basal expression controls (no and minimal promoters) is shown as the dashed line, and the dotted line corresponds to the mean UBC promoter activity (reproduced in panel D for scale). **D** Same as panel B, but for active cell-type-specific enhancers (red) and promoters (shown) from the loci shown in **Fig. 4** and **Fig. S11**. Points are filled when significantly above basal expression controls (ranksum test, B-H corrected, FDR<1%, **Methods**). Promoters (orange) show largely constant expression over time. Enhancers (red) show substantial induction over the time course. Bifunctional enhancer *Lamc1*:chr1_12189 displays initial decrease followed by and increase consistent with its activity in both undifferentiated and differentiated cells (*:p<0.05 Bonferoni corrected ranksum test between day 1 and day 5, and between day 5 and day 20.5).

**Figure S15.**
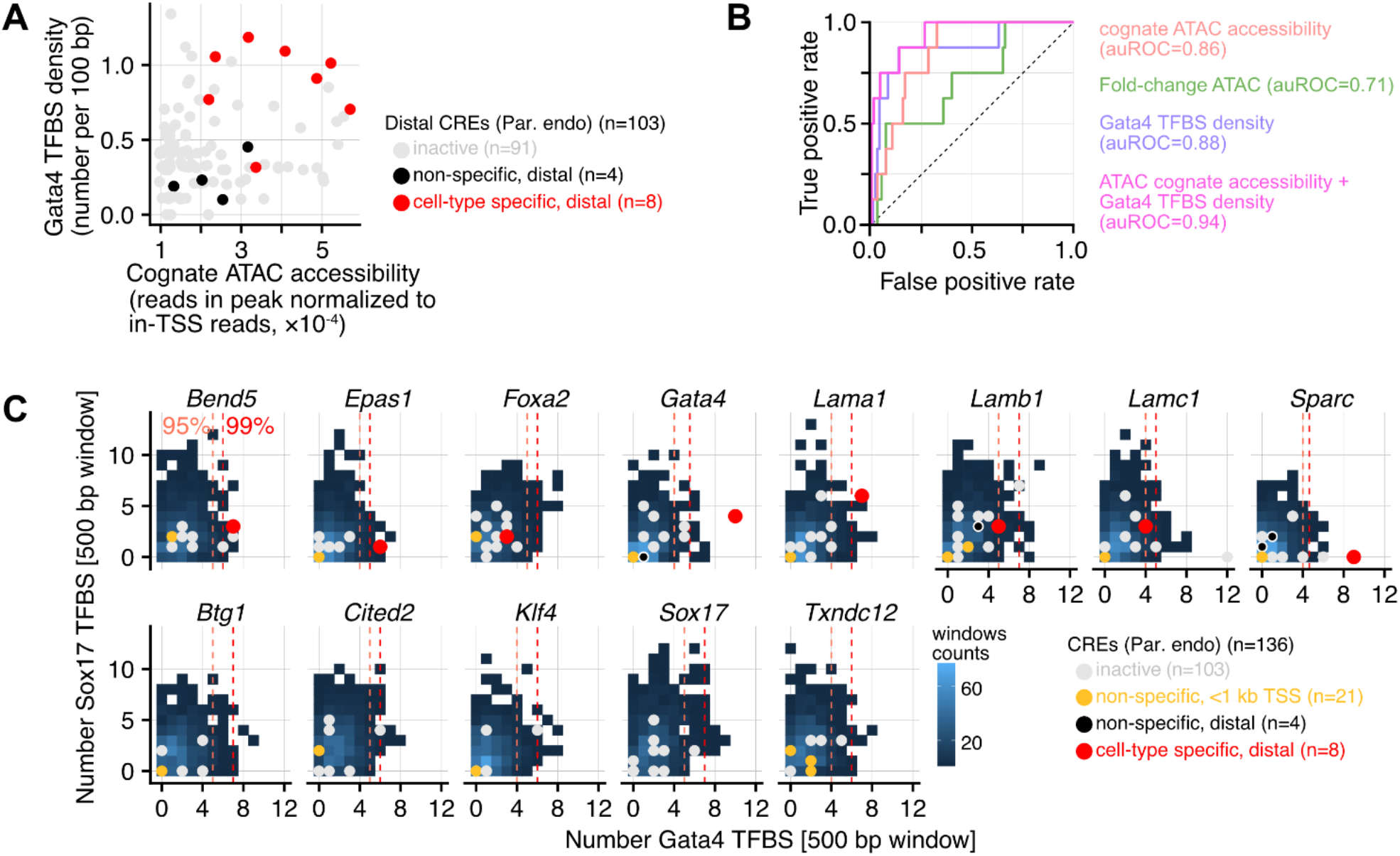
CRE features correlated to cell-type-specific activity. **A** Plot of two features highly enriched for autonomous cell-type specific enhancers: cognate (cell-type corresponding to differential expression of putatively associated gene) ATAC accessibility (x-axis): average in peak reads normalized by reads TSS (in each cell). y-axis: Density of Gata4 transcription factor binding sites per 100 bp (TFBS with affinity relative to the maximum affinity 8-mer >0.4, **Methods**). Red points mark cell-type specific enhancers. Distal (>1 kb TSS) CREs selected from parietal endoderm loci are shown (n=103). B Receiver operating characteristic (ROC) curves for the classification task (specific vs. non-specific/inactive) from different features. Density of Gata4 TFBS, cognate ATAC accessibility, and fold-change in ATAC signal have good predictive value to discriminate functional elements (auROC >0.7). A logistic regression classifier (**Methods**) including only cognate ATAC accessibility and Gata4 TFBS improves performance to auROC=0.94 (precision=0.6 at recall=0.75, not shown). Categories are unbalanced (active=8, inactive=95). **C** Sequence analysis of all 500 bp windows (sliding step 250 bp, excluding any window overlapping with CREs with buffer flank position 500 bp on either sides) for the 13 endoderm-specific developmental loci (±100 kb from TSS of indicated gene). For each genomic sequence window, the number of transcription factor binding sites to Gata4 and Sox17 (affinity relative to the maximum affinity 8-mer >0.4, **Methods**) is recorded. Panels show the two-dimensional distribution of binding sites numbers across all windows, stratified by loci (parietal endoderm elements). The number of binding sites is also determined for tested CREs (coloured points; red: cell-type specific, orange: non-specific, <1 kb from TSS, black: non-specific, distal ≥1 kb TSS; gray: inactive) and overlaid on the distributions for comparisons. Cell-type specific CREs (red points) have an elevated number of Gata4 binding sites compared to other inactive CREs as well as neighboring regions in the loci. Dashed lines mark the 95^th^ and 99th percentile in Gata4 binding site numbers at each locus. 7/8 autonomously active CREs in top 5%, 5/8 in top 1% of number of Gata4 binding sites.

