## Supplementary material for "Multiplex profiling of developmental enhancers with quantitative, single-cell expression reporters": Methods


**1. Benchmarking and optimization: promoter series in human cell lines** 3
1.1. Cloning and subassembly of dual-RNA reporters promoter series 3
1.2. Cell culture, transfection, bottlenecking, and harvesting 4
1.3. Bulk MPRA library preparation 5
1.4. Bulk MPRA data processing and quantification 6
1.5. Single-cell reporter libraries preparation 6
1.6. Single-cell reporter data processing 7
1.6.1. Gene expression libraries 7
1.6.2. mBC libraries 8
1.6.3. oBC libraries 8
1.7. Optimization of reporter RNA capture 9
1.8. Estimating per oBC per cell captured library complexity 9
1.9. Quantification of expression in single-cell assay and comparison to bulk 10
1.10. Estimating the probability to have multiple integration per cell for one plasmid 10
1.11. Clonal cell analysis 11
1.11.1. Clonotype identification 11
1.11.2. Mapping of cells to clonotypes 12
1.11.3. Systematic oBC dropout analysis (precision-recall) 12
1.11.4. Analysis of reporter barcode expression variability across clones 12

**2. Profiling developmental cis-regulatory elements in mouse embryoid bodies** 13
2.1. Cell culture 13
2.1.1. Mouse embryonic stem cells 13
2.1.2. Mouse embryoid bodies induction and maintenance 14
2.2. scATAC-seq on mEBs 14
2.2.1. Experimental method 14
2.2.2. Processing of scATAC-seq data 14
2.3. Prioritization of developmental loci and putative CRE selection 15
2.4. Construction of CRE series dual RNA reporter plasmid library 16
2.4.1. Recloning of oBC-mBC backbone plasmid 16
2.4.2. oBC-mBC subassembly 16
2.4.3. PCR cloning of putative developmental CREs and assembly in dual RNA plasmid 16
2.4.4. oBC-CRE subassembly 17
2.4.5. Final oBC-CRE-mBC triplet table 18
2.5. Experimental details of pooled screen for CRE in mEBs 18
2.5.1. Transfection, cell culture, and bottlenecking 18
2.5.2. End-point processing and single-cell sequencing 19
2.5.3. Construction of EEF1A1p-mCherry transposon plasmid 20
2.5.4. Optimization of high multiplicity of integration with piggyBac in mESCs 20
2.6. Single-cell reporter libraries preparation and sequencing 20
2.7. Single-cell reporter data processing 21
2.7.1. Quality filtering from gene expression libraries 21

2.7.2. mBC and oBC libraries 22
2.7.3. Single-cell quantification of reporter expression 22
2.7.4. Quantification of activity and specificity of CREs and statistical tests 22
2.8. Bulk MPRA (CREs, mEB time series experiment) 23
2.8.1. Library preparation and sequencing 23
2.8.2. Data processing and quantification 24
2.9. Single-cell data integration 25
2.9.1. Integration between scRNA-seq and Pijuan-Sala et al in vivo scRNA-seq 25
2.9.2. Integration between scRNA-seq and scATAC-seq and correlation with in vivo data 25
2.10. Clonal cell analysis 26
2.10.1. Clonotype identification, refinement, cell assignments, basic metrics, and dropout assessment 26
2.10.2. CRE expression pattern across clones 26
2.11. Analysis of features of profiled putative developmental CREs 27

**3. Pol III driven circular vs. linear barcode MPRA experiment** 28
3.1. Cloning of plasmids 28
3.2. Transfection, cell culture, and cell harvesting 28
3.3. Massively parallel reporter assay library generation and sequencing: 29
3.4. Data pre-processing and quantification 30
3.5. Estimating expression levels of oBC per cell per integrated cassette 30
3.6. Evidence of Pol III hU6 transcription not enhancing for Pol II activity 31

### 1. Benchmarking and optimization: promoter series in human cell lines

#### 1.1. Cloning and subassembly of dual-RNA reporters promoter series

To generate the dual-RNA reporter plasmid libraries, we first created a barcoded "cloning dock" plasmid, with restriction sites and homology regions to various cassettes enabling modular addition of 1) Tornado(*37*) RNAs cargos, 2) cis-regulatory element libraries, and 3) reporter mRNAs. To generate the cloning dock, plasmid p001 containing a piggyBac transposon backbone (*84*) was digested with XbaI and HpaI (NEB) and the backbone product purified by agarose gel extraction (Zymoclean Gel DNA recovery kit, Zymo Research). To generate the cloning dock insert, a GFP fragment with barcoded 3' UTR was amplified from plasmid pSGR017 with oJBL315+oJBL316 (all primers and oligos are listed in **Data S1**) and the resulting product gel purified by PAGE. The barcoded 3' UTR was combined with gene block gJBL008 with the piggyBac backbone by isothermal assembly (HiFi NEBuilder, NEB), the resulting plasmid, p022, was electroporated in *E. coli* (NEB, C3020), and the full complexity of the library maintained. Throughout, constructs were confirmed by colony PCR and Sanger sequencing of multiple clones.

We then added a barcode and capture sequence to the Tornado RNA plasmid pAV-U6+27-Tornado-Broccoli plasmid (*37*) (Addgene #124360). The Tornado plasmid was digested with NotI and SacII (NEB) and the backbone purified by agarose gel extraction. A barcoded insert fragment was generated by PCR using the pAV-U6+27-Tornado-Broccoli plasmid as template and primers oJBL220+oJBL291. The barcoded insert was assembled with the purified digested Tornado backbone and gene fragment gJBL007 by isothermal assembly and electroporated in *E. coli* (NEB, C3020), maintaining the full complexity of the library. The resulting plasmid, p019, contained the oBC with capture sequence 1 (CS1) cargo inserted in the Tornado cassette. Plasmids p019 was then digested with BamHI and XhoI (NEB) and p022 with BsbI, with the insert and backbone respectively purified by agarose gel extraction. The components were combined by isothermal assembly to generate plasmid library p025, which was electroporated in *E. coli*, maintaining complexity. Plasmid p025 contains the two barcodes (oBC and mBC) separated by 344 bp and is the starting point to clone scQers (**Fig. S7**).

To construct five libraries (one per promoter in the series, see below), p025 was separately bottlenecked to an estimated 300 clones five separate times, and the oBC and mBC were subassembled from the separate pools. Briefly, amplicons were generated from the bottlenecked p025 as template, and using primers oJBL345 and oJBL337-oJBL341 (indexed primer, one per library). Reactions were carried out in 50 uL volume with 20 ng input plasmid template (25 uL polymerase master mix, 2.5 uL 10 uM oJBL345, 2.5 uL 10 uM indexed primer oJBL337-oJBL341, 0.25 uL 100x SYBr green, water to 50 uL) using Kapa HiFi PCR master mix (Roche) with PCR conditions: 95C 3 min, cycling with 98C 20 seconds, 60C 20 seconds, 72C 30 seconds. Reaction was tracked by qPCR and collected at the inflection point. Amplicons were purified by 1x Ampure.

Libraries were diluted to 2 nM based on the Tapestation D1000 HS quantification, and sequenced on NextSeq 500 with the custom primers: read 1 primer oJBL346 (oBC, 26 cycles), index 1 primer oJBL347 (library index, 6 cycles), read 2 primer oJBL348 (oBC reverse complement, 25 cycles), and index 2 primer oJBL349 (mBC reverse complement, 20 cycles).

Sequencing data was demultiplexed using bcl2fastq. Raw fastq files were processed first by trimming unnecessary cycles from the 3' end (10 cycles from read 1, 5 cycles from read 2, 9 cycles from index 1) using seqtk (https://github.com/lh3/seqtk). Forward and reverse oBC reads were joined and error corrected with PEAR (*85*) (options -v 16 -m 16 -n 16 -t 16). Using custom python and R scripts, assembled oBC reads were combined with mBC reads, and oBC/mBC pairs were counted. The read count distribution displayed a clear bimodal distribution suggesting a saturated library, and oBC-mBC pairs with >500 reads were retained as valid. In order to further restrict the list of oBC-mBC pairs unique across the five bottlenecked libraries, all oBC/mBC pairs were combined, and any pair containing a oBC or mBC appearing more than once (either within a library, or across different libraries) was discarded to avoid mapping conflicts in the analysis of single-cell reporter data (amounting to 24% of high read count pairs), leaving 1122 unique oBC-mBC pairs across the five libraries (number of oBC-mBC pairs per library ranging from 139 to 306, with a median of 205).

Finally, each bottlenecked p025 library described above was digested with BglII, purified by 1x Ampure, digested with EcoRI (NEB), and the resulting backbone was purified by agarose gel extraction. Inserts comprised of various promoters with puromycin cassette and GFP linked by a P2A element were generated as follows. For the human EEF1A1 promoter (including the first intron), minimal promoter and promoterless cassette, primers oJBL254+oJBL314 were used to amplify respective constructs from plasmids pSGR017, pSGR018, and pSGR019 respectively, yielding a promoter puromycin-P2A-GFP fragment. For the human UBC promoter (including the first intron), puromycin-P2A-GFP fragment was obtained by amplifying from pSGR017 with primers oJBL254+oJBL392, and the promoter fragment was amplified from plasmid pB-rtTA with primers oJBL393+oJBL394. For the mouse Pgk1 promoter (no intron), puromycin-P2A-GFP fragment was obtained by amplifying from pSGR017 with primers oJBL254+oJBL392, and the promoter fragment was amplified from plasmid PGK1p-Cys4-pA with primers oJBL395+oJBL396. Promoter sequences are listed in **Data S2**. All fragments were gel purified, combined with their respective digested bottlenecked p025 backbones, and electroporated, resulting in five dual-RNA barcode reporter plasmid libraries, one for each promoter: p029 promoterless (noP), p027 minimal promoter (minP), p042 PGK1, p041 UbC, and p028 EEF1a. Given the a priori subassembly of mBC-oBC pairs for the starting bottlenecked plasmids, and the fact that each library above was assembled separately, each promoter was associated with a list of pairs of oBC and mBC, enabling downstream quantification in a single-cell context (see below).

Plasmid libraries were purified by midiprep (Zymo Research), concentrated by isopropanol precipitation, and pooled at 1:1 ratio by mass. This pooled library of the five promoters was used for both the benchmarking experiment in cell lines (**Fig. 2A**) and was also spiked in the developmental CRE experiment in mESC (**Fig. 3B**).

### 1.2. Cell culture, transfection, bottlenecking, and harvesting

K562 cells were grown in RPMI 1640 medium (ThermoFisher, cat. num. 11875119), supplemented with 10% FBS (Fisher Scientific, Cytiva HyClone™ Fetal Bovine Serum, cat. no. SH3039603) and 1x Penicillin/streptomycin (ThermoFisher, cat. num. 15140122). HepG2 and HEK293T cells were grown in DMEM (ThermoFisher, cat. num. 10313021) with 10% FBS and 1x Penicillin/streptomycin. Cells were kept at 37C and 5% CO2, and passaged every two days (K562, HEK293T) or when cells reached confluency (HepG2, typically every three days). For clonal expansion, we waited for near confluence from 12-well plates (1-2 weeks) before passaging.

All cells were transfected in mid-exponential phase. K562 cells were transfected using MaxCyte electroporation following manufacturer's protocol (1.5 M cells, with 15 ug reporter scQers promoter plasmid mix (see above), 0.5 ug superPiggybac transposase (SBI) in 50 uL volume). Two replicates of 1 M of HepG2 and HEK293 cells were transfected using lipofectamine 2000 (ThermoFisher, cat. no. 11668030, Gibco Opti-MEM cat. no. 31985) with 4 ug of reporter plasmid mix and 0.2 ug of super PiggyBac transposase (SBI). Medium was changed the next day, and cells passaged as usual thereafter. After 5 days, cells were put on puromycin selection (Gibco, cat. no. A1113803, concentration: 2 ug/mL), and grown for an additional 10 days to allow complete dilution of non-integrated plasmids. After >15 days of growth post-transfection, populations from each cell line were bottlenecked to an estimated 250 and 500 starting clones, and expanded to large populations. Notably, HepG2 cells displayed less robust growth at low densities, and required longer time for expansion, suggesting an effectively more severe bottleneck, in line with inferred clonal population properties (fewer final clones, see **Fig. S4C** and **S4E**).

The bulk vs. single-cell quantification experiment (**Fig. 2**) was performed in two replicates. The first replicate (replicate A) with populations bottlenecked at an expected 250 clones, and the second replicate (replicate

B) with populations bottlenecked at an expected 500 clones. For each replicate, at the same time, cells from each line were: 1) harvested separately and methanol fixed for bulk quantification, and 2) prepared as single cell suspension, hand-mixed at an expected 1:1:1 ratio, and profiled for single-cell transcriptomics. Briefly, for the bulk methanol fixation, K562 cells (and HEK293 and HepG2 cells following lifting off plate with 0.05% trypsin) were washed once with ice cold PBS, and resuspended in 80% ice cold methanol, to a concentration of 1 M cells/mL, and placed at -80C until further processing. For single-cell processing, cells were washed twice with PBS+BSA (0.04%) and diluted to 1000 cells/uL. Cell dilutions were mixed at estimated equal proportion and loaded to expected 10k recovered cells total on the 10x Chromium platform following manufacturer's protocol (CG000205 Rev D, Single Cell 3' v3.1 with feature barcoding, 10x Genomics), as one lane per replicate (two lanes total). Replicate B showed some evidence of a partial wetting failure, but otherwise displayed a good emulsion.

#### 1.3. Bulk MPRA library preparation

Genomic DNA was extracted from methanol fixed cells using the DNeasy kit (Qiagen), and RNA was extracted from cells using TRIzol LS (Thermo Fisher), following manufacturer's instructions in both cases. MPRA amplicon libraries from DNA were generated in two steps of PCR amplification with Kapa HiFi (Roche). 0.5-1 ug of genomic DNA input was used. For low-cycle number PCR1, gDNA was mixed with 50 µL 2× Kapa HiFi master mix, 5 µL 10 µM oJBL039, 5 µL 10 µM oJBL358, and water to 100 µL. Cycling parameters: 1 min at 95C, and 4 cycles of: 20 s at 98C, 20 s at 60C, 30 s at 72C, followed by 4C hold. Primer oJBL358 contains 10 random Ns to serve as a pseudo-UMI (hereafter referred to as UMIs for brevity) to correct for PCR jackpotting. Reactions were cleaned up with Ampure XP beads (Beckman Coulter) at 1×, and eluted in 20 µL of 10 mM Tris 8. Illumina adapters and sequencing indices were appended through PCR2, with 4 µL of the eluate from PCR1 taken as input, and 25 µL 2× Kapa HiFi master mix, 0.25 µL 100× SYBr green, 2.5 µL 10 µM oJBL077, 2.5 µL 10 µM indexed primers (oJBL359-oJBL364), and water to 50 µL. Libraries were amplified with tracking by qPCR with: 1 min at 95C, and cycles up to the qPCR inflection point: 20 s at 98C, 20 s at 60C, 30 s at 72C. Libraries were then cleaned up with Ampure XP beads at 1×.

Amplicons libraries for RNA were obtained by first DNase-treating RNA (5 µg RNA, 2 µL TURBO DNase [Thermo Fisher], 2 µL 10× buffer, and water to 20 µL, incubated at 37C for 30 min, cleaned up with RNA clean & concentrator [Zymo Research], and eluted in 11 Tris 7 10 mM). 1 µg of DNase treated RNA was then taken to reverse transcription. Briefly, 2 µL (500 ng/µL) RNA was mixed with 2 µL 1 µM oJBL358, incubated at 65C for 5 min, and placed on ice. 15 µL of reverse transcription master mix was then added (4 µL 5× FS buffer, 1 µL 0.1 M DTT, 1 µL 10 mM dNTP mix, 8 µL water, 1 µL SSIII [Thermo Fisher]), and the reaction incubated at 55C for 60 min, followed by 70C for 15 min. Half of the reverse transcription reaction was then directly amplified for PCR1 (37.5 2× Kapa HiFi master mix, 3.75 µL oJBL039 10 uM, 3.75 µL oJBL077 10 uM, water to 75 µL), with cycling parameters: 1 min at 95C, and 4 cycles of: 20 s at 98C, 20 s at 60C, 30 s at 72C, followed by 4C hold. Reactions were cleaned up with Ampure XP beads (Beckman Coulter) at 1×, and eluted in 20 µL of 10 mM Tris 8. PCR2 proceeded as for libraries prepared from genomic DNA, with oJBL077 and indexing primers (oJBL365, oJBL366, oJBL437-oJBL440), and reactions were stopped at inflexion point from qPCR tracking. Libraries were then cleaned up with Ampure XP beads at 1×.

Final libraries were quantified with Qubit dsDNA HS (Thermo Fisher), diluted to 3 nM, run on TapeStation D1000 HS (Agilent) for final quality assessment, and adjusted to final 2 nM based on the Tapestation quantification. Libraries were pooled, paired end sequenced on NextSeq500 with the following primers and cycle numbers: read1 (mBC forward): 28 cycles, primer oJBL369; index1 (UMI): 19 cycles, primer oJBL435; read2 (mBC reverse): 19 cycles, primer oJBL371; index2 (sample index): 10 cycles, primer oJBL370.

1.4. Bulk MPRA data processing and quantification

Sequencing data was demultiplexed using bcl2fastq. Raw fastq files were processed first by trimming unnecessary cycles from the 3' end (13 cycles from read 1, 4 cycles from read 2, 9 cycles from index 1) using seqtk (https://github.com/lh3/seqtk). Forward and reverse mBC reads were joined and error corrected with PEAR (*85*) (options -v 15 -m 15 -n 15 -t 15). Using custom python and R scripts, successfully assembled barcode reads were combined with UMI reads, mBC-UMI pairs were counted, and the read and UMI counts per mBC determined. The read and UMI counts for the mBC present in the reporter pool (determined *a priori*, see section on reporter cloning and subassembly above) were collected for downstream analysis and comparison to single-cell quantification.

Expression for each mBC from the UMI counts table was computed as follows. First, the total UMI per sample (per cell line and replicate) to the mBC in our list was determined for both RNA and DNA derived libraries. Each mBC UMI count was then normalized by the summed of counts in its respective sample type (DNA and RNA). The normalized RNA UMI count was then divided by the normalized DNA UMI count, to generate the bulk MPRA derived estimate of expression per mBC.

1.5. Single-cell reporter libraries preparation

For single-cell reporters, three libraries are generated: the standard 3' gene expression (GEx) library from 10x, and two custom derived libraries, one for each reporter RNA (oBC and mBC). The latter are obtained from nested PCRs from the amplified cDNA as we detail below.

Briefly, single-cell library preparation proceeded following the manufacturer's protocol (v3.1 manual CG000205 Rev D, 10x Genomics), with some critical modifications listed here. First, one of the replicate's cDNA (replicate B) was split in two equal halves (and brought to same final volume with elution solution 1) after GEM RT cleanup (step 2.1.s) prior to cDNA amplification to allow for a direct comparison the UMIs captured with different enrichment strategy (hereafter replicate B1 and B2). For cDNA amplification, primers specific to the mBC (oSR38) and oBC (oJBL246) reporter transcripts were spiked-in the reaction (similar to TAP-seq (*44*)) at final concentration of 0.5 uM to boost UMI capture for replicates A and B1 (but not for replicate B2, to allow direct comparison with replicate B1). Following cDNA amplification, both the bead and supernatant derived material (steps 2.3Ax and 2.3Bxiv respectively) were saved for downstream processing.

Gene expression libraries for all replicates were prepared following the manufacturer's protocol from 25% of the bead fraction amplified cDNA.

oBC enriched libraries were prepared as follows. For replicate B2 (no primer spiked in), a first outer PCR1 was performed using 25% of the supernatant amplified cDNA with primers oSR40+oJBL246 using Kapa Robust (Roche) and tracking with qPCR until the inflection point (50 uL 2x master mix, 12.5 uL supernatant cDNA, 5 µL 10 µM oJBL246, 5 µL 10 µM oSR40, 0.5 uL SYBr green, and water to 100 µL; run parameters: 3 min at 95C, and cycles 20 s at 95C, 20 s at 60C, 20 s at 72C). Amplicons were cleaned up with 1.75x Ampure XP beads, and 1/10 of the eluate was carried to the inner PCR with the remaining replicates. For replicates A and B1, the outer PCR was performed during the cDNA amplification via the spiked-in primer, and 25% of the supernatant amplified cDNA was taken as input for the next PCR. Semi-nested inner PCR was performed on all samples with primers NextP5_index1 and indexed primers oJBL425-oJBL427, with the same parameters as PCR1 and stopped before the inflection point. Final libraries were purified by 1.5x Ampure XP beads.

As a result of our Pol II reporter construct having a capture sequence (CS2, **Fig. S1A**) downstream of the mBC, reporter mRNAs could be captured from both the poly-dT and CS2 reverse transcription primers on the 10x beads. To systematically compare capture efficiency resulting from the two types of primers, two different libraries were generated (poly-dT captured, and CS2 captured). For poly-dT captured libraries, similar to oBC libraries, we first performed outer PCR on replicate B2 (no spiked-in primers in cDNA amplification) using primers oSR38+oJBL207, using the same PCR conditions as for oBC except for an elongation time of 50 s and an anneal temperature of 65C. 25% of the bead fraction of the purified amplified cDNA was used as template. Following 1x Ampure XP clean up, 10% of the eluate was taken for PCR2. PCR2 was performed on all replicates (directly using 25% of the bead-derived amplified cDNA for replicates A and B1) using primers oJBL324+oJBL495 and the same parameters as PCR1, tracking by qPCR and purifying by 1x Ampure XP beads. A final PCR was performed to index amplicons with primers oJBL076 and indexed primers (oJBL496-oJBL498), and the resulting amplicons purified by 1x Ampure XP beads. The CS2 libraries were prepared entirely analogously to poly-dT captured libraries, except with the following primers: PCR1 for replicate B2 (SR38+SR40), PCR2 all replicates (oJBL529+oSR40), PCR3 all replicates (NextP5_index1+ indexed primers oJBL530-oJBL532).

We note that for both mBC and oBC libraries, semi-nested PCR is necessary to obtain a clean amplicon library (multiple non-specific amplification products were visible following the outer PCR, but a highly specific product was obtained following the semi-nested inner PCR).

All libraries were diluted to 2 nM per the Tapestation D1000 HS reading, pooled, and loaded on a NextSeq 500 for paired-end sequencing the following custom conditions: read 1: 66 cycles (no custom primer); index 1: 10 cycles (primers spiked in: oJBL432, oJBL494); read 2: 76 cycles (primers spiked in: oJBL433, oJBL334). oBC libraries were resequenced to improve saturation of the highly complex oBC libraries following: read 1: 34 cycles (no custom primers); index 1: 10 cycles (primer oJBL432); read 2 38 cycles (primer oJBL433). CS2 mBC libraries were sequenced separately, with: read 1: 30 cycles (no custom primer); index 1: 15 cycles, primer oJBL534; read 2: 18 cycles, primer oJBL334. For mBC and oBC libraries, read 1 provided the cell barcode and UMI, and read 2 the reporter barcode (sequenced with custom primers).

#### 1.6. Single-cell reporter data processing

Four different components are needed to perform reporter quantification using our approach: 1) a triplet map connecting cis-regulatory elements with oBC and mBC sequences, 2) single-cell gene expression UMI counts, 3) single-cell oBC UMI counts, and 4) single-cell mBC UMI counts. For this promoter series experiment, the triplet CRE-oBC-mBC map was described above. We briefly describe below how the count data is obtained for the gene expression and barcoded RNAs. In each case, the output is a count table of the form (cell barcode, gene or barcode, UMI count).

##### *1.6.1. Gene expression libraries*

Data was converted to fastq using bcl2fastq, and fastqs were minimally processed (trimming read 1 to 28 cycles with seqtk, files renamed) to be compatible with cellranger (version 6.0.1, 10x Genomics), which was run using reference GRCh38-2020-A. Each CellRanger count output was processed with Seurat (*86*). Briefly, cell barcodes were filtered to those with >700 gene expression RNA UMIs, and between 2 and 15% mitochondrial UMI fraction. This led to 5787, 4278, and 3834, cell barcodes across the replicates A, B1, and B2. 10x data was normalized, scaled and clustered using standard commands (NormalizeData with LogNormalize method, finding 1000 top variable features with FindVariableFeatures, scaling with ScaleData over all genes, RunPCA and retaining top 50 principal components [PCs] calculated on the identified variable features, FindNeighbors on the top PCs, FindClusters with 0.1 resolution, and RunUMAP with n.neighbors of 20 and using the top PCs as input features). The UMAP revealed three clear clusters (**Fig. 2B**, **Fig. S3A**), hypothesized to correspond to the three cell lines profiled. Replicates B1 and B2 also displayed an intermediate cluster, likely as a result of the lane partial wetting failure, found to share marker genes from the neighboring clusters, which was excluded as plausibly composed of doublets. To confirm the cellular identity of each cluster, in addition to assessment from canonical marker genes (e.g., HBG1/2 in K562, ALB in HepG2), we compared the pseudo-bulked expression (mean across UMI counts for each gene) to bulk expression quantification in the three lines (as assessed from the average of stranded bulk RNA-seq ENCODE(*87*) datasets in K562 and HepG2, and in HEK293), finding unambiguous correspondence of each clusters to a single line (average log-transformed $R^2$=0.72 for matches, vs. 0.39 for non-match).

Following preliminary filtering described above, cell barcodes corresponding to putative doublets were further filtered by two stringent methods. First, each large cluster was further sub-clustered using the same method as above, revealing focal subclusters which shared marker genes from large neighboring clusters, and usually had nearly 2-fold more total RNA UMIs. Cell barcodes contained in these clusters were excluded as likely doublets. Second, scrublet (*88*) was run on the filtered cell barcode set (>700 RNA UMIs, 2 to 15% mitochondrial RNAs), and a doublet score threshold of 0.25 was selected for filtration based on the separation of the bimodal peaks in the simulated score distribution. Cells either belonging to doublet subclusters or having a scrublet doublet score > 0.25 (we observed high concordance between the two approaches) were filtered out. Finally, cells with anomalously high gene expression UMI (>4000) or anomalously high multiplicity of reporter integration (>100, see below), also likely doublets, were removed, leaving 5505 high confidence cells for replicate A (K562: 2184, HEK293T: 2090, HepG2: 1231), 3533 for replicate B1 (K562: 1303, HEK293T: 1238, HepG2: 992), and 3172 for replicate B2 (K562: 1298, HEK293T: 1056, HepG2: 818).

*1.6.2. mBC libraries*

Data was converted to fastq using bcl2fastq, and fastqs were minimally processed (trimming read 1 to 28 cycles and read 2 to 22 cycles with seqtk, files renamed) to be compatible with cellranger (version 6.0.1, 10x Genomics), which was run to perform error correction on cell barcodes. The resulting position sorted bam files were then parsed for the mBC reads as follows using a custom python script: reads aligning to the reference genome or without either corrected cell barcode or UMI (tags CB and UB in the bam file) were discarded. Only reads with the exact expected 7 nt sequence (TCGACAA) downstream of the mBC (positions 16 to 22) were retained. List of all UMIs corresponding to a cell barcode and mBC pair were stored, discarding chimeric UMIs (taken to be UMIs for which the proportion of reads associated to a given mBC vs. all other mBC in the specified cell barcode falls below 0.2). mBC comprised of all Gs (empty read) were discarded. Importantly, the mBC UMI counts were error corrected as follows. For each given mBC and cell barcode, the Hamming distance between all UMIs was calculated, a graph created by connecting UMIs that with a Hamming distance ≤ 1, and the resulting the number of connected components in the graph was taken as the error-corrected UMI count for a given cell barcode-mBC pair. These error corrected UMI counts were taken as the per single-cell quantification of the reporter mRNA expression (see below for a normalization strategy to correct for technical factors). Given that cell barcodes derived from capture sequence vs. poly-dT reverse transcription primer are different on the 10x Genomics beads (bases 8 and 9 reverse complemented) on the same bead (and not error corrected by cellranger in our application), we converted the CS2 cell barcodes to their poly-dT counterparts to enable matching across the different libraries.

*1.6.3. oBC libraries*

oBC libraries were processed in an entirely analogous way to the strategy for mBC described above, with the following modifications: two sequencing runs were combined in a single fastq prior to processing, read 2 were trimmed to 23 cycles, and only reads with the GCTTTAA (constant region after the oBC) at positions 17 to 23 were retained. The number of UMIs per oBC per cell barcode was also taken as the error corrected (1 Hamming distance) count and our measure of oBC expression in single cells (see below for a normalization strategy to correct for gene expression UMIs). Similarly to the CS2 mBC data above, we again converted the CS1 cell barcode to poly-dT cell barcodes.

1.7. Optimization of reporter RNA capture

Two experiments were performed to quantitatively characterize UMI capture in our system. First, as described above, one of the sample (replicate B) cDNA was split in two prior to amplification. This enabled a direct comparison of the number of UMIs captured with vs. without the addition of reporter specific primers during cDNA amplification (as opposed to relying on the template switching oligo). For both oBC and mBC, we compared the UMI counts across cell barcode-mBC pairs (valid cell barcode from gene expression data, valid reporter barcode from subassembly) in replicate B1 (with spike in primers in cDNA amplification) and B2 (without spike in primers in cDNA amplification), **Fig. S1D**. For mBC, we found a median 2.0x increase in UMI counts (for mBC/cell barcode pairs with 4 or more UMIs in both replicates, **Fig. S1D**) in replicate B1 compared to B2, suggesting increased captured resulting from spike-in (both replicates had number of reads per UMI much larger than 1, and in fact larger for replicate B2 [median reads/umi =17.9] compared to B1 [median reads/UMI=6.8], such that this difference cannot be attributed to increase sequencing coverage for replicate B2), consistently with the range previously reported in TAP-seq(*44*). A similar increase (2.1x increase from replicate B1 to B2) in mBC UMI count was seen also for the CS2 captured mRNAs (not shown). Performing the same analysis for oBC led to a much larger boost in the number of UMIs captured (45x increase, also not attributable to high coverage of replicate B1, **Fig. S1E**) as a result of the spiked-in primer. This larger difference was expected from the circular nature of barcode: given the absence of 5' end from which template switching can occur from circular RNAs, the initial cDNA amplification (primed from the template switching oligo) effectively could not happen except from the linear oBC intermediates (expected to represent a minor fraction) in replicate B2.

In addition, we tested which of poly-dT vs. capture sequence derived primers captured more reporter mRNA. Our reporter cassette (**Fig. S1A**) harbors capture sequence 2 (CS2) downstream of the barcode, enabling a direct comparison, for the same reporter in the same cell, of the different number of UMIs captured from the two different RT primers. Comparing across valid cell barcodes and mBC pairs with at least one UMI captured from both poly-dT and CS2, we found a median of 15.7x more UMIs captured from poly-dT primers, likely a direct reflection of the higher stoichiometry of these primers on the 10x Genomics beads (**Fig. S1F**). As such, in all mRNA expression quantifications, we use the poly-dT captured number of mBC UMIs. In addition, we note that using artificial poly-A sequence in place of CS1 on the oBC would likely result in a similar boost in capture and complexity from this system (RT primer saturation should not be a problem given successful overloading of fluidic emulsions without observed decrease in capture efficiency (*89*)).

1.8. Estimating per oBC per cell captured library complexity

The single-cell oBC libraries were highly complex and as result were not sequenced to saturation. UMI count distributions shown in **Fig. 2C**, **S3B** (and similarly in mEB, **S8F**) were therefore not a measure of the full complexity of the libraries (median duplication rate, i.e., read counts over UMI counts minus 1, of 8.3% and 8.8% respectively for oBCs in the high count mode [≥12 UMIs]). To estimate the total complexity for each oBC in each cell, we used the maximum likelihood estimator from the zero-truncated Poisson distribution (*90*), i.e., if for a given oBC with a given cell barcode x:=(reads counts)/(UMI counts), and $\lambda$:=(read counts)/(oBC complexity), then $x = \lambda(1\text{-}\exp(-\lambda))^{-1}$. Inverting the relation for each cell barcode and oBC pair in high count mode (provided reads counts is not equal to UMI counts), we find a median complexity of 3652 (K562), 2306 (HEK293T), and 1675 (HepG2) UMI per oBC per cell barcode (replicate A). As expected from splitting the cDNA in two, the estimated complexity was essentially halved for replicated B1 (1731 in K562, 1217 in HEK293T, 850 in HepG2).

1.9. Quantification of expression in single-cell assay and comparison to bulk

To quantify reporter expression via our single-cell experiment, we first determined the set of valid oBC (present in our oBC-promoter-mBC subassembly table generated *a priori*) detected in each cell. As a tradeoff between specificity and sensitivity (see clonotype precision-recall analysis below), we selected a threshold of ≥12 UMI (**Fig. 2C**) to deem a oBC as present for a given cell barcode. The UMI counts for valid mBC cell-barcode pairs were then joined to the detected oBC in all valid cell barcodes by using the predetermined oBC-mBC (uniquely matchable) association table. In cell barcode/oBC combinations for which there were no detected mBC UMI, a value of 0 was taken (detection of reporter integration from oBC, but no captured reporter mRNA). Importantly: while not detected, given our dual RNA strategy, this represented a "true" zero and contributes to our measurement of expression. mBC UMI counts were normalized by the number of gene expression UMI (from the full transcriptome GEx libraries) detected in each cell, i.e., (mBC UMI)/(GEx UMI)*mean(GEx UMI), where the scaling with the mean gene expression UMI across all cells served to maintain an intuitive unit in the data. Normalization by simple scaling by gene expression UMI was performed as the mBC UMI counts were correlated ($R^2$ of log-transformed values=0.09) with gene expression UMI with a slope close to 1 (least square fit on log-transformed data, slope: 0.93). We find in both our comparison to bulk data and our clonal analysis (see below) that direct normalization of mBC by GEx slightly improves the precision of the expression measurement. To quantify single-cell expression for each mBC (**Fig. 2E**), we then directly averaged the normalized mBC UMI counts across all cells with a detected associated oBC.

The averaged normalized mBC UMI described above was directly compared to the bulk expression quantification (from bulk MPRA), **Fig. 2E** and **S3D**. In these analyses, we only include well-represented barcodes in the comparisons to focus attention on technical noise resulting from the two methods and not noise from sparse sampling of rare barcodes (mBC with 5 or more cells with oBC detected integrations, at least 1 mBC UMI captured across all integrations, and at least 100 DNA UMI from the bulk quantification).

For quantification without conditioning on oBC detection (**Fig. S3G**), the average normalized mBC UMI across all cells with any captured counts was taken. Including an additional step to filter possible chimeric amplicons (removing events for which the number of reads equaled the number of UMIs, unlikely in a saturated library) did not substantially improve performance without oBC detection.

In addition to the accuracy comparison to the bulk quantification, we also directly assessed the number of incorrectly detected mBC (mBC UMI count >0, but not detected as determined by absence of the associated oBC (<12 oBC UMI) in the same cell) for the different promoters. We found the following proportions of valid (oBC matched) mBC detection events (mean proportion from replicates A and B1): no promoter: 60%, minimal promoter: 45.9%, UBCp: 51.4%, Pgk1p: 40.4%, EEF1A1p 10.5%. Spurious detections thus constituted a substantial, and sometimes dominant, proportion of events in all cases.

1.10. Estimating the probability to have multiple integration per cell for one plasmid

In order to estimate the probability that a unique oBC-promoter-mBC plasmid ends up integrating multiple times in a given cell, we simulated the genomic integration process by drawing multiplicity of integration from the empirically observed distribution (assuming no multi-integration events), and drawing oBC-promoter-mBC combination with replacement with frequency taken as the quantified proportion in our pool (as assessed during the subassembly stage by sequencing the barcodes). We find the probability of multiple integration, regardless of barcode, to be less than 5% in both replicates. This is somewhat higher than expected from the multi-integration probability from a scenario with fixed number of integration sampling exactly equiprobable barcodes (equal <2% under these circumstances). Spread in both MOI (here 2 to 8 integrants per cell interquartile range) and non-even barcode representation (1122 unique oBC-promoter-mBC triplets in our pool, with $10^{th}$ to $90^{th}$ percentile of representation in the pool spanning a 10-fold range: 0.00018 to 0.0017), thus contribute to modestly inflate the likelihood of a multi-integration event, which we nevertheless expect to be rare (<5%) from this empirically derived estimate.

### 1.11. Clonal cell analysis

Identifying clonal cells harboring multiple genetic payloads from single-cell data is a non-trivial computational problem, even with high signal to noise ratio, in part due to doublets and barcodes multiply represented across different clones. After assessments of existing approaches (*47*, *48*, *91*) and our own attempt (iterative clustering in high dimensional PCA space from oBC expression), we settled on a modification of the heuristic put forth by Wang and colleagues (*48*), based on one-sided Fisher's exact test, followed by our own addition of custom quality filtering. Details are provided below.

#### *1.11.1. Clonotype identification*

We identified high-confidence integration genotypes (hereafter clonotypes) in a two step procedure. First, a raw clonotype identification, followed by a refinement step.

As a first pass identification of clonotypes, we followed (*48*) and looped through cells (considering cell barcodes assigned to different cell lines from their transcriptome and different replicates separately), assembling a list of clonotypes. Specifically, for each cell, the list of detected oBC was extracted (≥12 oBC UMI, see below for justification for threshold in addition to corresponding to the minimum of the UMI count distribution, **Fig. 2C** and **S3B**). The list of barcodes from the cell was then compared to the oBCs detected in all other stored clonotypes via a one-sided Fisher's exact test, with contingency table given by (number of oBC in cell and clonotype, number of oBC in clonotype but not in cell; number of oBC in cell not in clonotype, number of oBC from library neither in the clonotype nor in the cell). The test serves to assess the probability that random sampling of oBC leads to as much overlap between cell and clone as observed. A 5% Bonferonni corrected ($0.05/(n_{cells}^2/2)$) p-value was used as a threshold to determine whether a cell was a likely member of a clonotype or not. If oBC from the cell did not overlap significantly from any stored clonotypes, the cell was taken as the representative of a new clonotype. If overlap with multiple existing clonotypes was identified, the cell was marked as a likely doublet. Given the number of cells and barcodes in our experiment (promoter series), *bona fide* clones with only two reporters integrated did not meet the stringency threshold of our test and were thus excluded *de facto*. We note that this heuristic takes the set of oBCs detected from the first representative of a clonotype as the set for all cells (no aggregative correction applied), and as such the resulting cell assignments and clonotypes depend on the order at which cells are considered in the loop. To address this, we implemented an additional downstream filtering step, and returned to the problem of assigning cells to clonotypes after the final list of high-confidence clonotypes was determined.

To refine the raw list of clonotypes identified above, we first exclude clonotypes assigned to a single cell. Then, for a given clonotype, we obtained the union of all detected oBCs ≥12 UMI (clonotype oBCs) from cells assigned to that clonotype. For each of these clonotype oBC, the fraction of cells assigned to the clonotype with detection of that oBC was then determined. We then determined the number of clonotype oBC which were detected between 25% and 75% of cells within the clonotype ($n_{25\% \text{ to } 75\%}$), and the number of clonotype oBC detected in more than 75% of the same cells ($n_{>75\%}$). We also stored the maximum fraction of cells with detection of any one of the oBC within a clonotype (max_frac_detect). We found these quantities to be useful to filter out likely doublets and clonotypes with too much barcode overlap from valid and easily distinguishable clonotypes. We retained clonotypes for which $n_{>75\%}>2\ n_{25\% \text{ to } 75\%}$ and max_frac_detect>0.9. The list of oBC corresponding to a clonotype was then taken as those detected in >50% of cells assigned to that clonotype. Finally, completely nested clonotypes (ones whose set of oBC was a strict subset of another) were eliminated.

*1.11.2. Mapping of cells to clonotypes*

Using the list of clonotypes and associated oBC (described above), we returned to the complete dataset to assign cells to clonotype (thereby avoiding the issue of cell ordering affecting the outcome). Specifically, we obtained the list of detected oBCs (≥12 oBC UMI) in each individual cell. We then computed two quantities across all clonotypes, 1) $f_1$:= fraction of oBCs detected in the cell of interest also present in the clonotype, and 2) $f_2$:= fraction of clonotype oBCs detected in the cell of interest. In words, $f_1$ tracks possible additional barcodes detected in the cell not associated with the clonotype (e.g., doublets), while $f_2$ monitors possible dropouts. For each cell, the top clonotype was taken as the one with the largest $f_1$, and the associated $f_2$ was also retained. Cells were assigned the status of a high confidence singlet if $f_1>0.975$ and $f_2>0.5$. Hence, we stringently filter out possible doublet (require high $f_1$), but remain loose on possible dropouts (allow for low $f_2$, compared to performance, see below). Cells with $f_2\leq0.5$ were considered missed clonotypes.

Through this procedure, we obtained a high proportion of cells assigned as singlets to high-confidence clonotypes (replicate A: K562 67%, HEK293T 66%, HepG2 75%; replicate B1: K562 61%, HEK293T 58%, HepG2 55%), and substantial fraction of the non-singlet cells had an MOI of 2 or lower (replicate A: K562 82%, HEK293T 63%, HepG2 61%; replicate B1: K562 61%, HEK293T 54%, HepG2 30%). Hence a high proportion of missed clonotypes came from low MOI cells and the high stringency of our p-value threshold. The number of clones was somewhat lower than estimated going in the bottleneck, possibly as a result of clonal competition. HepG2 in particular displayed a more severely bottlenecked population (**Fig. S4C**, **S4E**), in line with the longer time necessary for those populations to expand (slow growth was observed at the low plating density). Final clonotypes with cell assignments are listed in **Data S7**.

Final assignments (with clonotypes with 3 or more cells assigned) were displayed on oBC expression space UMAP (**Fig. S4C**, **S4E**) using Seurat (Normalization method "RC", PCA run on variable features with 100 principal components, UMAP with n.neighbors=10 on top 50 PCs).

*1.11.3. Systematic oBC dropout analysis (precision-recall)*

The high-confidence clonotypes identified through the consensus of co-detected barcodes served as an approximate ground truth to systematically assess the detectability of oBC in our assay. Specifically, for all clonotypes (with 3 or more cells assigned) and singlet-assigned cells as described above, we computed for different oBC UMI count detection threshold the number of true positives (TP:= number of oBC detected in cell also in the clonotype), false positives (FP:= number oBC detected in the cell not present in the clonotype), and false negatives (FN:= number of oBC in the clonotype not detected in the cell). At each oBC threshold, the false discovery rate was taken as FDR:=sum(FP)/(sum(FP)+sum(TP)), and the false negative rate FNR:=sum(FN)/(sum(FN)+sum(TP)), where the sums are over all cells and clonotypes. Results stratified by cell lines are shown in **Fig. 2H** (stratified by replicates: **Fig. S4A-B**). Direct representative oBC count distributions are shown in the count matrices to two typical clones shown in **Fig. S4D** and **S4F**. In order to prioritize stringency, we selected a UMI of 12 as threshold for the expression analysis presented throughout (different threshold for experiment in embryoid bodies, see below). We note that given the loose stringency of our threshold for assignment (tolerating cells with up to 50% dropout in oBC), this analysis should be relatively unbiased given that the FNR is in the few percent range at the threshold of 12 oBC UMI.

*1.11.4. Analysis of reporter barcode expression variability across clones*

In addition to providing assessment of oBC dropout, clonal cells present an opportunity to measure variability in the number of captured reporter mRNAs, while controlling for possible positional effects. For each singlet-assigned cells to high confidence clonotypes, the count distribution of oBC UMI and mBC UMI (GEx normalized) corresponding to the integrated reporter cassettes was obtained (e.g., **Fig. S4G-H** for two example clones). The mean across cells assigned to clonotypes and standard deviation in these quantities was determined (analysis restricted to clones with >4 cells assigned to allow for a robust assessment of the standard deviation). The coefficient of variation (standard deviation over mean) was displayed as a function of the mean (**Fig. S4I**), showing scaling close to the limit set by Poisson counting even for some of the highly expressed promoters, and typically much lower than one. This provides direct evidence that when controlling for positional effects and conditioning on presence of the reporter by orthogonal means (here with oBC detection), single-cell measurements can be highly precise.

To assess the proportion of variance attributable to positional effects vs. other technical and biological factors, we used the law of total variance to decompose in mBC UMI variability. For each separate cell line and promoter, we computed the variance of the mean mBC UMI per clone-reporter pair (explained variance) and the mean variance of mBC UMI across clones-reporter pairs (unexplained variance). We find the proportion of unexplained variance to be (average of replicates A and B1): EEF1A1p K562=0.46; HEK293T=0.37, HepG2=0.37; Pgk1p K562=0.44, HEK293T=0.73, HepG2=0.60; UBCp K562=0.33, HEK293T=0.41, HepG2=0.55. The reporter values in the main text are the average over the cell lines.

Variation of the mean expression for the different promoters across clonotypes also provided estimates for the magnitude of genomic context positional effects. We found restricted variability for most promoters, although with evidence for cell-type specific differences (interquartile fold-change range, for all promoters listed from K562, HEK293T, and HepG2: UBCp: 2.1, 2.2, 2.4; Pgk1p: 5.9, 4.8, 7.2; EEF1A1p: 1.5, 1.5, 4.1), somewhat smaller than the positional effects observed from the Pgk1 promoter in mES cells (*50*) which had an observed fold-change interquartile of ≈8, suggesting that positional variegation arising from local the epigenetic environment might be partially mitigated by presence of insulators (core cHS4 (*43*)) in our construct (**Fig. S1A**).

Clonal analysis also allowed us to compare the distribution of CV across clones for the raw UMI counts, and the GEx normalized UMI counts. We found small but consistent decreases in variability both for oBC (median GEx normalized CV/raw CV = 0.80) and some promoters (UBCp and EEF1A1p: median GEx normalized CV/raw CV=0.85, no difference for, the no promoter, minimal, and Pgk1p promoters), justifying our use of this normalization in our quantification of reporter mRNA.

### 2. Profiling developmental *cis*-regulatory elements in mouse embryoid bodies

#### 2.1. Cell culture

##### *2.1.1. Mouse embryonic stem cells*

A low-passage number monoclonal male BL6 (WD44) mouse embryonic stem cell line stably expressing dCas9-BFP-KRAB was used. Cells were grown on gelatin (0.2%) (Sigma, cat. No. G1890) coated plates and cultured in DMEM (ThermoFisher, cat. num. 10313021) supplemented with 15% FBS (Biowest, Premium bovine serum, cat. no. S1620), 1x MEM non-essential amino-acids (ThermoFisher, cat. no. 11140050), 1x Glutamax (ThermoFisher, cat. no. 35050061), $10^{-5}$ beta-mercaptoethanol, and $10^{-4}$ leukemia inhibitory factor (Sigma-Aldrich, ESGRO Recombinant Mouse LIF Protein ESG1107), hereafter referred to serum+LIF medium were necessary, with daily medium changes (aspirate medium, replace with pre-warmed medium), and transfer every two days (aspirate medium, wash with PBS [without $Ca^{2+}$, $Mg^{2+}$], add 2.5 mL [for 10 cm plate] 0.05% trypsin , incubate 2 minutes at 37C, deactivate trypsin and triturate with 10 mL pre-warmed medium, spin down 5 min at 300g, aspirate supernatant, resuspend in pre-warmed medium, and transfer to new gelatinized plate).

*2.1.2. Mouse embryoid bodies induction and maintenance*

Exponentially growing mESCs are lifted from the plate (aspirate serum+LIF medium, wash with PBS, add 2.5 mL [for 10 cm plate] 0.05% trypsin , incubate 2 minutes at 37C, deactivate trypsin and triturate to a single-cell suspension with 10 mL pre-warmed medium). Cells are then counted and spun down (5 min at 300 g). Supernatant is aspirated and cells are resuspended to 2 M/mL in CA medium (medium for EB induction: DMEM, 10% FBS, 1x MEM non-essential amino-acids, 1x Glutamax, $10^{-5}$ beta-mercaptoethanol). Cells are counted again, and density adjusted to 1 M/mL with CA medium. 3 mL (3 M cells) are added to 12 mL of CA medium in 10 cm plates (suspension plates: non gelatinized, non adherent). One the next day, plates are gently agitated to promote cell aggregation. Following induction, embryoid bodies (mEBs) are passaged every two days (no daily medium change). mEBs are collected using a serological pipette and transferred to a 50 mL conical tube (typically three plates are pooled). Leftover mEBs on plates are recovered by a CA medium wash and pooled in the conical tube. mEBs are left to settle (initially up to 15-20 min, faster as the mEBs grow in size). Once mEBs have settled, medium is aspirated from the top, carefully avoiding disturbing the loose pellet. Fresh, pre-warmed, CA medium is then added to 15 mL/plate and mEBs redistributed to plates.

### 2.2. scATAC-seq on mEBs

*2.2.1. Experimental method*

Single-nuclei preparation for scATAC-seq were prepared from day 21 mEB as follows: At day 21 of mEB cultures, mEBs (two 10 cm suspension plates) are collected into a 50 mL conical tube and washed 2x with 1x PBS (without $Ca^{2+}$, $Mg^{2+}$). After consecutive PBS washes, mEBs are treated with 1.5mL of 0.25% trypsin and incubated in 37C bath with gentle agitation (steady concentric swirls in 50mL conical tubes) for 3 minutes. For further dissociation, mEBs are then gently triturated 10 times with a P1000 pipette and again incubated at 37C for 3 minutes with gentle agitation. After second incubation, mEBs are gently triturated 10 times with a P1000 pipette. Trypsin digestion is inactivated with CA medium and cells filtered to single-cell suspension through a 100 um filter into a new 50 mL conical. Single cell suspension was counted, and cells spun down at 300 g for 5 minutes. After removing supernatant, wash 1x with 1 mL of 1x PBS + 0.04% BSA and gently pipette mix 5x. Transfer to a 1.5 mL tube and spin at 300g for 5 min at 4C. Wash again with 1mL of 1x PBS + 0.04% BSA. Again, spin at 300 g for 5 min at 4C, and proceeded with 10x Genomics "Nuclei isolation for Single Cell ATAC Sequencing" protocol (V1), with two biological replicates (different mEB differentiation) each with two lanes of 10x (four reactions total).

*2.2.2. Processing of scATAC-seq data*

Fastq files were generated by running makefastq. Fastq files were then processed to fragment files using 10x Genomics cellranger count (cellranger-atac-cs version 1.2.0, reference = refdata-cellranger-atac-mm10-1.2.0), which were processed through the ArchR pipeline (*92*). Arrow files were created with function createArrowFiles (minTSS=4, minFrags=1000). Two different double scores were computed with function addDoubletScores (LSI based: k=10, knnMethod="LSI", LSIMethod=1); UMAP based: k=10, LSImet=1, UMAPparam: n_neighbors=40, min_dist=0.4, metric="euclidiean"). In addition, we used AMULET (*93*) (from its v1.0-beta version, running function ATACDoubletDetector.py and adding as problematic region a union of the ENCODE excluded list, segmental duplications, simple repeats, repeat masker, and microsatellites from mm10 obtained from UCSC). Nuclei were then filtered for: TSS enrichment >8 and >1995 fragment counts. Following dimensional reduction (addIterativeLSI: useMatrix = "TileMatrix", name = "IterativeLSI", iterations = 2, varFeatures = 100000, dimsToUse = 1:30), and clustering (AddClusters: reducedDims = "IterativeLSI", method = "Seurat", name = "Clusters", resolution = 0.5), doublets were stringently removed by inspecting distribution of fragment counts, doublet scores (ArchR derived), and AMULET doublet scores per clusters. All nuclei from clusters with anomalously high doublet scores across metrics were removed. In addition, individual nuclei with either >17782 fragment counts, LSI doublet score > 0, UMAP doublet score > 0, or AMULET score > 0.3 (thresholds assessed from the distribution of anomalous doublet clusters) were filtered out as likely doublets. In the end, 31% of nuclei were removed with these filters, leaving 46408 nuclei passing quality filters (20329 nuclei from replicate 1, 26079 nuclei from replicate 2).

The resulting filtered nuclei were dimensionally reduced and clustered (same parameters as above), leading to 9 clusters with >200 nuclei. Clusters with highly correlated accessibility (determined from pseudobulk averaging over cells in cluster) over all peaks ($R^2$ on log-transformed accessibility >0.55) and proximal in the low-dimensional projection were merged. The intermediate endoderm cluster (connecting the visceral and parietal clusters) was kept separate to avoid diluting the signal from the two otherwise well-delineated extraembryonic endoderm clusters. The final 7 clusters are depicted in **Fig. S5E** (see later section for integration/annotation). scATAC pseudobulk pileup traces (e.g., **Fig. 3A**, **4B**) were generated by first the normalized data using ArchR's function groupRegionSumArrows. A subset of all cell-type pseudboulks was shown due to space limitations in figures.

2.3. Prioritization of developmental loci and putative CRE selection

In order to select regulatory elements possibly implicated in control of gene expression in our system, we prioritized loci on the basis of a number of criteria. First, we identified highly differentially expressed genes within neuroectoderm, endoderm, mesoderm, and pluripotent clusters from mEBs (SGR and SD, unpublished data) using Seurat FindMarkers function, retaining genes with at least 25% detected expression in the respective clusters, and either a fold-change in expression >1.6x or a fold-change in fraction of cells detected with expression >2. We then identified all peaks from the scATAC data with score (as generated by ArchR) >20 within 100 kb of the TSS of each gene. The resulting gene-peak data table was then augmented with information about the ATAC peaks (accessibility in cell-type cognate to the differential expression, fold-change in accessibility, average phyloP score(*69*), distance to the nearest gene, overlap with ccRE (*11*), orthology to a reciprocal human ccRE). Genes were retained for further assessment if their ±100 kb neighborhood included 3 or more highly accessible and differentially accessible peaks (top 90$^{th}$ percentile) in the cell-type cognate to the differential expression. In addition, loci harboring one or more non-exonic peaks with evidence of conservation (average phyloP>0.75 or presence of orthologous human ccRE) and within 50 kb of a very highly differentially expressed genes (>3.8 fold-change in expression or >7.7 fold-change in fraction of cells with detected expression) were retained. Finally, loci with one or more non-exonic peaks with either: 1) strong conservation (>2 average phyloP score) and high differential accessibility (>15x fold change), or 2) evidence of conservation (average phyloP>0.75 or presence of orthologous human ccRE) and very high differential accessibility (>15x fold change), were retained. Filtering on these different criteria led to a list of 89 loci, which were manually evaluated. To arrive at our final list of 22 loci (**Data S3**), genes were ranked by the number of peaks satisfying the above criteria (conservation and differential activity), and examples from parietal endoderm, neuroectoderm, and mesoderm cell-types with overall low gene density (to avoid the possible complication of neighboring gene regulation) were selected.

Following loci prioritization, the final set of putative CREs selected was any peak within 100 kb of the annotated differentially expressed genes above $>9.4\times10^{-3}$ accessibility (normalized by TSS reads, average in all cells annotated to differential-expression-cognate cell-type) reproducibly in both scATAC replicates, leading to 206 regions. A strongly differentially accessible peak 2 kb upstream of the gene *Tubb2b* TSS (*Tubb2b*:ch13_2580) which had fortuitously not passed our thresholding criteria was included. We finally added the 4 constituents of the core Sox2 control region as enhancers of interest to include, for a total of 211 elements. Robust primers for PCR-cloning could be designed for 209/211 of them (primers: **Data S1**, list of CRE sequences and positions: **Data S3**), see section below, and 204/209 were sufficiently represented in our constructed scQer libraries to allow for quantification (**Fig. 4A**, **S7B-C**, **S8H**).

### 2.4. Construction of CRE series dual RNA reporter plasmid library

#### *2.4.1. Recloning of oBC-mBC backbone plasmid*

Doubly barcoded backbone plasmid (p025) was recloned in order to increase the complexity of the barcode pairs. Briefly, new barcodes were appended by amplifying the region between the oBC and mBC with primers with random (5'VNNVNNVNNVNN for the oBC) primers (oJBL513+oJBL514) with Kapa HiFi (15 cycles). Following 1.5x Ampure XP beads clean up, the barcoded insert was further amplified (15 cycles Kapa HiFi) to append homology arms for Gibson assembly (oJBL515+oJBL516). The final insert was PAGE purified. The insert-compatible backbone was reconstructed from two PCR products from p025 (oJBL524+oJBL527, oJBL526+oJBL525) of about 2.6 kb each (agarose gel purified). The three pieces were then combined by Gibson assembly, and electroporated in *E. coli* (C3020, NEB). Full complexity of the library was maintained, and estimated to be ≈1M clones by colony counting.

#### *2.4.2. oBC-mBC subassembly*

Following re-cloning of p025, the oBC-mBC pairs in the library were obtained as described for the promoter series experiment by a single step of PCR (primers oJBL337+oJBL345) to append handles for sequencing (on NextSeq 500, with library structure: read 1 oJBL346 (oBC, 30 cycles); index 1 oJBL347 (library index, 15 cycles); read 2 oJBL334 (mBC reverse complement, 18 cycles); index 2 oJBL348 (oBC reverse complement). Pre-processing also was carried out as described, resulting in oBC-mBC pairs with each associated with a read count. Given the complexity of the library (unsaturated), a cutoff of at least 5 reads was applied to retain oBC-mBC pairs (1.2M pairs). To mark possible non-uniquely paired barcodes, we computed the proportion of read counts to each oBC and mBC from a given oBC-mBC pair. oBC-mBC pairs with oBC or mBC with read counts proportion belonging to the pair of less than 95% were marked as likely non-unique (78.1% likely unique pairs by this criterion).

#### *2.4.3. PCR cloning of putative developmental CREs and assembly in dual RNA plasmid*

Putative CREs selected for profiling (see above) were cloned by PCR from mouse genomic DNA. A compromise amplicon size of 0.9 kb was taken as rough target size in order to balance testing large regions without overly compromising success rate. In order to increase specificity, a nested PCR approach was taken: a first unburdened PCR with selected primers (below), followed by a second nested PCR using primers with homology arm for cloning in the common backbone.

Outer primers for the first PCR (**Data S1**) were selected by running Primer-BLAST (*94*) with as PCR templates the 1200 bp sequences for the putative CREs (350 bp symmetric extension on both sides of the ArchR called 500 bp ATAC peak window with bedtools (*95*) slop, followed with bedtools getfasta to obtain sequences from mm10 genome) with the following run criteria: PCR product size 800 to 1000 bp (forward primer between 0 and 200 bp and reverse primer between 1000 and 1200 bp), primer melting temperature Min 57.0 Opt 60.0 Max 63.0, Max Tm difference 3, no intron junction preference, specificity check to *Mus musculus* (taxid:1009). For certain CREs, Primer-BLAST did not return any specific result with these constraints. Constraints (on product size) were then sequentially relaxed to increase the search space, with ultimately requiring only that the product be at least 500 bp within the windo. Five regions (Foxa2_chr2_13861, Sparc_chr11_7210, Lamb1_chr12_2182, Lamb1_chr12_2183, Sox17_chr1_58) were still too repetitive for Primer-BLAST to return results but had non-repeat sequences enabling manual primer selection. Two regions were too repetitive to find any primer pairs whatsoever and were thus not included in the screen (Sparc_chr11_7186, Foxa2_chr2_13842). Overall, primers were ordered to PCR clone 209/2011 CREs from our initial selected set.

Inner primers for the second nested PCR were selected using batch primer3 (*96*) (non default options: GC clamp=1, max poly-X=4) using the first PCR product as a template (but allowing for at most 8 bp overlap between inner primers and the PCR1 product). Primer pairs leading the largest nested PCR product were selected and handles homologous to the backbone were added (forward: 5'accgcatcgatctcgagg[inner forward], reverse: 5'tcccaaagcagatgtagttgac[inner reverse]). Handles were added to the forward/reverse primer so that the orientation of the CRE relative to the promoter matched their relative orientation on the genome relative to the gene.

The first PCR was performed in 20 uL reactions with 40 ng of genomic DNA (harvested from the mESC line used [DNeasy, Qiagen] following manufacturers' instructions) with Kapa Robust (Roche) with following parameters: 95C 3 min; 40 cycles: 95C 15 sec, 60C 20 sec, 72C 1 min 40 sec; final extension 72C 1 min 40 sec; with individual reactions in separate wells of a 96-well plate with primers distributed using a 96-liquidator (Rainin). Products were cleaned up (1x Ampure XP beads) and visually checked on agarose gel (with >95% success rate as judged by presence of ~1kb-sized band, possibly with non-specific products), and eluted in 100 uL of 10 mM Tris 8. 0.5 uL of the purified up PCR1 products was taken as template for the second nested PCR using the same conditions, but with the inner primers. The resulting products were cleaned up (0.6x Ampure XP beads) and visually checked on agarose gel, showing a <2% failure rate and highly clean products (little non-specific bands/smears). The products were quantified with a spectrophotometer (Nanodrop), and pooled to a 1:1 ratio by weight. This pool was used as insert for a pooled Gibson assembly as described below.

Prior to addition of the putative CRE PCR products, the minimal promoter GFP cassette (reporter mRNA) was inserted in the doubly barcoded backbone p025 digested with EcoRI and BglII (NEB) (**Fig. S7A**) and maintained at highest clonal complexity upon transformation (electroporation without bottleneck) to generate plasmid library p043. The minP-GFP insert was generated by splice PCR (templates: minP fragment: amplification of p027 with primers oJBL314+oJBL416; GFP fragment: amplification of p027 with primers oJBL254+oJBL414) followed by gel extraction. Plasmid library p043 was then digested with NheI/MfeI, combined with the pooled PCR amplified CREs via Gibson assembly, and transformed (electroporation) with a bottleneck via 100-fold dilution to an estimated complexity of ~50k clones (**Fig. S7A**). The resulting plasmid library (p055) was then subjected to the final subassembly step to connect oBC to the CRE.

*2.4.4. oBC-CRE subassembly*

Given the length of the inserted CREs (≈1 kb) and diversity of sequences, amplification of the region from minimal promoter to oBC was not a feasible strategy to subassemble oBC to CRE (≈1.3 kb from minP to oBC). We thus relied on tagmentation followed by semi-specific PCR. Briefly, plasmid library p055 was tagmented with Tn5 (Illumina, Nextera Tagment DNA enzyme, cat. no. 15027916) at a concentration such that the expected fragment size would be larger than the oBC to minP distance (≈1.3 kb), determined by a Tn5 titration curve experiment. Following tagmentation (5 uL 2x Tagmentation DNA buffer [Illumina, cat. no. 15027866], 0.4 uL Tn5 enzyme 1, 3.6 uL water, 1 uL 10 ng/uL plasmid library; 30 min at 37C), the tagmented plasmids were cleaned up (Zymo clean and concentrator, 3:1 binding buffer), eluted in 10 uL Tris 8 10 mM. 1 ng (1 uL of the elution) was amplified via semi-specific PCR with a Nextera primer with a P5 handle (oJBL512, binding to all P5 tagmentation events) and a oBC-specific upstream primer (oJBL502, binding to specific portion of the plasmid) in 25 uL (8.9 uL water, 12.5 uL 2x NEBNext master mix, 1.25 uL 10 uM oJBL502, 1.25 uL oJBL512, 1 uL tagmented plasmids, 0.1 uL 200x SYBR green) with the following conditions (gap fill: 72C for 5 min, 98C for 30 sec, then 12 cycles of 98C for 10 sec, 65C for 30 sec, 72C for 1 min). As controls for the non-specific product size distribution, the tagmented plasmids were also amplified with oJBL512 exclusively. Following purification (Zymo clean and concentrators), the amplified libraries were run on PAGE (6% TBE, 180V, 30 min). As anticipated, the amplicons with primers oJBL502+oJBL512 (semi-specific products) displayed reduced size distribution compared to oJBL512 alone amplified (non-specific) products, with most oJBL512 exclusive amplicons >1.2 kb. Semi-specific oJBL502+oJBL512 products between 450 bp and 800 bp were size selected on the PAGE gel, purified (minimum size size from CRE ≈ 75 bp), and sequenced (read 1: CRE sequence, Illumina Nextera primer [no custom], 34 cycles; index 1: P7-idx, primer oJBL432 15 cycles; read 2: oBC, primer oJBL433, 30 cycles).

Following demultiplexing (from the P7 index), the sequencing data was processed by first aligning read 1 (mapping to CRE) using bowtie2 (v2.4.4) (*97*) using option '-k 2' to report multi-mapping regions (some of our CRE segments overlapped given the proximity of the called peaks and extension from 500 bp to ≈1 kb tested regions). The resulting alignment sam file was then sorted, converted to bam using SAMtools (*98*), and merged with the oBC (read 2) using custom scripts into a file storing the oBC, CRE identity of the mapping, position and strand of aligned read within the CRE. Total read counts to each oBC-CRE pair were then summed up with custom scripts, retaining information about distribution of alignment positions and strand within the CRE for downstream processing.

The piled-up count data on oBC-CRE pairs was then filtered to identify *bona fide*, unique pairs. First, pairs with median mapping position outside the expected range from the size selection step (<30 bp and >300 bp) and mapping on the incorrect strand were filtered out. Then, the proportion of oBC reads mapping to any given CRE was calculated across oBC-CRE pairs, and only pairs with >95% of oBC reads mapping to a unique CRE were retained. The read count distribution across all oBC-CRE pairs was bimodal suggesting a saturated library, and only pairs with >30 reads (separating the two modes) were retained. Finally, pairs with anomalously small or large mapping position dispersal (90th to 10th percentile difference mapping positional spread <30 bp or >300 bp) were filtered out. We note that the positional filters enabled unambiguous discrimination between different but overlapping CREs (given that in all cases one of the CRE would have out of range mapping positions compared to the expected size from the amplicon library). Two elements (*Gata4*:chr14_5749, *Txndc12*:chr4_7975) shared a short identical sequence complicating the mapping, and were treated separately to not confound the fraction of oBC reads mapping to a given CRE. Following these filtering steps, we were left with 43.6k valid oBC-CRE pairs.

##### *2.4.5. Final oBC-CRE-mBC triplet table*

These oBC-CRE subassembled pairs were then linked with the previously determined oBC-mBC pairs from the starting plasmid library p025. Briefly, oBC (from final oBC-CRE pairs) were joined to mBC via valid oBC-mBC pairs (restricting to the uniquely mapped pairs). The resulting valid triplets oBC-CRE-mBC were then joined with the oBC-promoter-mBC triplets of the exogenous promoter library (experiment from **Fig. 2A**), and any oBC or mBC appearing twice in both libraries were removed from the final triplet list. The final number of valid oBC-CRE-mBC triplets was 33.0k, with a median of 145 valid mBC-oBC pairs per CRE. The resulting triplet map was used to deconvolute single-cell data in reporter quantification. Through the cloning and subassembly process, 5 out of the attempted 209 CREs dropped out (<20 valid mBC-oBC pairs), and consequently could not be quantified (*Col1a1*:chr11_15306, *Col1a2*:chr6_65, *Cited2*:chr10_1265, *Txndc12*:chr4_7952, *Btg1*:chr10_9570).

#### 2.5. Experimental details of pooled screen for CRE in mEBs

##### *2.5.1. Transfection, cell culture, and bottlenecking*

Low passage number mESCs were expanded in serum+LIF medium on gelatin coated plates as described above (passaged every two days, medium change every day) on 10 cm plates. Cells were transfected using Lipofectamine 2000 (Thermo Fisher Scientific) using reverse transfection. Briefly, cells washed with 1x PBS, and lifted by adding 2.5 mL/10 cm plate of trypsin 0.05% (Gibco). Following incubation at 37C for 5 min, cells were triturated with an added 7.5 mL of medium, spun down at 300g for 5 minute, and resuspended by pipetting at an estimated 1.5 M/mL to obtain a single-cell suspension. Following straining (40 um), cells were counted and diluted to 0.5 M/mL with medium. Concurrently, the lipofectamine+opti-MEM (12 uL lipofectamine + 238 uL opti-MEM) and the opti-MEM+DNA (240.4 uL optiMEM + 4 uL 50 ng/uL transposase + 5.6 uL transposon mix) were separately prepared and mixed by pipetting. The 500 uL lipofectamine+DNA+optiMEM mix was then added to a gelatin coated plate, 1 M cells (2 mL) from the single-cell suspension was added to the plate, and gently mixed. No transposase and no DNA controls were included. The transfected transposon was a uneven mix of three components (too boost MOI, see below): 1) 89% of the p055 oBC-CRE-minP-GFP-mBC library, 2) 10% of the oBC-promoter-puromycin-GFP-mBC series (same as for experiment in cell lines, **Fig. 2A**), and 3) 1% of the EEF1A1p-mCherry plasmid (p060, see below). Two biological replicates were transfected in parallel, one with the hyPBase plasmid (*46*), and one with super PiggyBac (SBI). We did not find substantial difference in MOI in the two replicates (**Fig. S8D**, replicate A vs. B).

Transfected cells were passaged and expanded to allow for integration and unintegrated plasmid dilution. Five days post transfection, cells were split with a portion selected on puromycin (2 ug/mL), and another portion remaining unselected. After 5 days on puromycin, cells from no DNA controls and no transposase controls were dead. While a large proportion of cells in samples with integrated cargos samples died, the puromycin resistant population was expanded for two weeks post transfection to ensure complete dilution of the unintegrated plasmids (maintained on puromycin).

The two replicates were induced to form mEBs in CA medium (no puromycin) on suspension plates as described above (day=0, 14 days post transfection), starting with 24 M cells per replicate (8 10 cm plates with 3 M cells each in 15 mL of CA medium). Replicate A was the sample transfected with hyPBase (and selected on puro), replicate B the sample transfected with the SBI super PiggyBac. Following induction, mEBs were passaged every two days, with sampling 5-10% of EBs at each time point for bulk MPRA (for harvesting, mEBs were pelleted at 5 min at 300 g, medium aspirated, fixed with ice cold 80% methanol, and stored at -80C until processing).

At the 12 day time point, a subset of expanded cells from replicate B were sorted by FACS for mCherry signal, and plated on a MEF monolayer (Thermo Fisher, CF1 Mouse Embryonic Fibroblasts, MitC-treated, cat. no. A34958, plated at 0.4M cells per well) in the wells of 6-well plate at approximately 1000 cells/well for bottlenecking. Following colony expansion for 4 days with daily medium change, colonies were lifted as follows: two washes with 1x PBS, add 750 uL collagenase type IV (0.1%, Stemcell Technologies, cat. no. 07909), 8 min incubation at 37C, lifted colonies aspirated by pipetting. The collagenase treated colonies on MEFs were then gently washed twice with 1 mL of serum+LIF medium added dropwise to recover additional colonies, and pooled with the previous ones. Lifted colonies were then spun down (400 g, 5 min), medium aspirated, trypsin treated to single-cell suspension (250 uL 0.05% trypsin used to mix the pellet, incubated 3 min at 37C, inactivated and triturated with 2 mL of fresh medium, and plated on gelatin-coated plates for expansion. Counting colonies suggested about half, or 500 clones, were obtained in this way. Following expansion for 8 days, mEB induction with 24 M cells (8 10 cm plates with 3 M cells each) was initiated as above. mEBs were passaged every two days, with sampling 5-10% of EBs at each time point for bulk MPRA as before. The bottlenecked replicate was termed 2B.

*2.5.2. End-point processing and single-cell sequencing*

For both non-bottlenecked and bottlenecked experiments above, mEBs were processed at the three weeks end point as follows (for each replicate): 2 suspension 10 cm plates of mEBs were pooled into a 50 mL conical left to settle. Medium was aspirated and mEBs were washed twice with 1x PBS, resuspended in 3 mL 1x PBS in the second wash, and split in two 1.5 mL aliquots in 2 mL tubes. PBS was aspirated from the tubes, and 500 uL of trypsin 0.25% was added per tube. Tubes were then agitated on a thermomixer at 37C and 650 rpm for 4 minutes. Cells were then gently dissociated by pipetting 10 times, and placed back on the thermomixer for 2 min. 1 mL of medium was then added per sample and pipetted to obtain a single-cell suspensions, the two samples were combined in a 15 mL conical, after passing them through a 100 um strainer. The strained single-cell suspension was counted, and cells were spun down (300 g, 5 min), resuspended to 4 M/mL, and taken to FACS. >600k cells were then FACS sorted (in <50 min) in pre-warmed medium to ensure the single-cell nature of the suspension (no gating on fluorescence, only on forward and side scatter) prior to generating the emulsions for single-cell RNA-seq. Sorted cells were then spun down at 400 g at 4C for 5 min, the medium gently aspirated, and cells resuspended to an expected 2.5 M cells/mL (based on FACS sort event counts) in ice cold 1x PBS + 0.04% BSA, cells were further counted and volume adjusted to 1200 k/uL with ice cold PBS+BSA.

Single-cell suspensions in PBS+BSA were taken as the starting point for the 10x Genomics protocol (v3.1 with feature barcoding). Emulsion and reverse transcription were performed per the manufacturer's instruction. Given prior empirical experience with mEBs processing, each 10x lane was slightly overloaded (by an additional 20%) to approach the expected recovery of 10k cells/lane. Each replicate was profiled with 2 lanes of 10x, for a total of 6 lanes.

*2.5.3. Construction of EEF1A1p-mCherry transposon plasmid*

To obtain an orthogonal selection for co-transfection to boost MOI (see below), we cloned a constitutively expressed red fluorescent protein into the piggyBac transposon. Briefly, p001 (piggyBac transposon backbone) was digested with XbaI and EcoRI (NEB), and size selected on agarose. The EEF1A1 promoter was amplified from p003 using primers oJBL536+oJBL537, and mCherry was amplified from a puro-mCherry containing plasmid with primers oJBL538+oJBL539 with Kapa Robust. The resulting fragments were size selected on agarose, and combined with the digested backbone by Gibson assembly, and transformed. The final plasmid taken from an individual colony was confirmed by Sanger sequencing and used for co-transfection.

*2.5.4. Optimization of high multiplicity of integration with piggyBac in mESCs*

As described above, we used co-transfection of selectable carrier transposon to boost multiplicity of integration following previous successful reports (*63*, *64*) of the procedure. Importantly, to prevent any bias on expression of the integrated reporters associated with developmental CREs, we leveraged orthogonal selection modalities not associated with the CREs (puromycin and red fluorescent protein, not green fluorescent protein). We directly confirmed the increase in MOI by comparing qPCR-estimated cargo DNA doses from genomic DNA of cells at 11 days post transfection with and without puromycin selection. The qPCR was performed as follows: gDNA extraction with DNeasy, dilution to 100 ng/uL, per well reaction with 5 uL 2x PowerUp master mix (Thermo Fisher, cat. no. A25741), 2 uL 10 mM Tris 8, 1 uL 5 uM forward+reverse primer pair, 1 uL 100 ng/uL gDNA. Primer pairs used: GFP: oJBL039+oJBL040, puromycin cassette: oJBL043+oJBL044, *Tfrc1* (endogenous locus for normalization): oJBL276+oJBL277. We observed on average a 4.0 to 7.0-fold increase in MOI (with vs. without puro selection, cargo dose with per-sample normalization from endogenous locus) across three biological replicates (with 10% co-transfection of puromycin containing cargo). We note that in our hands, other approaches used to optimize MOI (selecting on higher dose of puromycin, tuning relative and absolute concentration of transposon and transposase, selection on top 10% GFP intensity) did not improve MOI to the same extent as this co-transfection method. Notably, by adding a second selection round on mCherry positive cells on puro selected expanded cells (mCherry plasmid co-transfected at 1% of the cargo DNA), we saw a further increase in MOI in replicate 2B (median MOI ≈20 for replicates A and B, and up ≈50% to median ≈30 for replicate 2B, **Fig. S8D**) suggesting further optimization might be possible to increase median MOI beyond what has been achieved and boost power. We note that transfecting more cells might then be necessary to avoid extensive bottlenecking (already some detectable through clonal analysis, see below, even for the non explicitly bottlenecked populations replicates A and B).

2.6. Single-cell reporter libraries preparation and sequencing

The three single-cell libraries (gene expression GEx, oBC, mBC) for the mEB experiment were prepared as described for the benchmarking experiment in human cell lines (with no splitting of the cDNA, spike-in at 0.5 uM of primers oJBL246 and SR38 at first cDNA amplification to enrich for the reporter barcodes), with the following modifications.

oBC libraries were prepared the same way as for human cell line experiments, but with different P7-indexed primers for the final inner PCR (oJBL501-oJBL506). In addition, to avoid loop-the-loop products in the oBC libraries (anecdotally decreasing sequencing quality), the lowest band in the circularized ladder amplicons (see e.g., **Fig. S1B**, **S2D**) was size selected on PAGE for each library and used for sequencing.

Given the limited added value of CS2 capture for mBC (**Fig. S1F**), only the poly-dT captured libraries were generated for the mBC, with the following primers for the two rounds of PCRs (following initial cDNA amplification). PCR2: oJBL324+oJBL529. PCR3: oJBL076+ P7-indexed primers (oJBL530-oJBL533).

GEx, oBC, and mBC libraries were sequenced at the same time for replicates A/B on NextSeq500 (read 1: 28 cycles, no custom primers; index 1: 10 cycles, spike in primers oJBL432, oJBL534; read 2: 54 cycles, spike in primers oJBL433, oJBL334). The three libraries for replicate 2B were similarly sequenced, except with 8 cycles on index1 and 56 cycles on read 2. The oBC libraries for replicates A and B were re-sequenced as part of a NextSeq2000 run, with 28 cycles on read1, 10 cycles on index1, and 20 cycles on read2, with primers SR40+oJBL433 in well 1, and primer oJBL432 in well 2.

### 2.7. Single-cell reporter data processing

#### *2.7.1. Quality filtering from gene expression libraries*

Fastq files were generated using the makefastq command from cellranger (v6.0.1), and the gene expression count matrices were then generated with cellranger count, with transcriptome reference mm10-3.0.0. Raw count matrices were then imported as a Seurat (*86*) object (filtering genes expressed in less than 3 cells, and cell barcodes with less than 50 genes measured). Cell barcodes in the high total UMI mode with low mitochondrial RNA proportion were filtered as likely *bona fide* cells (fraction of mitochondrial UMI >1% and <15%, total gene expression UMI > 400 for samples from replicates A, B, and 2B lane1, and >1000 for 2B lane2, which was fortuitously sequenced more deeply). The filtered count matrices were then used to evaluate doublet scores using scrublet(*88*) (scrub_doublets command, 30 principal components, mean_center=true, normalize_variance=true), and cell barcodes with doublet score > 0.3 (separating the two modes of the simulated doublet distribution from scrublet) were filtered out. Datasets from all replicates were then combined in a single Seurat object, dimensionally reduced and clustered (NormalizeData, normalization.method= "LogNormalize", scale.factor=10000; FindVariableFeatures with selection.method = "vst", nfeatures=1000; ScaleData with all genes as features; RunPCA with identified variable features and 100 principal components; FindNeighbors, dims=1:50; FindClusters, resolution=0.2; RunUMAP, dims=1:50, n.neighbors=50) without batch correction given the good correspondence between replicates (**Fig. S8C**). The cluster identities were taken as categories for cell-type expression testing (see integration section below).

The following additional quality filtering steps were applied to retain high confidence singlet cells. Clusters comprising less than 1% of cells were considered likely doublets/artifacts, and corresponding cells were removed. Cells members of each cluster identified were separately sub-clustered with the same procedure as above (except resolution 0.5 in FindNeighbors). Any sub-cluster with a median doublet score above 0.15 was deemed composed of likely doublets, and corresponding cells were removed. Cells with anomalously high gene expression UMI counts were removed (with technical lane specific thresholds: >10k for A.1, >9k A.2, >12k B.1, >9k B.2, >15k 2B.2, no anomalous cells in 2B.1). Finally, cells with an estimated MOI > 200 (roughly corresponding to the top 0.1% of the distribution, MOI estimated through oBC UMI > 10, see below) were filtered out. In the end, n=43799 cells passed all these quality filters (12859 replicate A, 15422 replicate B, 15518 replicate 2B).

*2.7.2. mBC and oBC libraries*

Raw data was processed in the same way as for the human cell line promoter experiment to obtain a table of barcodes (mBC or oBC) with read and UMI counts per cell barcode. For oBC libraries of replicates A and B, two sequencing runs (for higher depth) were combined into one by concatenating their fastqs (trimming to the same read size) prior to running in cell ranger for the first processing step. Only cell barcodes passing the QC filters from the GEx analysis were retained in the final count tables.

*2.7.3. Single-cell quantification of reporter expression*

A similar approach as the promoter series experiment was taken to quantify expression in individual cells. For a given CRE of interest, all cells with associated oBC (in the list of valid oBC-CRE-mBC triplets) captured at >10 UMI counts were retained. The associated mBC UMI counts, in the respective cells, was then divided by the depth normalized GEx total UMI counts, and multiplied by the mean normalized GEx total UMI count across all cells (as before, to on average have a normalization factor with a mean of 1 to not systematically distort the scale of UMI counts while correcting for systematic factors such as cell sizes and overall efficiency of in-emulsion reverse transcription). To correct for slight differences in coverage between replicates, total GEx UMI count across cells was taken per replicate, and used to normalize (by direct division, under the valid assumption that the GEx libraries are far from saturation) the GEx UMI count in individual cells (different scaling factor per replicate). In cells in which multiple reporters (with different oBC-mBC pairs) corresponding to the same CRE were detected (via the oBC), the average across this normalized mBC UMI count was taken to obtain the per-cell estimate (for displaying the single-cell enhancer activity maps). For statistical tests and quantification, the average was taken across integration events as opposed to across cells (not first averaging internally within each cell, and then averaging over all cells, instead directly averaging over all integration events, such that each detection event carries the same weight).

*2.7.4. Quantification of activity and specificity of CREs and statistical tests*

The following stringent tests were performed to identify active and specific CREs. Each CRE and biological replicate was considered separately.

To assess activity, all integration events (oBC UMI > 10) for the CRE considered were identified, and the total number of such integration events for the CRE recorded. $10^4$ bootstrap resamplings (random sampling with replacement) of the integration events were then performed. In parallel, sampling with replacement of integration events (same number sampled as the CRE considered to control for difference in representation) from both basal promoter controls (minimal and no promoters). For each bootstrap sampling, the average normalized mBC UMI counts (see above), stratified by cell-type clusters (Seurat identified, see **Fig. S5A**), were determined both for the CRE and the basal promoters. The maximum expression cluster identity and expression level in that cluster was stored. Mean expression of the reporter without stratification by cluster identity was also obtained (over all bootstrap resampled integration events irrespective of cell types). Following bootstrapping, an empirical p-value was determined as follows: the null distribution was taken as the maximum cluster expressions across all bootstrap samplings of the two basal promoters. The empirical p-value of expression for the CRE considered to have activity in excess of the basal control (activity p-value) was taken as the probability that maximum cluster bootstrap CRE expression was below that of the basal controls, averaged over all bootstrap sampling for the basal control events (effectively corresponding to a rank-sum test). Empirical activity p-values (over all CREs within a replicate) were Benjamin-Hochberg corrected to obtain a false discovery rate. Corrected empirical p-value without stratification over clusters was similarly performed (mean probability that expression from the CRE over all integration was below that of basal control null bootstrap values). To identify active CREs, we considered elements with either per-cluster maximum expression FDR <10% in all three replicates and/or all cells expression FDR<1% (higher statistical power from more integration events) in all three replicates. 58/204 CREs passed these stringent criteria and were considered active in excess of our basal expression controls.

To assess CRE specificity, a similar approach was taken, but instead of performing comparison to basal promoters, comparisons were performed to datasets with permuted cell cluster identities. For each CRE, $10^4$ repeats were performed were a bootstrapped resampled (no cluster identity permutation) set of integration events was generated, and the fold-change in reporter expression (average normalized mBC UMI) between the maximum expression cluster and the rest of cells was computed. The corresponding quantity, but for a cluster-identity permuted sampling was also performed for each sampling. The specificity empirical p-value for each CRE was taken as the average (over resamplings) probability that the cluster permuted fold-changes in expression (null distribution over all permutations) was higher than the non-permuted one. As before, these empirical p-values were Benjamin-Hochberg corrected (over all CREs, separately for different biological replicates). CREs that were identified as active were further marked as specific if in all biological replicates, the reporter expression fold-change (maximum cluster vs. all other cells) was >5 and the permutation derived FDR < 10%, leading to 9/58 elements.

To systematically assess whether elements had pleiotropic activity (active in multiple cell types), we computed the fold-change in expression in all pairs of clusters vs. the rest of cells, storing the maximum fold-change value and specific cluster pair for each CRE and biological replicate. The median (across biological replicates) fold-changes for pairs vs. individual clusters were compared. Only a single CRE had a paired/single cluster fold-change in excess of 3x was *Lamc1*:chr1_12189 (also elevated: 2.6x for *Foxa2*:chr2_13858 which displayed some activity in visceral endoderm in addition to parietal, **Fig 4E** second row; and 1.5x for *Sox2*:chr3_2007 which had some activity in epiblast cells, **Fig. 3F**). Other elements showed no substantial excess activity in pairs over single clusters (95% percentile in pair/single fold-changes was at 1.3x and 90% percentile at 1.1x). Permutation tests similar to above confirmed Lamc1 bifunctional activity was highly significant (non permuted fold-change highest in all $10^3$ samplings), leading to a final set of 10/58 active CREs labeled as specific.

To summarize the function of individual CREs, the median activity (defined as the maximum cluster mean reporter expression) and specificity (defined as the fold-change between maximum cluster mean reporter expression vs. mean reporter expression in the rest of cells) across the three biological replicates was determined (shown in **Fig. 4A**).

Some elements were active and/or specific in only a subset of replicates (those marked in **Fig. S10B**, e.g., *Bend5*:chr4_8174, *Foxa2*:chr2_13820, *Sox17*:chr1_77, *Bend5*:chr4_8179, *Lama1*:chr17_7791, *Lamc1*:chr1_12185). These are likely candidates for active elements (falling below our limit of detection possibly because too few integration events were captured due to uneven CRE representation), but were not retained to maintain stringency in our downstream analyses. Quantification summary can be found in **Data S5**.

Pseudobulk expression in separate cell-types (e.g., **Fig. S9B**, **S13**) were determined as the average normalized mBC UMI counts over all cells with detected reporters belonging to GEx clusters identified and annotated in **Fig. S5A**.

### 2.8. Bulk MPRA (CREs, mEB time series experiment)

#### *2.8.1. Library preparation and sequencing*

Bulk MPRA libraries for the CRE time series were generated similarly as described for the human cell lines promoter series experiment (**Fig. 2A**) with the following modifications. Genomic DNA and RNA were extracted with the AllPrep kit (Qiagen). 40 samples (different replicate/batch/time points) were processed overall, comprising 13 samples for replicates A and B across 12 time point (day 0, 4, 6, 10, 12, 14, 16, 18, 20, 21; with two technical replicates for day 16) and 14 samples for rep2B (two technical replicates for day 0, 8, 12, 16, 18, 20; one replicate each for day 4, 6). From each sample, 2 libraries (1 gDNA-derived, 1 RNA derived) were constructed, for a total of 80 libraries. Libraries were prepared in three batches (batch1: replicates A and B, days 0, 4, 8, 12, 16; batch2: replicates A and B, days 2, 6, 10, 14, 16, 18, 20, 21; batch3: all samples from replicate 2B). For RNA, DNase treatment was applied to the first batch, but was found to be unnecessary (comparing to no reverse transcription controls), and was consequently not performed on the other two batches. As before, reverse transcription used primer oJBL358. The first PCR using the cDNA for the RNA-derived libraries was with primers oJBL077+oJBL039. The first PCR from genomic DNA was with primers oJBL039+oJBL358. The second PCR (performed on both RNA and gDNA derived samples) using primer oJBL077 and a set of indexed primers (oJBL359-oJBL366, oJBL437-oJBL448, oJBL555-oJBL564).

Each preparation batch was sequenced separately. Batch 1: Nextseq500; read 1: 28 cycles, primer oJBL369 (mBC forward); index 1: 8 cycles, primer oJBL435 (UMI); read 2: 43 cycles, primer oJBL371 (mBC reverse); index 2: 6 cycles, primer oJBL370 (P5-index). Batch 2: Nextseq2000, same set of primers (well 1: oJBL369+oJBL370, well2: oJBL370+oJBL435), read 1: 28 cycles, index 1: 10 cycles, read 2: 20 cycles, index 2: 6 cycles. Batch 3: Nextseq500: read 1: 18 cycles, index 1: 10 cycles, read 2: 20 cycles, index 2: 6 cycles.

*2.8.2. Data processing and quantification*

Data was pre-processed in the same way as for the human cell line bulk MPRA. Briefly, following demultiplexing with bcl2fastq (v2.20), mBC reads were trimmed to their expected lengths (15 nt) with seqtk's trimfq. mBC reads were then joined/error-corrected using PEAR (v0.9.11, options -v 15 -m 15 -t 15). The correctly assembled barcodes were then reformatted and merged with the (pseudo-)UMI read using custom python scripts, resulting in a list of mBC-UMI paired reads. The read counts for each mBC-UMI pair was determined, and a final pileup performed to generate a table of total UMI and read counts for each mBC.

From these raw mBC UMI counts, only mBC sequences from valid oBC-CRE-mBC triplets (including the promoter series) from our subassembly (34121 mBC total, 33001 from CREs, and 1120 from exogenous promoters) were retained, and appropriate metadata information (sample, time point, RNA/DNA, etc) was appended. DNA UMI counts for each barcode (across each sample) were then normalized for sequencing depth dividing by the summed UMI counts from that sample. RNA UMI counts were similarly normalized. To obtain an activity for each CRE, we first only included well-represented mBC (requiring >20 DNA reads) for quantification. Then, we 1% winsorized DNA and RNA normalized UMI counts (to mitigate extreme outliers) across all barcodes from a given CRE. The winsorized normalized UMI counts were then summed across mBCs (for a given CRE) for DNA and RNA, and the ratio was taken to be the activity of the CRE in that sample. For a given CRE, the averaged activity from all samples from two adjacent time points (days 0 & 2, 4 & 6, 8 & 10, 12 & 14, 16 & 18, 20 & 21) were shown in **Fig. S14**, with error bar the standard deviation of the mean across these samples.

As a statistical test of activity, we used a Wilcoxon rank-sum test (one-sided). At each aggregate time point, the activity from all samples from the CRE of interest was compared to the activity of basal expression controls (minimal and no promoter) from all samples/time points. The resulting p-values (across all time points and CREs) were Benjamin-Hochberg corrected. Activity displayed as significant when the false discovery rate was below 1% (**Fig. S9C**, **S14C**, summary of quantification in **Data S6**). The fold-change in activity over time (**Fig. S14B**) was taken as the mean activity for day 20.5 (all samples from days 20 and 21) over day 1 (all samples from day 0 and 2).

### 2.9. Single-cell data integration

#### *2.9.1. Integration between scRNA-seq and Pijuan-Sala et al in vivo scRNA-seq*

We compared our day 21 mEB (containing scQers) scRNA-seq data to available *in vivo* data from mouse development (*65*) (E6.5 to E8.5) to annotate identified clusters from low dimensional projections of our data. Samples spanning time points E6.5 to E8.5 were obtained (using the R library 'MouseGastrulationData', function EmbryoAtlasData with all samples except ids 11, 22, and 23). The count matrix was extracted together with the metadata, and a Seurat object was created after converting the gene names for compatibility, and merged with the mEB dataset. We performed integration as previously described (*99*). Briefly, a list of objects was generated from the merged Seurat object, the two datasets were separately normalized and features identified (NormalizeData; FindVariableFeatures, selection.method="vst", nfeatures=2000). Functions SelectIntegrationFeatures, FindIntegrationAnchors, and IntegrateData were sequentially applied to the list, and the integrated data was then dimensionally reduced via scaling and PCA (ScaleData, RunPCA with 30 principal components). The PCA embedding space from the integrated dataset was used to identify neighbors using a method adapted from (*100*). For each cell in the mEB dataset, the top 10 closest distance neighbors in the dataset-integrated PCA space from the *in vivo* dataset were identified, and their cell-type annotation stored. Cell annotation from the *in vivo* data was transferred if >6/10 nearest neighbors had the same cell-type label, and taken as 'uncertain' otherwise. This provided an annotation label for each cell in our mEB dataset. To aggregate the annotation across clusters in the mEB data, we determined the fraction of cells per mEB derived clusters with *in vivo* cell-type annotation, shown in the heatmap of **Fig. S5C**. In that representation, *in vivo* cell types with a maximum fraction across all mEB clusters <5% were not displayed for brevity. Final mEB cluster annotations were determined by inspection, and coarse-grained clusters (**Fig. 3C**) naturally combined cell-types from the same lineage. One important distinction was the label of pluripotent cells, not present in the *in vivo* dataset given that the earliest time point covered was E6.5. The putative cluster of pluripotent cells was closest to epiblast cells within this constrained label-transfer assignment (**Fig. S5C**), but inspection of key marker genes of naive pluripotency (*66*, *101*) such as *Esrrb*, *Dppa3* (**Fig. S5B**), *Zfp42*, and *Tdh* (not shown) were sharply expressed in that cluster, in contrast to markers of primed pluripotency (*Fgf5*, *Dnmt3b*) which were expressed in other clusters (**Fig. S5B**). These justified our identification of this cluster as pluripotent cells.

Performing the label transfer on coarse-grained annotations (grouping all endodermal cells, ectodermal cells, etc.) decreased the proportion of the 'uncertain' label, which in some instances was spuriously created by mEB cells associated with mixed populations from otherwise well-defined lineages (e.g., the multiple different mesodermal cell types). We verified that the final label transfer was robust to the number of neighbors (5 to 20) considered in the PCA integrated embeddings and to another integration method (Harmony (*102*), not shown).

#### *2.9.2. Integration between scRNA-seq and scATAC-seq and correlation with in vivo data*

The scRNA-seq and scATAC-seq in mEBs was not performed on the same set of cells or as a co-assay (but samples were derived from the same mESC line). We therefore relied on computational approaches to relate the clusters of the low dimensional representations from the two modalities. To that end, we performed unconstrained integration using ArchR (*92*) function addGeneIntegrationMatrix which uses the functionalities of Seurat (*99*). The resulting assignments unambiguously mapped clusters from the RNA to the ATAC (**Fig. S5D**), with some of the finer resolution achievable in the scRNA-seq (e.g., different mesodermal and neuroectodermal clusters) not distinguishable in the scATAC possibly as a result of the fewer number of nuclei sampled from these cell types.

As additional verification for the validity of these cell-type assignments on the scATAC data, we compared the data to available scATAC datasets from mouse embryos at E7.5 and E8.5 (*57*). We downloaded pileup bigWig scATAC files from all cell types (GEO: accession GSE205117), and generated bigWig pileup from our mEB datasets (ArchR's getGroupBW function, tileSize=50, maxCells=100000, ceiling=10, normMethod="ReadsInTSS"). We then computed the average accessibility across all peaks called by ArchR using UCSC utility function (*103*) bigWigAverageOverBed. Restricting to the top 25% scoring peaks called in the mEB scATAC dataset (ArchR score >20, corresponding to 65k peaks), when then computed the $R^2$ on log-transformed peak accessibility in the *in vivo* and mEB datasets across all cell-types/clusters. The overwhelming majority clusters in the mEB scATAC data assigned from the comparison to scRNA-seq had their highest correlations to corresponding *in vivo* cell types: mEB parietal endoderm vs. *in vivo* parietal endoderm $R^2$=0.77; mEB mesoderm vs. *in vivo* mesenchyme $R^2$=0.76, vs. Pharyngeal_mesoderm $R^2$=0.72, vs. Paraxial mesoderm $R^2$=0.72; mEB neuroectoderm vs. *in vivo* Forebrain, Midbrain, Hindbrain $R^2$=0.78, spinal cord $R^2$=0.75; mEB pluripotent/epiblast vs. *in vivo* epiblast $R^2$=0.86; mEB visceral endoderm vs. *in vivo* ExE endoderm $R^2$=0.61, vs. visceral endoderm $R^2$=0.48. Only mEB surface ectoderm had a higher correlation with another in vivo cell-type (highest *in vivo* correlation to gut, $R^2$=0.67, we note that the label-transfer from scRNA-seq datasets suggests partial recognition of surface ectoderm as gut, **Fig. S5C**), but still had accessibility highly correlated to the expected cognate cluster (second highest correlation $R^2$=0.60 to *in vivo* surface ectoderm). Taken together, these show the mEBs harbor complex epigenetic states broadly representative of *in vivo* gene regulation.

### 2.10. Clonal cell analysis

#### *2.10.1. Clonotype identification, refinement, cell assignments, basic metrics, and dropout assessment*

Analysis proceeded as described above for the human cell line experiment with minor modifications. First, only oBC associated with CREs (not exogenous promoters) were considered for clonal assignment. That was to minimize the likelihood of spurious doublets being called as a result of the lower complexity of the promoter library (increasing the likelihood of co-integration of the same pair of barcodes in otherwise unrelated clones). The UMI cutoff for oBC detection per cell was set to >10. Other parameters for the procedure (raw clonotype identification with Fisher exact test, clonotype refinement, cell assignment to high confidence clonotype) were as before.

Across the three replicates, the fraction of cells assigned to high confidence clonotype was 4535/12859 (29%, 896 clonotypes) for replicate A, 6854/15422 (53%, 866 clonotypes) for replicate B, and 8406/15518 (54%, 360 clonotypes) for replicate 2B. The mean numbers of cells assigned for these high confidence clonotypes were respectively 5.0, 7.9, and 23.4 for replicates A, B, and 2B, consistent with replicate 2B having been directly bottlenecked. We note that evidence of substantial clonal expansion in the non explicitly bottlenecked replicates (A and B) suggests that our procedure to select high MOI cells (selection on puromycin from ≈5% of plasmid transfected containing the expressed resistance cassette) did severely reduce the complexity of the cell population. While replicate 2B corresponded to a sub-sampling of replicate B (we were not aware at the time of substantial bottlenecking in our population), most clones and cells did not overlap between the two samples (44 clonotypes identified in both samples, or 44/866 of clonotypes comprising 499/6854 clonotype-assigned cells for replicate B; 44/360 of clonotypes 2533/8406 clonotype-assigned cells for replicate 2B). Summary of clonotypes and assigned cells can be found in **Data S8**.

oBC dropout analysis (**Fig. S8I-K**) from the clonotypes and cell assignment was performed as described for the human cell line experiment.

#### *2.10.2. CRE expression pattern across clones*

From the assignment of cells to high confidence clonotypes, we sought to characterize how the expression of CREs varied across clones, with the assumption that different clones correspond to different genomic positions of integration of the reporter driven by the CRE. For each of the 10 active cell-type specific CREs identified, we obtained the list of high confidence clones harboring at least one reporter integration corresponding to the CRE. In order to obtain sufficient statistical power to estimate expression, we then restricted the analysis to clones with 5 or more assigned cells in both the cell type of expected CRE expression (e.g., pluripotent for *Sox2*:chr3_2007, parietal endoderm for *Gata4*:chr13_5729, etc.) from the analysis over all cells, and 5 or more cells assigned to the rest of cell-types. We then computed the fold-change in mean reporter expression (average normalized mBC UMI) over cells in these two compartments (cognate vs. rest of cells), and calculated the number of clones per CRE for which fold-change was >5. For 9/10 CRE, more than ⅔ of clones retained a >5 specificity (**Fig. S12**).

2.11. Analysis of features of profiled putative developmental CREs

Various features of the profiled CRE were considered for correlation with cell-type specific activity. These were determined as follows:

ATAC accessibility: for each peak in each cell, the corresponding read count was normalized by the total number of TSS reads in that nucleus. The average overall cells assigned to a given cluster was then taken as the mean accessibility for the given peak in that cluster. Fold-change in accessibility was taken as that measure of accessibility over the mean accessibility averaging over cells from well-delineated clusters (pluripotent/epiblast, neuroectoderm, mesoderm, and extraembryonic endoderm/parietal), such that for example fold-change accessibility for parietal endoderm was: mean accessibility in parietal endoderm divided by mean accessibility in all cells from epiblast/pluripotent, neuroectoderm, and mesoderm clusters. The visceral and intermediate parietal endoderm clusters were not considered for the fold change computation to not have cells from the same lineage be included in the comparison, which could have artificially decreased the effect size for parietal endoderm.

Pseudotime opening: In the absence of a time series scATAC-seq dataset, we considered pseudo-time trajectories (*70*). First, scATAC data was clustered at higher resolution (resolution=2) using ArchR's addClusters function. A trajectory from pluripotent to parietal endoderm, passing through these more highly resolved clusters, was then defined and created with the addTrajectory function. Accessibility information along the trajectory was extracted with function getTrajectory (useMatrix="PeakMatrix", log2Norm=TRUE, and smoothWindow=10). Pseudotime smoothed accessibility values for each considered peak (distal parietal endoderm) was then obtained. To estimate the pseudotime at which a peak became accessible, we fit an exponential sigmoid (logistic function, using SSlogis and nls in R) to each accessibility vs. pseudotime trace. The pseudotime at which the sigmoid reached 20% of its maximum from baseline was selected as the heuristic value to compare opening times of the different peaks.

Evolutionary conservation: to assess evolutionary conservation, we calculated the average phyloP (*69*) score (mm10.60way.phyloP60way.bw) over ArchR-defined 500 bp ATAC peaks using function bigWigAverageOverBed (*103*) similarly to previous assessment of non-coding element conservation (*11*).

Transcription factor binding sites: To characterize the transcription factor binding composition of tested elements using a biophysically grounded empirical approach (in the absence of high resolution ChIP-seq data in our system), we took an approach inspired by Farley and colleagues (*3*, *29*). Briefly, we downloaded protein array binding data from key endodermal transcription factors Gata4, Foxa2, and Sox17 from Uniprobe (*73*–*75*), which provides affinity measures for all 8-mers. We first converted the raw measurements ("Median" column in the raw data files) to relative affinities. To do so, we treated the mode value of affinities as the experimental noise floor, and computed the relative affinity as: (affinity-baseline)(max(affinity)-baseline). We note that the final list contains a relative affinity for an 8-mer and its reverse complement (as the protein binding arrays hold double stranded DNA). Before computing the maximum affinity, we divided the score of palindromic 8-mers by two, as we found those to be anomalously high. The resulting relative affinity 8-mer table was then used to scan all regulatory elements and genomic regions, yielding a value for each 8-bp stretch (in a strand agnostic manner). We then identified local maxima in the relative affinity trace. For a given relative affinity threshold, the local maxima above threshold were retained. To collapse maxima close to each other, we generated a graph between maxima (one node per maximum) with an adjacency matrix determined by distance (connect maxima less than 3 bp apart). Connected components of the graphs were identified, with one TFBS assigned per connected component (typically a single maximum) at the highest-affinity position. The procedure was applied across a wide range of affinity thresholds, and across full genomic loci (±100 kb from TSS, 500 bp windows with 250 bp sliding step and excluding tested CREs and surrounding 500 bp, **Fig. S15C**).

For single-feature classifiers, a simple thresholding on the feature was used to generate the ROC curves (**Fig. S15B**). To combine cognate ATAC accessibility and number of Gata4 binding sites, we used scikit-learn (*104*) function LogisticRegression with an l1 penalty (mean=0 and standard deviation=1 input variables) and the roc_curve function to compute the performance metric.

### 3. Pol III driven circular vs. linear barcode MPRA experiment

#### 3.1. Cloning of plasmids

The Tornado cassette was first cloned in a piggyBac transposon. The piggyBac cloning dock p022 was digested with BbsI (NEB) and the U6-Tornado-Broccoli insert excised from the pAV-U6+27-Tornado-Broccoli plasmid (*37*) (Addgene #124360) with BamHI and XhoI (NEB) digestion. Both backbone and insert were purified by agarose gel extraction (Zymoclean Gel DNA recovery kit, Zymo Research), the fragments combined by isothermal assembly (HiFi NEBuilder, NEB) into plasmid p051, and transformed in *E. coli* (NEB, C3040H). A single clone was selected and the plasmid confirmed by Sanger sequencing. A truncated version of the Tornado cassette, excluding the 5' and 3' portion of the ribozyme not overlapping with the final circular RNA sequence, was generated by digesting p051 with XbaI and SalI, and combining with gblock linear_TB_CS by isothermal assembly. The resulting plasmid, p052, was transformed in *E. coli* (NEB, C3040H). A single clone was selected and the plasmid confirmed by Sanger sequencing.

To generate complex libraries barcodes, barcoded inserts (5'VNNNVNNNVNNNVNNN) with downstream capture sequence 1 (CS1, 5'GCTTTAAGGCCGGTCCTAGCAA) were amplified from ultramer uJBL519. For circular barcodes, primers oJBL520+oJBL521 were used for amplification, the resulting product PAGE purified, and inserted in NotI+SacII digested p051 purified by agarose gel extraction by isothermal assembly. The plasmid library, p053 (**Fig. S2B**), was concentrated and eluted in water (Zymo Clean and Concentrator, Zymo research), and electroporated in *E. coli* (NEB, C3020) following manufacturer's instruction. A similar procedure was taken for linear barcodes, except that primers oJBL522+oJBL523 were used to amplify uJBL519, and integrated in NotI+SacII digested p052, resulting in plasmid library p054 (**Fig. S2C**). Of note, both circular and linear barcode constructs were compatible for reverse transcription and amplification from the same primers for library preparation to minimize biases. Following outgrowth post electroporation, a dilution series was plated to assess library complexity, and populations estimated at 50k clones were expanded, and resulting plasmids libraries purified (ZymoPure II plasmid Midiprep kit, Zymo Research), and further concentrated to 1 µg/µL by isopropanol precipitation.

#### 3.2. Transfection, cell culture, and cell harvesting

2.75 µL each of plasmids libraries p053 and p054 were mixed to 0.6 µL (0.3 µg) of SBI super piggyBac, and transfected 2.5 M of exponentially growing K562 cells in duplicates using a Nucleofector following manufacturer's protocol for K562 (kit V4XC-2024, Lonza BioResearch) in duplicates. After two weeks of exponential growth with ⅕ split every two days to allow for dilution of unintegrated plasmids, cells were harvested in exponential phase (<0.75 M/mL), and methanol fixed. Briefly, cells were pelleted at 500 g for 5 min, washed with ice cold 1x PBS to 2 M/mL, pelleted at 500 g for 5 min, resuspended to 15 M/mL, and ice cold methanol was added drop by drop to 80%. Aliquots of 4 M fixed cells were stored at -80C until DNA or RNA extractions.

3.3. Massively parallel reporter assay library generation and sequencing:

Bulk MPRA for Pol III barcodes proceeded similarly as described before. Fixed cells were split, and genomic DNA was extracted from methanol fixed cells using the DNeasy kit (Qiagen), and RNA was extracted from cells using TRIzol LS (Thermo Fisher), following manufacturer's instructions in both cases.

Amplicon libraries from DNA were generated in two steps of PCR amplification with Kapa HiFi (Roche). For genomic DNA, 500 ng of input was used, and for plasmids (to map barcodes present in both constructs), 3 ng was used. For low-cycle number PCR1, 500 ng of DNA was mixed with 50 µL 2✕ Kapa HiFi master mix, 5 µL 10 µM oJBL246, 5 µL 10 µM oJBL424, and water to 100 µL. Cycling parameters: 1 min at 95C, and 4 cycles of: 20 s at 98C, 20 s at 60C, 30 s at 72C, followed by 4C hold. Primer oJBL424 contains 10 random Ns to serve as a pseudo-UMI (hereafter referred to as UMIs for brevity) to correct for PCR jackpotting. Reactions were cleaned up with Ampure XP beads (Beckman Coulter) at 1.75✕, and eluted in 20 µL of 10 mM Tris 8. Illumina adapters and sequencing indices were appended through PCR2, with 4 µL of the eluate from PCR1 taken as input, and 25 µL 2✕ Kapa HiFi master mix, 0.25 µL 100✕ SYBr green, 2.5 µL 10 µM oJBL076, 2.5 µL 10 µM indexed primers (DNA rep1: oJBL501, DNA rep2: oJBL502, plasmid p053: oJBL427, plasmid p054: oJBL504), and water to 50 µL. Libraries were amplified with tracking by qPCR with: 1 min at 95C, and cycles up to the qPCR inflection point (typically 15-17 cycles) of: 20 s at 98C, 20 s at 60C, 30 s at 72C. Libraries were then cleaned up with Ampure XP beads at 1.75✕.

Amplicons libraries for RNA were obtained by first DNase treating the RNA (5 µg RNA, 2 µL TURBO DNase [Thermo Fisher], 2 µL 10✕ buffer, and waster to 20 µL, incubated at 37C for 30 min, cleaned up with RNA clean & concentrator [Zymo Research], and eluted in 11 Tris 7 10 mM), and taking 1 µg of DNase treated RNA to reverse transcription. Briefly, 2 µL (500 ng/µL) RNA was mixed with 2 µL 1 µM oJBL424, incubated at 65C for 5 min, and placed on ice. 15 µL of reverse transcription master mix was then added (4 µL 5✕ FS buffer, 1 µL 0.1 M DTT, 1 µL 10 mM dNTP mix, 8 µL water, 1 µL SSIII [Thermo Fisher]), and the reaction incubated at 55C for 60 min, followed by 70C for 15 min. ¼ of the reverse transcription reaction was then directly amplified for PCR1 (37.5 2✕ Kapa HiFi master mix, 3.75 µL oJBL246, 3.75 µL oJBL076, water to 75 µL), with cycling parameters: 1 min at 95C, and 4 cycles of: 20 s at 98C, 20 s at 60C, 30 s at 72C, followed by 4C hold. Reactions were cleaned up with Ampure XP beads (Beckman Coulter) at 1.75✕, and eluted in 20 µL of 10 mM Tris 8. PCR2 proceeded as for libraries prepared from plasmids and genomic DNA, with indexing primers oJBL508 and oJBL509 for replicates 1 and 2 respectively, and reactions stopped at inflexion point from qPCR tracking (cycle 7). Libraries were then cleaned up with Ampure XP beads at 1.75✕. Notably, given the circular nature of the Tornado barcodes, rolling-circle loop-the-loop RT products were prominently visible (at least 4-loops products detectable) at the expected size laddering from repeats of the circular RNA length (**Fig. S2D**). To prevent possible phasing issues on the sequencer, the product of the lowest size, which was the same for both linear and circular barcodes, was purified by PAGE extraction.

Final amplicon libraries were quantified with Qubit dsDNA HS (Thermo Fisher), diluted to 3 nM, run on TapeStation D1000 HS (Agilent) for final quality assessment, and adjusted to final 2 nM based on the Tapestation quantification. Libraries were pooled, loaded as a fraction of a NextSeq500 lane, and paired end sequenced with the following parameters: read1 (barcode forward): 25 cycles with primer oJBL431, index1 (index): 20 cycles with primer oJBL432, read2 (barcode reverse): 20 cycles with primer oJBL433, index2 (UMI): 10 cycles with primer oJBL434.

3.4. Data pre-processing and quantification

Sequencing data was demultiplexed using bcl2fastq. Raw fastq files were processed first by trimming unnecessary cycles from the 3' end (9 cycles from read 1, 4 cycles from read 2) using seqtk (https://github.com/lh3/seqtk). Forward and reverse barcode reads were joined and error corrected with PEAR(*85*) (options -v 16 -m 16 -n 16 -t 16). Using custom python and R scripts, successfully assembled barcode reads were combined with UMI reads, barcode/UMI pairs were counted, and the read counts and UMI count per barcode was determined. These barcode count files served as the processed inputs for downstream analysis.

For downstream processing, the identity of barcodes present in each transfected plasmid library (p053 and p054) was first determined by inspecting the distribution of barcode UMI counts from separate libraries directly prepared from the respective plasmids. The count distribution displayed clear bimodal nature, and barcodes in the high count mode (>9 UMIs for p053, >5 UMIs for p054) were retained as valid. Following removal of barcodes present in both libraries (34 out of 193705), we were left with a list of barcodes for expression analysis (59.0k for p053, 134.6k for p054).

For quantifying steady-state expression of linear and circular barcodes, we tallied the UMI counts for all valid barcodes for genomic DNA and RNA derived libraries. DNA UMI counts were reasonably correlated from genomic DNA to plasmid ($R^2$ of log-transformed BC UMI counts= 0.42, and 0.43 respectively for replicate 1 and 2). Steady-state expression (referred to as "activity") was defined as the normalized RNA UMI counts over the normalized DNA UMI counts, with normalized UMI counts defined as UMI counts over all UMIs mapping to valid barcodes in the respective libraries. For BC well represented in the library (>50 DNA UMI counts), the activity was >150-fold higher for Tornado barcodes compared to linear barcodes (**Fig. S2E**, median activity fold-change 162✕ in replicate 1 and 186✕ in replicate 2). Difference in activity was largely insensitive to threshold selection on DNA UMI, and the summed RNA/DNA UMI counts across all linear vs. circular barcodes irrespective of DNA UMI counts confirmed >100-fold higher in activity for circular over linear barcodes (107✕ for rep1, 113✕ for rep2). Circular barcodes had a tight range in activity across barcode sequences (interquartile range in activity spanning 2.5-fold and 2.6-fold for replicates 1 and 2 respectively).

3.5. Estimating expression levels of oBC per cell per integrated cassette

To estimate the relative steady-state expression level of oBC driven by human U6 Pol III promoters, we used two different reverse-transcription qPCR quantifications, from K562 cells with genome-integrated dual reporter constructs. First, following cell harvesting and RNA extraction as previously described, 1 µg of DNase-treated RNA was combined with 100 pmole of random hexamer in 2 µL of 10 mM Tris 7 buffer, incubated at 65C for 5 min, and placed back on ice. 8 µL of MuLV mix (1 µL 10✕ buffer, 0.5 µL 10 mM dNTP mix, 6 µL DEPC treated water, 0.5 µL MuLV [NEB]) was added to the RNA and random hexamer mix, and incubated at 25C for 5 min, 42C for 60 min, and 65C for 20 min. RNA was hydrolyzed from the reverse transcription mix by adding 2 µL of 1M NaOH and heating to 95C for 5 min. The cDNA was subsequently neutralized by adding 2 µL of 1M HCl, and diluted ten-fold by adding 86 µL of 10 mM Tris 8. For each primer pair, 2 µL of diluted cDNA was directly used for qPCR, and run by adding 2 µL of 10 mM Tris 8 and 5 µL of PowerUp master mix (ThermoFisher) and 1 µL forward+reverse 5 µM primer mix per well. Each primer pair/sample was run in technical triplicate wells, with PCR conditions (2 min at 50C, 2 min at 95C, and cycles: 15s at 95C, 15s at 60C,, 15s at 72C). qPCR primers targeting both the reporters (Pol III oBC: oJBL246+oJBL247, Pol II GFP mRNA: oJBL039+oJBL040), and highly expressed endogenous genes EEF1A1 (oJBL001+oJBL002, with these primers obtained from (*105*)) were used. oBC expression was normalized to EEF1A1 level using a ΔCt method. To normalize for multiplicity of integration in the genome, we performed qPCR from extracted genomic DNA extracted with DNeasy (Qiagen), using 100 ng gDNA input per triplicate (5 µL PowerUp SYBr mix, 1 µL forward+reverse 5 µM primer mix, and 10 mM Tris 8 to 10 µL), using the same cycling parameters as for RT-qPCR, and primers targeting the piggyBac reporter payload (GFP: oJBL039+oJBL040, puromycin cassette: oJBL043+oJBL044) in addition to endogenous genes for normalization (RPPH1: oJBL085+oJBL086, TERT: oJBL091+oJBL092). Relative levels of oBC RNA and Pol II GFP mRNA compared to the endogenous EEF1A1 mRNA were normalized by DNA dose per cell inferred from qPCR, leading to 5.2±0.8 for oBC and 0.12±0.04 for GFP (± standard error of the mean from 4 biological replicates), as the estimated expression per integrated copy expression. GFP level was the average produced by the five promoters included in the exogenous library (no promoter, minimal promoter, UBCp, Pgk1p, EEF1A1p), most of the expression coming from EEF1A1 promoter, and indeed close to the expected level of the endogenous EEF1A1 mRNA level when correcting for this factor (5×0.12=0.6≈1).

As an additional measurement, which might be not affected by possible systematic underestimation given the fact that oBCs are short (134 bp), leaving fewer space for priming from random hexamers, we used the qPCR cycle number obtained from preparation of sequencing libraries, which involves reverse transcription from target specific primers instead of random hexamers. For oBC, the same approach described for the linear vs. circular barcode MPRA was taken. For the mBC (the Pol II reporter mRNA), we used the same procedure, except with the following primers: reverse transcription with primer oJBL358, PCR1 with primers oJBL077+oJBL039, PCR2 with primers oJBL077 and one of oJBL359-oJBL366. Comparing the qPCR Ct value for oBC vs. mBC libraries, we estimated a ΔCt of 10.4±1.2 (± spread across two biological replicates) corresponding to a relative abundance fold-change of ≈1300 for oBC vs. mBC (we note that different RT or PCR primer efficiency could drive part of this difference), which is internally controlled for multiplicity of integration as both are part of the reporter construct. Correcting for the difference between the endogenous EEF1A1 mRNA and GFP reporter (per integrated copy) seen with random hexamers led to an estimate of 1300×0.12=156-fold higher oBC expression compared to the EEF1A1 mRNA, which is one of the most highly expressed mRNA in K562 cells (as assessed from the average of stranded bulk RNA-seq datasets from ENCODE (*87*) in K562). Given the presence of multiple rolling-circle reverse transcription products (**Fig. S2D**), we note that this quantification can be considered a slight overestimate. Taking the geometric mean of the random hexamer and target specific primer as an estimate of oBC abundance leads to ≈32-fold higher expression of oBC relative to the EEF1A1 mRNA. Given that EEF1A1 comprises 1.2% of mRNAs in K562 (estimated from bulk RNA-seq TPM), and taking 200,000 total mRNAs per cell (BNID109916 (*106*)), this leads to an estimate of 1.2%×200,000×32 >75,000 oBC RNAs per cell per integrated copy of the cassette in the genome, which converted to concentration assuming a radius of 10 um for K562 cells, leads to ≈30 µM.

#### 3.6. Evidence of Pol III hU6 transcription not enhancing for Pol II activity

To assess whether the presence of an upstream Pol III co-directional promoter could enhance Pol II activity, we compared expression of the minimal vs. EEF1A1 promoters with and without the oBC cassette in K562 cells (the constructs without Pol III promoters had a slightly different design compared to scQers: a barcode in their 5' UTR instead of 3' UTR and had GFP only mRNA, but otherwise were driven by the same promoters). The fold-change in activity from bulk MPRA (in the context of scQers, i.e., with Pol III cassette) between EEF1A1 and minimal promoter was 1430±300 (aggregating mBC quantification from **Fig. 2E** and **S3D**: sum 1% winsorized normalized RNA UMI over sum 1% winsorized normalized DNA UMI, error: range across the two replicates). Integrating the reporters without Pol III cassettes (separate transfections for minimal promoter: plasmid p002, EEF1A1 promoter: plasmid p003) via piggyBac as previously described, and harvesting cells >15 days post transfection (to allow for plasmid dilution), we then quantified DNA dose by qPCR (two primer pairs to GFP [oJBL039+oJBL040, oJBL041+oJBL042], normalized to endogenous genes (RPPH1: oJBL085+oJBL086, TERT: oJBL091+oJBL092) and RNA levels by RT-qPCR (two primer pairs to GFP [oJBL039+oJBL040, oJBL041+oJBL042], normalized to endogenous EEF1A1 [oJBL001+oJBL002]) as described above. The ratio RNA levels (reporter DNA-dose normalized) between the EEF1A1 promoter to minimal promoter was 1030±360 using this modality (standard deviation of mean across 4 comparisons from 2 biological replicates per promoter). While the experimental approaches used differ, these data suggest (since the fold-change in expression level minimal to EEF1A1 promoter is similar), that a Pol III promoter does not grossly enhance Pol II activity, at least from the minimal promoter (under the assumption of already saturated EEF1A1 promoter activity).
